# Non-genetic intra-tumor heterogeneity is a major predictor of phenotypic heterogeneity and ongoing evolutionary dynamics in lung tumors

**DOI:** 10.1101/698845

**Authors:** Anchal Sharma, Elise Merritt, Xiaoju Hu, Angelique Cruz, Chuan Jiang, Halle Sarkodie, Zhan Zhou, Jyoti Malhotra, Gregory M Riedlinger, Subhajyoti De

## Abstract

Impacts of genetic and non-genetic intra-tumor heterogeneity (ITH) on tumor phenotypes and evolvability remain debated. We analyzed ITH in lung squamous cell carcinoma (LUSC) at the levels of genome, transcriptome, tumor-immune interactions, and histopathological characteristics by multi-region profiling and using single-cell sequencing data. Overall, in LUSC genomic heterogeneity alone was a weak indicator of intra-tumor non-genetic heterogeneity at immune and transcriptomic levels that impacted multiple cancer-related pathways including those related to proliferation and inflammation, which in turn contributed to intra-tumor regional differences in histopathology and subtype classification. Genome, transcriptome, and immune-level heterogeneity influenced different aspects of tumor evolution. Tumor subclones had substantial differences in proliferation score, suggestive of non-neutral clonal dynamics. Scores for proliferation and other cancer-related pathways also showed intra-tumor regional differences, sometimes even within the same subclones. Neo-epitope burden negatively correlated with immune infiltration, indicating potential immune-mediated purifying selection on acquired mutations in these tumors. Taken together, our observations suggest that non-genetic heterogeneity is a major determinant of heterogeneity in histopathological characteristics and impacts evolutionary dynamics in lung cancer.

---

Despite growing from a single, renegade somatic cell, by the time of detection, a human lung tumor typically comprises of billions of cells that show considerable genetic and non-genetic differences among them, a phenomenon known as intra-tumor heterogeneity (ITH). Genetic and non-genetic ITH appear to be hallmarks of nearly all types of malignancies, providing substrate for evolvability and emergence of drug resistance, complicating treatment strategies for patients, and leading to unpredictable prognosis (Brock et al., 2009; Marusyk et al., 2012; Mcgranahan and Swanton, 2017). Indeed, certain patterns of genetic heterogeneity, as measured by the number of clones and presence of subclonal genetic alterations, predict poor survival in multiple cancer types including subtypes of lung cancer (Andor et al., 2016; Jamal-Hanjani et al., 2017). Likewise certain signatures of transcriptomic and immune heterogeneity appear to change with tumor subtyping, differentiation hierarchy, and response to treatment (Dalerba et al., 2011; Karaayvaz et al., 2018; Patel et al., 2014; Thorsson et al., 2018). Also, recent studies (Jia et al., 2018; Rosenthal et al., 2019) have suggested that immune editing influence both lung cancer evolution and survival, pointing towards complex interplay between tumor and microenvironment. But it is not well understood whether much of genetic and non-genetic intra-tumor heterogeneity correlate, and impact phenotypic variations and tumor evolution (De and Ganesan, 2017; Jia et al., 2018; Karaayvaz et al., 2018; Li et al., 2016; Patel et al., 2014; Suda et al., 2018; Zhang et al., 2018). In particular, there is a major gap in understanding the inter-relation between genetic, transcriptomic and immunogenic heterogeneity and their contribution in tumor evolution and variability in clinically relevant phenotypic characteristics in lung squamous cell carcinomas. In this study, we analyzed ITH in lung squamous cell carcinoma (LUSC) at the levels of genome, transcriptome, tumor-immune interactions, and histopathological characteristics by multi-region profiling and using single-cell sequencing data using a computational pipeline called metaITH.

## RESULTS

Here, we investigated patterns of intra-tumor spatial heterogeneity in 9 surgically resected stage I-IIIB lung squamous cell carcinoma specimens at genomic, transcriptomic, tumor-immune cell interactions, and histological characteristics by multi-region profiling (**Figure 1A**), and supplemented that with analysis of single cell sequencing data from additional 5 samples, as described later. For each tumor, 3-6 geographically distant regions were profiled. For each patient, tumor-proximal pathologically normal, benign lung tissue was also analyzed to serve as control. In total, 42 biopsies (10 normal and 32 tumor) were processed in this study. Clinical and pathological attributes of the samples are provided in **Supplementary Table 1A**, and representative H&E staining images are shown in **Supplementary Figure 1**. Pathological tumor purity for most regions was 40-80% (**Supplementary Figure 2**). For 29 biopsies representing 6 patient samples, DNA and RNA were co-extracted and analyzed jointly, while for the 13 biopsies representing 3 patient samples only DNA was extracted to investigate genetic heterogeneity.

**Figure 1.**
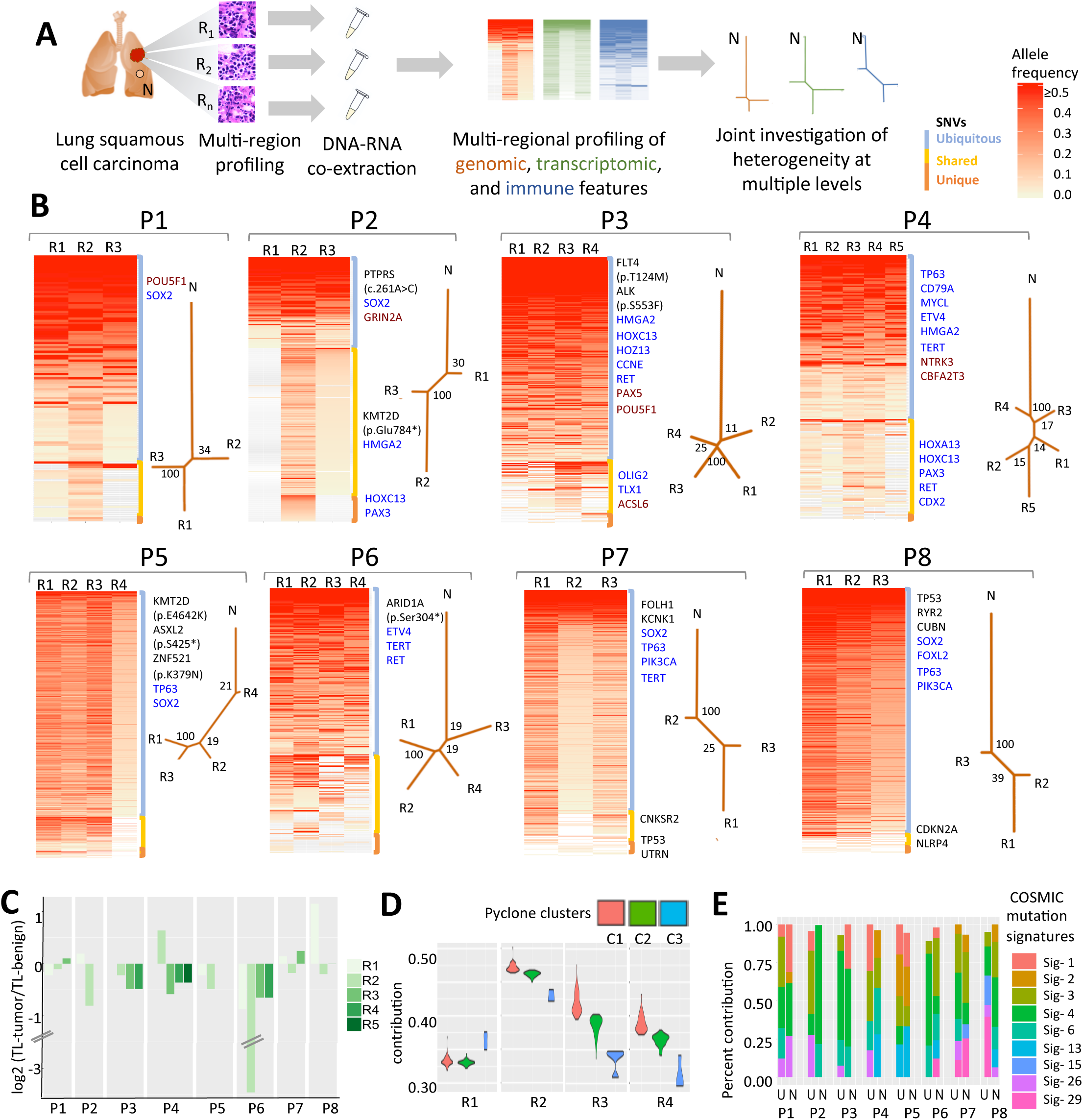
Assessment of genetic heterogeneity in lung squamous cell cancer. **A)** Schematic representation of the study design showing multi-dimensional analysis of intra-tumor heterogeneity based on multi-region profiling of tumor specimens. **B)** Heatmap shows regional variation in allele frequency of somatic single nucleotide variants (SNVs) for 8 patient samples. Somatic variants below detection threshold (<2% allele frequency) are marked in grey. Variations are categorized into three categories; Ubiquitous - present in all regions (blue), shared – present in multiple regions but not all (yellow), unique – present in single region (orange). Dendrograms represent genetic similarity (represented by branch length) between different regions within a tumor for each patient sample based on variant allele frequency of somatic SNVs. Numbers on the nodes represents bootstrap values. **C)** Regional differences in inferred telomere length in tumor regions, relative to matched normal tissue. **D)** Relative abundance of somatic mutation clusters identified in different tumor regions for P6. **E)** COSMIC mutational signatures inferred from somatic base changes in tumors corresponding to the mutations that are ubiquitous (U) and non-ubiquitous (NU-i.e. shared and unique). Signature.1: Mutational process initiated by spontaneous deamination of 5-methyl cytosine, Signature.2: Mutational process due to APOBEC activity, Signature.3: Mutational process due to HR defect, Signature.4: Mutational process due to smoking, Signature.6: Mutational process due to defective DNA mismatch repair, Signature.13: Mutational process due to APOBEC activity, Signature.15: Mutational process due to defective DNA mismatch repair, Signature.26: Mutational process due to defective DNA mismatch repair, Signature.29: Mutational process due to tobacco chewing habit.

### Landscape of genomic changes in the tumor samples

In the tumor specimens, using multi-region sequencing we identified 120-1606 somatic exonic SNVs, which included missense, nonsense and silent variants (**Figure 1B; Supplementary Table 2A**). The frequency of detectable somatic mutation (2/Mb-26/Mb) was comparable to that reported elsewhere (Alexandrov et al., 2013) (**Supplementary Figure 3**). A majority of the somatic SNVs (34-91%) were ‘ubiquitous’ across all regions in a tumor, while 2-55% SNVs were ‘shared’ among multiple tumor regions, and relatively small proportion of the somatic SNVs (0.8-10%) was ‘unique’ to a single region (**Figure 1B; Supplementary Figure 4**). We also inferred genome-wide copy number alterations (CNA) based on exome sequencing data (**Supplementary Table 2B, Supplementary Figure 5**). In general, the proportion of CNAs that were ubiquitous was smaller than the corresponding proportion of SNVs. This is in line with observations that as much as half of the somatic mutations in tumor genomes likely arise in benign progenitor cell prior to tumor development (and thus are ubiquitous) (Tomasetti et al., 2013), while copy number alterations in pre-neoplastic tissues are rare (Aghili et al., 2014; De, 2011). Occasional genome doubling in non-small cell lung cancer has been reported (Jamal-Hanjani et al., 2017), but we found no evidence for genome doubling in our samples based on allele-specific copy number profiles. Cancer cells survive cellular crisis through telomere maintenance mechanisms, so we also estimated telomere length in tumor regions relative to matched normal regions using TelSeq (Ding et al., 2014). Inferred telomere length varied across different regions within tumors, but overall, a majority of the tumor regions had shorter inferred telomere length than matched benign, pathologically normal regions (**Figure 1C**).

Among known cancer genes, *PTPRS, KMT2D, FT4, ALK, ASXL2, ZNF521* and *ARID1A* had ubiquitous somatic SNVs in P2, P3, P5 and P6 that were deemed potentially pathogenic (**Supplementary Table 2A**). Known oncogenes such as *SOX2, HOXC13, PAX3, TERT, TP63* were amplified in multiple samples, while tumor suppressor genes such as *POU5F1, NTRK3, GRIN2A* were deleted in either all or some regions in different tumor samples (**Supplementary Table 2B**). Copy number amplification was slightly more common than deletions, and impacted known cancer genes proportionally more (**Supplementary Figure 6**). A majority of the detected known oncogenic point mutations and copy number alterations were ubiquitous and therefore arose reasonably early during tumor development. But there are exceptions, e.g. in P2 potentially pathogenic mutation in *KMT2D,* a histone methyl-transferase implicated in non-small cell lung cancer (Ardeshir-Larijani et al., 2018), was only found in regions R2 and R3 but not in R1 region. Similarly, sample P3 harbors a potentially deleterious mutation in *CASP9*, a tumor suppressor, only in region R3. Copy number estimates from exome sequencing data tends to be less reliable, but in this case, inferred CNA estimates were consistent with gene expression changes of known cancer-related genes. For instance, copy number amplification of *SOX2* resulted in higher *SOX2* expression in tumors relative to matched benign tissue (**Supplementary Figure 7**).

To determine if mutation detection was biased by depth of coverage of sequencing, we also performed deep targeted sequencing covering 257 cancer-related genes from multiple regions from two samples (P8 and P9). Therefore, P8 had a total of 6 geographic regions profiled, 3 using exome sequencing, and 3 regions with targeted deep sequencing. This analysis did not change ubiquitous/shared/unique assignment of the existing catalog of somatic mutations in this sample, and no additional known oncogenic drivers were identified (**Supplementary Table 3**). Ubiquitous mutations (present in all three regions) in *FOXL2* gene in exome data were also identified in deep sequencing data (**Supplementary Figure 8A, B**).

### Patterns of genetic heterogeneity

To estimate the extent of inter-region genetic heterogeneity, we constructed dendrograms (**Figure 1B, Supplementary Table 4**; see **Methods**) for every patient based on variant allele frequencies for somatic SNVs across different regions, such that branch-lengths in the dendrograms indicate the extent of genetic divergence amongst the tumor and benign regions in a patient. The dendrograms were bootstrapped and corrected for tumor purity. The purity-corrected dendrograms were not substantially different from uncorrected ones (**Supplementary Figure 8C**). Overall, intra-tumor regional genetic heterogeneity was less compared to tumor-paired normal tissue differences. Patient samples P1, P3, P4 and P6 showed conservative patterns of genetic ITH, whereas patient samples P2, P5, P7 and P8 showed relatively high levels of ITH with subset of tumor regions harboring either substantially less tumor mutation burden (R1 of P2) or excess of low frequency variations as compared to other regions (R4 of P5 and R2 of P7). When targeted deep sequencing data was taken into consideration, the key conclusions about the tree topology in P8 was unchanged (**Supplementary Figure 8B and Supplementary Table 5**).

We also identified somatic mutation clusters using PyClone (Roth et al., 2014) based on somatic SNV and allele-specific CNA from multi-region tumor genomic data. Somatic mutation clusters may be considered as proxies for subclonal entities, but it remains a computationally challenging problem and caution is warranted while inferring clonal makeups from mutation clusters. Anyhow, typically 2-4 clonal clusters were identified for each tumor, and a majority of the clusters were present in all or most regions in a tumor, but their proportions varied between different regions (**Supplementary Figure 9**). For instance, in P6, we identified 3 prominent mutation clusters; all 3 were present in all regions in the tumor, but at varying proportions (**Figure 1D**). In addition, we adopted an orthogonal approach, and applied the method proposed by Williams et al. (Williams et al., 2018) to assess clonal architecture and compare that with our estimates. Overall, the clonal architecture the tumors are similar to that inferred by PyClone, with 1-2 subclones per sample (**Supplementary Figure 10**). In general, our analysis showed LUSC tumors to have low genetic heterogeneity, similar to that reported elsewhere (Jamal-Hanjani et al., 2017).

Anyhow, we further computed Shannon’s and Simpson’s indices for each region to quantify intra-region species richness and diversity in terms of abundance of different mutation clusters (**Supplementary Figure 11**); Shannon’s and Simpson’s indices were comparable across regions within a tumor, partly because a majority of the major mutation clusters were detected in all regions. All in all, mutation clusters and dendrograms suggested modest level of genetic ITH in the samples. Genomic ITH in LUSC is known to be lower than that observed in some other non-small cell lung cancer subtypes (McGranahan and Swanton, 2017).

Somatic mutation signatures provide insights into cancer etiology, exposure and patterns of disease progression. We identified signatures for smoking, consistent with the etiology of this disease (Jamal-Hanjani et al., 2017), as well as signatures of defective DNA mismatch repair, APOBEC and 5-methylcytosine deamination (**Figure 1E, Supplementary Figure 12**). APOBEC mutation signature was proportionally higher in non-ubiquitous mutations, which is consistent with reports that this signature probably arises late during tumor development (Jamal-Hanjani et al., 2017). Overall, genetic heterogeneity manifested by regional differences in oncogenic mutations, genomic alterations, telomere length, and mutation signatures, was moderate. As we show in a later section, an integrative analysis of ITH, mutation signatures, and transcriptomic profiles provide insights into different selection patterns and evolutionary dynamics in these tumors not apparent at the genome level data alone.

### Landscape of transcriptomic heterogeneity in the tumor samples

In the tumor and matched benign, pathologically normal tissues ∼18K genes had detectable expression based on 150bp paired end sequencing. First, an unsupervised principal component analysis of gene expression data across all samples showed that (i) benign tissues cluster together, indicating low level of between-sample variation as expected in non-diseased tissue specimens, and (ii) different regions from the same tumors typically cluster together, suggesting that intra-tumor transcriptomic variations were still relatively smaller than inter-individual differences (**Figure 2A**). For instance, different regions from within one tumor clustered together for P3, P4 and P6, but for P1, P2 and P5 some tumor regions showed greater variation. In case of P5, region 4 (P5-R4) clustered separately from rest of the three regions, which is consistent with the extent of genetic divergence this region had relative to benign tissue and other regions in P5. We note that P1-R1, P1-R3 and P2-R1 were relatively closer to the benign samples – which might be due to low tumor cell fraction in these regions (**Supplementary Figure 2**).

**Figure 2.**
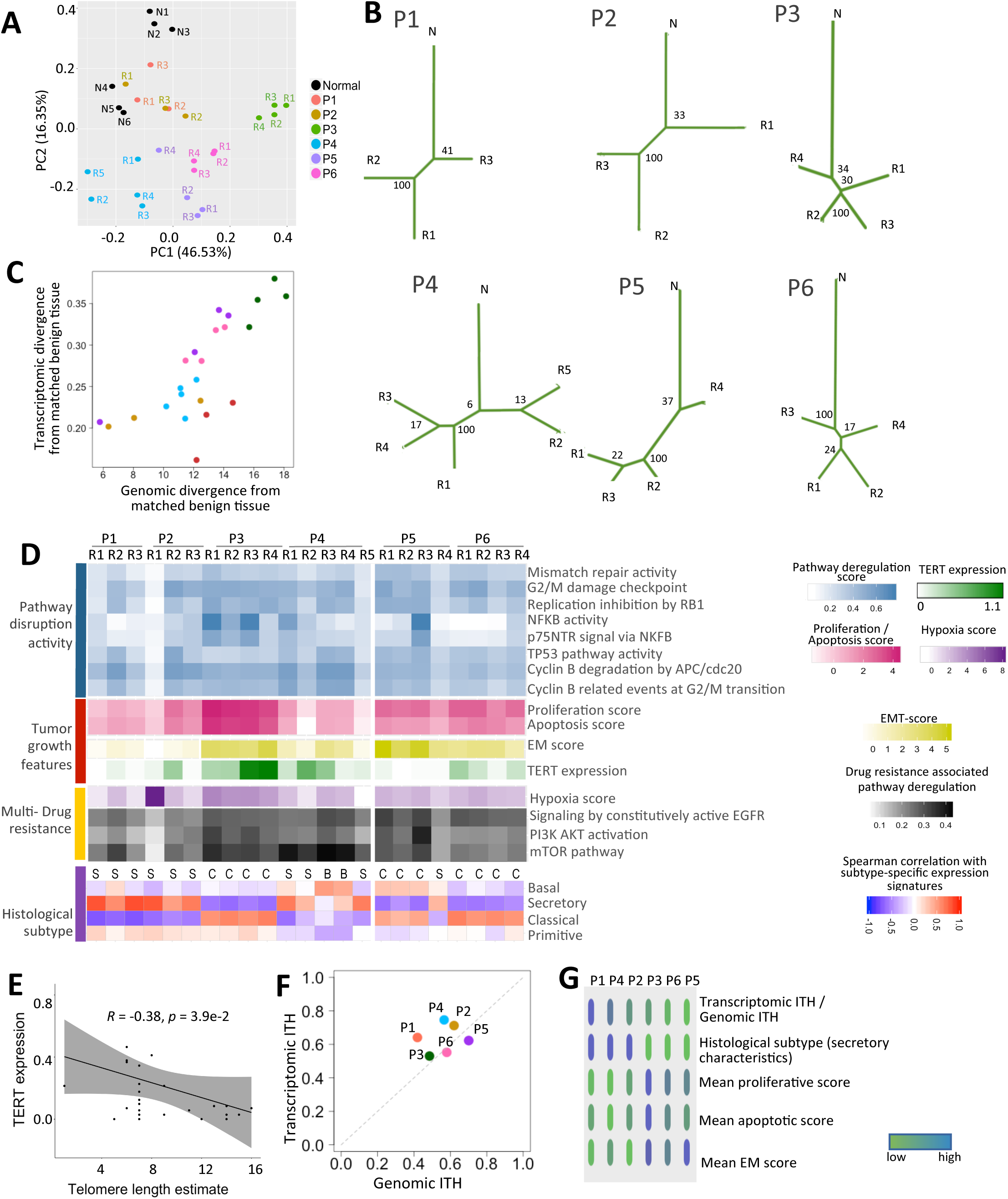
Assessment of intra-tumor transcriptomic heterogeneity in lung squamous cell cancer. **A)** PCA plot showing the extent of transcriptomic variation within and across tumor and benign tissue regions from the patients. Apparently normal, benign tissues from different patients are shown in black, while tumor tissues from different patients are shown in other colors. The proportion of variation explained by the first and second principal components are 46.53% and 16.35%, respectively. **B)** Dendrograms represent similarity (represented by branch length) between different regions for each patient sample based on gene expression profiles for all genes. Numbers on the nodes represent bootstrap values. **C)** Scatterplot comparing the extent of transcriptomic and genomic divergence for different tumor regions from their respective matched benign tissues. Color codes are same as Figure 2A. **D)** Heatmaps showing the extent of pathway disruption activity measured using nJSD, proliferative score, apoptosis score, epithelial-mesenchymal score, TERT expression, hypoxia score, multi-drug resistant pathway disruption activity and histological subtype score based on expression of published biomarker genes for different tumor regions. **E)** Scatterplot showing inverse association between TERT expression and telomere length estimates. Spearman correlation coefficient and p-value are shown at the top. **F)** Scatterplot comparing the extent of transcriptomic and genomic intra-tumor heterogeneity for the tumor samples. The color code is same as in Figure 2A. **G)** Oncoprint plot showing histological subtype (secretory characteristics), proliferative score, apoptotic score, and epithelial v/s mesenchymal characteristics score for the tumor samples ranked according to their ratio of transcriptomic ITH over genetic ITH.

We computed pairwise transcriptomic distance between regions, and accordingly constructed bootstrapped dendrograms for each patient using multiregional RNA expression data (see **Methods** for details; **Figure 2B, Supplementary Table 6**) to study the extent of transcriptomic divergence and ITH. Intra-tumor transcriptomic heterogeneity was generally lower compared to tumor-paired benign tissue transcriptomic divergence. Some tumors had overall lower transcriptomic heterogeneity compared to others. For instance, P3 and P6 had lower transcriptomic ITH compared to P4 or P5. We also compared the extent of divergence of tumor regions from paired benign regions both at genomic and transcriptomic levels, and found that those are highly similar (**Figure 2C**). For instance, in Sample P2, the region R2 showed both increased genetic and transcriptomic divergence from other tumor regions. The correlation between genomic and transcriptomic levels is not entirely due to regional variation in tumor purity, because (i) the correlation persisted even when adjusted for estimated tumor cell fraction in the cohort (Spearman partial correlation coefficient: 0.58, p-value: 4.36e-03), (ii) the regions with low tumor cell fractions did not necessarily have transcriptome profiles similar to the benign tissue, because in certain cases they had higher immune cell infiltration instead (**Supplementary Figure 13**), and (iii) such similarity was not apparent when we investigated ITH at other levels, as discussed later. Taken together, similarity in the general topology of the dendrograms and the extent of divergence from the matched benign, pathologically normal tissue both at the genomic and transcriptomic levels in a majority of the samples suggested that the patterns of heterogeneity at these two levels followed reasonably parallel patterns. This is in line with reported convergent patterns of genetic and epigenetic evolution and heterogeneity in some other cancer types (Mazor et al., 2015).

### Impact of transcriptomic ITH on cancer-related processes

Genetic changes involving cancer genes impacted transcriptome of associated pathways in a consistent manner. Oncogenes such as *SOX2, ETV4, CCNE1* and *TP63* had >=4 fold increased expression in multiple tumors, consistently across all regions therein, as compared to benign tissues (**Supplementary Table 7**), while tumor suppressor genes like *ERBB4* and *HNF1A* had low expression consistently in all regions in P4 and P6, consistent with their copy number status. Some other cancer genes such as *RET, PAX3, PAX5, PAX7* and *WT1* had altered expression only in some tumor regions. Multiple HOX family genes (*HOXA11, HOXA13, HOXC13*and *HOXD11*) were deregulated in multiple tumors; for instance, *HOXC13* showed subclonal duplication in samples P4 and P5 and consistent alterations in gene expression. Apobec-family genes (e.g. *APOBEC3B*) had high expression in some tumors such as P4 and P5, and indeed those tumors had APOBEC-related mutation signatures in somatic SNVs (**Figure 1E**). Likewise, altered mismatch repair activity was consistent with higher mutation burden and mismatch repair mutation signatures in P5.

We measured dysregulation in key cancer-related pathways (**Figure 2D, Supplementary Figure 14, Supplementary Table 8**) using a network based Jensen-Shannon Divergence (nJSD) score (Park et al., 2016), with values ranging from 0 to 1, where high values represents increased perturbation compared to matched normal tissue. NFKB activation pathway, which is associated with cell survival, was found to be highly deregulated in certain regions of tumors P3-R1 and P3-R3, P4-R1 and P5-R3. Immune pathways and steroid biosynthesis pathways were among the most highly deregulated pathways, with pathway entropy scores as high as 0.98. On the other hand, pathways like G2M DNA damage checkpoint, APC/CDC20 mediated degradation of Cyclin B and hedgehog pathways were moderately deregulated across all regions of all samples, except P2-R1 region.

Gene expression changes of certain genes also correlated with relevant genomic characteristics in a consistent manner. *TERT* is not expressed in benign tissues, but it was detected at varying levels in the tumors. Telomere length had negative correlation with *TERT* expression (Spearman correlation coefficient r=-0.32, p-value = 3.9e-02; **Figure 2E**) across the tumor regions, which is consistent with reports that *TERT* expression is negatively regulated by telomere position effect—over long distances (TPE-OLD) mechanism (Kim et al., 2016).

Altered oncogenic pathway activities are expected to impact clinically relevant growth characteristics of tumors e.g. rate of proliferation and apoptosis. We determined proliferation index (PI) for all the samples using gene expression data for 131 genes most associated with *PCNA* expression (see **Methods** for details). For every patient, PI for all tumor regions were higher than adjacent benign, pathologically normal tissues. Some tumors had regional differences in PI scores, e.g. P2-R2, P4-R3, P4-5 and P5-R1-3 showed high PI score as compared to other regions in the same tumors (**Figure 2D, Supplementary Table 9**). We note that these regions clustered away from other regions in the same tumors (**Figure 2A**) in transcriptomic profile and showed different genomic mutational patterns (**Figure 1B**). We also calculated apoptotic index (AI) in a similar manner based on known apoptosis-related genes. For samples P2, P4 and P5 some regions showed lower AI as compared to other regions (**Figure 2D, Supplementary Table 9**). Interestingly, in P4 and P5, AI also showed similar patterns of variability as PI. In general, AI and PI correlated across tumor regions (Spearman correlation coefficient: 0.51), which is consistent with TCGA data, where *PCNA*, *CCNB1* and *Caspase 7* expression correlate significantly in the non-small cell lung cancer cohorts (**Supplementary Figure 15**). This is probably due to the fact that replication stress is a major driver of apoptosis in rapidly proliferating tumors (Macheret and Halazonetis, 2015).

Phenotypic characteristics of tumor cells provide insights into tumor aggressiveness and histological classification. We computed epithelial-messenchymal (EM) characteristic score (see Methods for details) to determine whether some regions are more mesenchymal-like than others. P4 and P5 samples also showed heterogeneity in EM scores across different regions (**Figure 2D, Supplementary Table 10**) with P4-R3 and P4-R4 depicting more of epithelial signature as compared to the other three regions. Similarly P5-R1 and P5-R3 have higher EM score as compared to other regions. Intra-tumor heterogeneity can contribute to heterogeneous response/resistance to drugs, leading to therapeutic failure. For instance, overexpression *EGFR* is a common oncogenic event in lung cancers which leads to activation of tumor associated signaling cascades such as PI3K/AKT, MAPK and mTOR pathways. We examined the extent of deregulation of cellular pathways and gene-sets known to be associated with multi-drug resistance in lung cancer (Wangari-Talbot and Hopper-Borge, 2013) using nJSD score (**Figure 2D, Supplementary Figure 16A**) and found consistent patterns of regional heterogeneity. For instance, in P4, regions R3 and R4 showed higher levels of deregulation of mTOR pathway relative to that in other regions, and incidentally R3 and R4 also had gene expression signatures of high nJSD score in hypoxia and proliferation-related genesets. We further evaluated a 25 gene expression signature of resistance to Pemetrexed (Hou et al., 2012), a widely used drug for treatment of NSCLC patients, such that higher expression of the signature genes are associated with poor response. Pemetrexed-resistance signature showed intra-tumor heterogeneity - in P4, once again R3 and R4 showed signatures consistent with higher predicted resistance (**Supplementary Figure 16B**). Coherent patterns of regional heterogeneity in multi-drug resistance not only re-emphasizes need of considering intra-tumor heterogeneity while designing therapies but also power of integrating multi-level heterogeneity for meaningful inferences focusing on better treatment.

Using gene expression data for subtype-associated biomarkers we examined classical, basal, secretory, primitive-like lung cancer subtype characteristics within and across tumors (**Figure 2D, Supplementary Table 11**). Classical subgroup is associated with xenobiotic metabolism functional theme and is known to be enriched in smokers as compared to non-smokers, while secretory subgroup has very distinctive functional theme of immune response, and basal subgroup shows a cell adhesion and epidermal developmental functional theme (Wilkerson et al., 2010). Two regions in P4 had basal-like characteristics whereas other three regions had more secretory type signatures. Similarly, one region in P5 (current smoker) had signatures of secretory subgrouping and others were classified as classical. P5 also showed amplification and high expression of TP63 gene, which is another key characteristic of classical subgroup. P1, P2, P4-R1, R2, R5 and P5-R4, which had enrichment for secretory subgroup also showed high immune score as compared to other samples. Taken together, regional patterns of trancriptomic heterogeneity impacted molecular signatures of proliferation, epithelial-mesenchymal characteristics, and ultimately clinically relevant expression subtypes. These regional variations at transcriptome level are suggestive of ongoing selection, which is further supoorted by analysis of single cell sequencing data, providing evidence for non-neutral evolution,.

### Comparative assessment of genomic and transcriptomic ITH

There were parallels between ITH at genetic and transcriptomic levels; intra-tumor spatial heterogeneity was generally lower compared to tumor-paired benign tissue divergence at both levels, and that the patterns of transcriptomic divergence between regions matched corresponding patterns of genetic divergence (**Figure 2B-C**). Nonetheless, differences between heterogeneity at these two levels were phenotypically relevant. We calculated Ω – ITH to tumor-benign tissue divergence ratio for each tumor at genomic and transcriptomic levels (Ω_g_and Ω_t_), and compared them to study differences in ITH patterns between the two levels. Ω was generally higher at transcriptomic level than that at the genetic level (**Figure 2F**). Samples P1, P2, P4 showed relatively higher transcriptomic heterogeneity while P5 and P6 showed proportionally lower transcriptomic heterogeneity. Interestingly, Ω_t_/Ω_g_showed consistent patterns with histological subtyping and phenotypic characteristics, with P1, P2, and P4 showing secretory characteristics, with low mean proliferative, apoptotic, and EM scores, compared to P3, P5 and P6 (**Figure 2G**). The indices and subtyping were based on different sets of genes, so the observed patterns are unlikely due to random variation in a small number of genes. Lung squamous cell cancers with secretory subtype have modest proliferative characteristics and show substantial transcriptional heterogeneity beyond that observed at genomic level. Overall, while signatures of selection was not apparent at genomic level, regional differences in proliferative characteristics point towards ongoing evolvability.

### Heterogeneity in immune cell infiltration and associated signatures

We then investigated the extent of immune cell infiltration and heterogeneity across all tumor regions. Based on computational predictions by ESTIMATE (Yoshihara et al., 2013) P1, P2 and P4 samples had relatively high immune signatures (**Figure 3A**). P5 showed regional heterogeneity in terms of the extent of immune infiltration with R4 showing much higher infiltration as compared to the other three regions. Further deconvolution of gene expression (Gentles et al., 2015) revealed that CD8+, CD4+, plasma cells and M2 macrophages were the most abundant immune cell types (**Figure 3A** and **Supplementary Table 12**). CD4+ and M2 macrophages were present consistently across all tumor regions, whereas CD8+ T cells showed more regional variations. We constructed dendrograms using estimated proportions of different immune cell types, and found overall high intra-tumor immune heterogeneity (**Figure 3B**). We were also able to detect multiple potential T cell clonotypes based on T cell receptor status (Shugay et al., 2015), but few of them were ubiquitous (**Supplementary Figure 17**). We note that T cell clonotype identification is computationally non-trivial, especially when T cell infiltration is low. But in general, the results indicate regionally heterogeneous tumor infiltrating lymphocytes.

**Figure 3.**
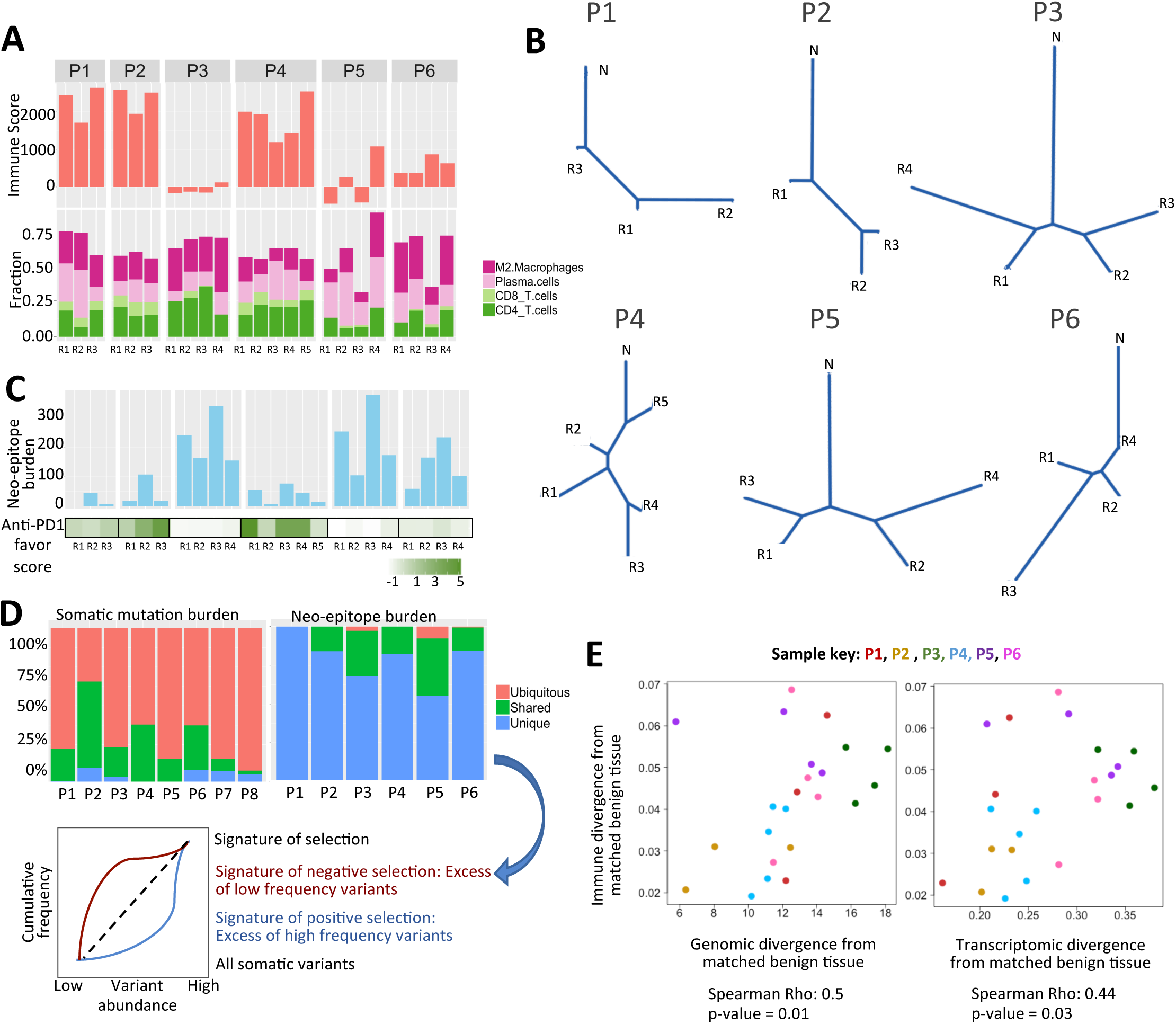
Assessment of intra-tumor immune heterogeneity in lung squamous cell cancer. **A)** Estimated immune score (upper panel), relative proportion of different immune cell types including M2 macrophages, plasma cells, CD4 and CD8 T cells (bottom panel). **B)** Dendrograms represent similarity (represented by branch length) between different regions for each patient sample based on estimated immune cell fractions in different regions. **C)** Neo-epitope (total) burden, as predicted using mutation and expression data, in different regions of all patient samples (upper panel), anti-PD1 favor score, measure of responsiveness against anti-PD1 therapy, in different regions of all patient samples (bottom panel). **D)** Somatic mutations in the tumor samples and those that are designated as neo-epitopes are grouped as ubiquitous, shared, and unique depending on their regional presentation. An excess of ‘unique’ epitopes relative to the patterns observed for all somatic variants is indicative of negative selection on the neo-epitopes. **E)** Scatterplot comparing the extent of immune and genetic divergence (left panel), and the extent of immune and transcriptomic divergence (right panel) for different tumor regions from their respective matched benign tissues. Color codes are same as Figure 2A.

Next, we analyzed both HLA and somatic mutation status (see **Methods** for details) in different tumor regions to determine neo-epitope burden. We inferred HLA types for each tumor sample conservatively using both exome and RNAseq data (**Supplementary Table 13**), and those detected in both were considered further. Both HLA I and II genes had detectable expression as shown in **Supplementary Figure 18**. Further, using HLA types, mutation burden and expression data, neo-epitopes were predicted for all the regions across all tumors (See methods). Tumor neo-epitope burden (TNB) varied from 8 to 341 per region across different samples (**Figure 3C**), of them ∼10% were classified as high affinity neo-epitopes (**Supplementary Figure 19**). Neo-epitope burden could not be calculated for P1-R1 because no HLAs were detected both at genomic and transcriptomic levels that were also present in NetMHCpan database. The neo-epitopes were classified into three categories viz. ubiquitous, shared and unique depending upon their abundance in different regions of same tumor. We observed that 55-100% of the neo-epitopes were present in only one tumor region and only up to 7% were ubiquitous (**Figure 3D**). This indicates high level of intra-tumor heterogeneity in neo-epitope burden, unlike that observed at the level of somatic mutations (**Figure 1B** and **3D**), even though tumor mutation burden (TMB) and tumor neo-epitope burden (TNB) were highly correlated (Spearman correlation coefficient = 0.82, p value: 2.1e-06; **Supplementary Figure 20**). Reduction in the burden of neo-epitopes that are present in multiple tumor regions, relative to that observed for somatic mutations is suggestive of immune-mediated negative selection purging tumor cells which had mutations that could elicit immune response. Interestingly, as reported earlier (Rosenthal et al., 2019), we also observed that TNB negatively correlated with immune infiltration (Spearman correlation coefficient: −0.85; **Supplementary Figure 21**) and was high in samples with low immune infiltration (P3, P5 and P6), and vice versa. This was not biased by mutation burden, and the negative correlation was significant even after adjusting for TMB (Spearman partial correlation coefficient: −0.57; p-value: 5.1e-03). We further analyzed expression status of 28 genes associated with anti-PD1 response (see Methods for details) in these samples. The samples with low immune infiltration also showed low anti-PD1 favor score (**Figure 3C and Supplementary Figure 22**). These observations, together with absence of ubiquitous, dominant T cell clonotypes suggest that these appear to be cold tumors in terms of their potential to elicit strong response to immunotherapy. Recent report (Cristescu et al., 2018) shows that both tumor mutation burden and T-cell inflamed gene expression profile (GEP) are needed to elicit a strong immune response, and lung squamous cell carcinoma is known to generally escape immune surveillance (Thorsson et al., 2018).

We observed high intra-tumor immune heterogeneity, as shown by dendrograms drawn using both neo-epitopes (**Supplementary Figure 23, Supplementary Table 14**) and computationally estimated proportions of different immune cell types (**Figure 3B, Supplementary Table 14**), although the latter was relatively moderate than the former. We also computed Shannon’s and Gini-Simpson’s indices for each region to quantify intra-region species richness and diversity in terms of abundance of different immune cell populations (**Supplementary Figure 24**). These intra-region diversity and richness measures at the level of immune features have higher values and showed greater regional variation than that observed at genetic level (**Supplementary Figure 11**), which is consistent with recent reports (Jia et al., 2018). Further comparative assessment of ITH at genetic, transcriptomic and immunogenic levels show that in general, ITH patterns at the levels of immune cell abundance were high and less similar to the genomic and transcriptomic ITH patterns, than latter were to one another. We compared the extent of divergence of tumor regions from paired normal regions, and found that immune divergence correlate with genomic divergence weakly (Spearman correlation coefficient 0.5, p-value = 0.01; **Figure 3E**). In a similar note, comparing Ω between genomic, transcriptomic, and immune levels, we found no strong correlation between heterogeneity patterns at immune and other levels (**Supplementary Figure 25**).

Overall, abundance of immune cell types showed ITH higher than that at other levels, and joint patterns of regional variations in immune cell infiltration and neo-epitopes suggest that immunological pruning of tumor cell populations by neo-epitope depletion impose negative selection during tumor evolution, an emerging theme observed elsewhere as well (Jia et al., 2018; Zhang et al., 2018).

### Meta-heterogeneity and regional difference in histopathological characteristics

Overall, regional differences in genomic, transcriptomic, and immune signatures appear to be broadly consistent, and collectively impact clinically relevant histopathological characteristics. As an example, we showcase patient P4, a male, 56 year old ex-smoker with a stage IIIA malignancy (**Figure 4A**). The five geographically distant regions profiled from the patient had regional differences in histological characteristics and immune infiltration, which was reflected in gene expression based classification, with R1, R2, R5 designated as secretory, and R3, R4 designated as basal subtype. R3 and R4 also had higher proliferative potential than other regions and more epithelial like features which corroborated with their basal subtype classification.

**Figure 4.**
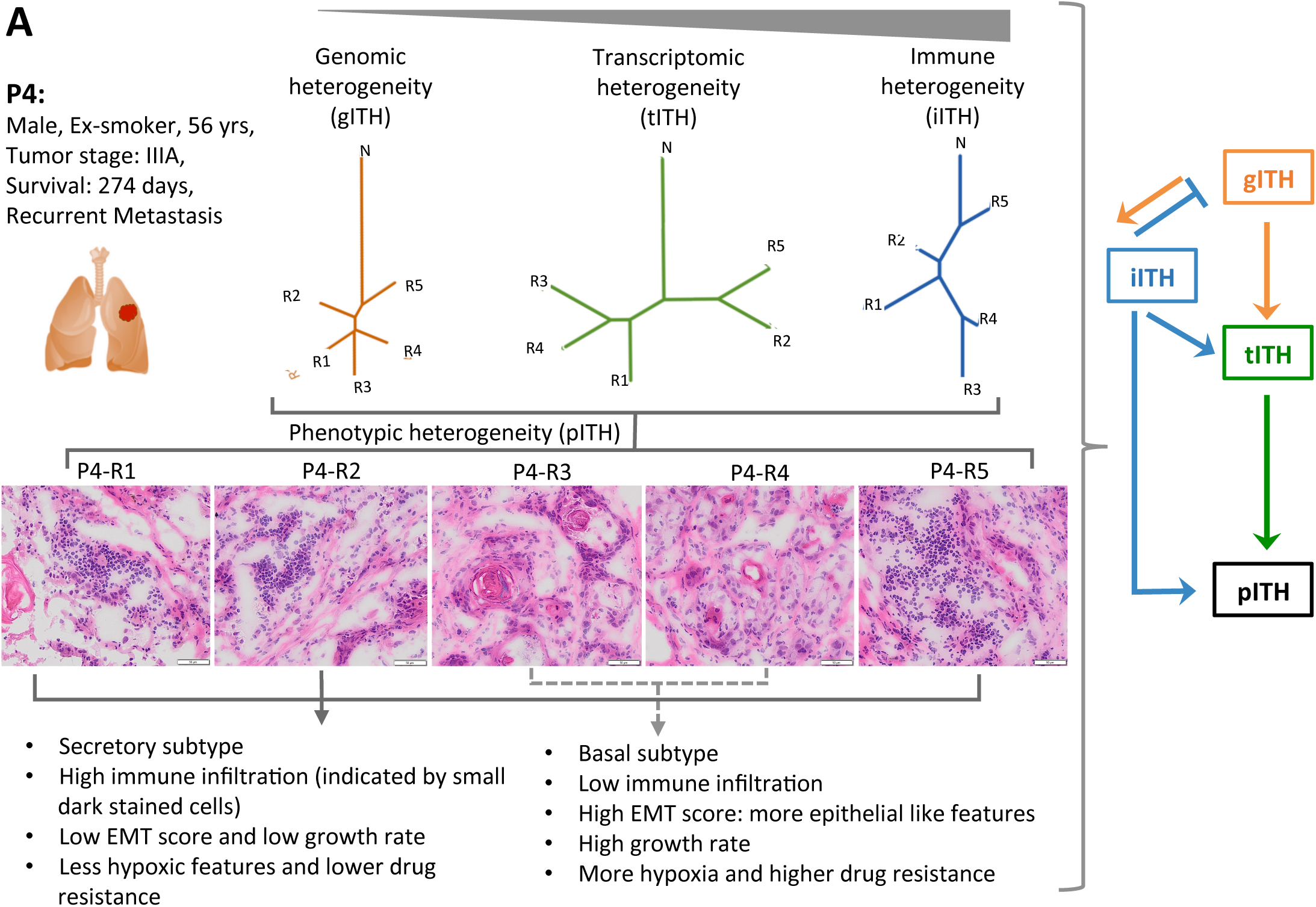
Impact of multi-level intra-tumor heterogeneity. **A)** As an example, multi-level intra-tumor heterogeneity in patient P4, a 56 year old, male ex-smoker patient with stage IIIA tumor is presented. Regional variations in histological characteristics and immune cell infiltration correlate with predicted subtype characteristics and immune scores. Furthermore, the tumor shows regional variations in proliferation and apoptosis scores, indicating coherence in multi-level intra-tumor heterogeneity. H&E stained slides for different regions of P4 have been shown with scale bars of 50um at bottom right of the images.

This observation for P4 is in line with the comparative analyses of ITH at multiple levels in Figure 2 and 3, where we observe that transcriptomic and immune-level heterogeneity rather than genetic heterogeneity correlates with intra-tumor regional differences in histopathological characteristics. Furthermore, immune infiltration negatively correlates with the mutation burden at the genetic level. These observations show that ITH at different levels are related, and impact phenotypic characteristics (**Figure 4B** and **Supplementary Figure 1**); yet their complex inter-relation may not be sufficiently captured by assessment of ITH at any one level.

### Differences in the burden of subclonal clock-like signature

Clock-like mutation signature #1 arises due to replication-associated processes, providing estimates of age of the clones in terms of the approximate number of mitosis (**Supplementary Figure 26A**) - the subclones with larger clock-like burden likely arose late. Burden of clock-like signature was calculated as weight of signature 1 for a subclone * number of mutations private to that subclone. The burden of clock-like signature differed between the mutation clusters in the samples. For instance, in Sample P5 cluster#0 had the highest burden of clock-like signature while mutation cluster #1 had the highest abundance among all clusters in all regions. (**Supplementary Figure 26B**). We further analyzed relevant data for the TracerX cohort(Jamal-Hanjani et al., 2017), which analyzed clonal phylogeny for 100 non-small cell lung tumors. For each tumor, we identified the subclones that are terminal leaves of the clonal phylogenetic tree, and tabulated their private somatic mutations, and estimated the burden of clock-like mutation signature, which accumulates progressively with cell division. In a majority of the tumors, detectable subclones had large differences in the estimated burden of clock-like signatures (**Supplementary Figure 26C**), and there was no correlation between the burden of clock-like signature and mean cell fraction of the subclones in the tumors - suggesting that the older subclones did not necessarily have larger cell population, such that these subclones are unlikely to have similar growth rates. Anyhow, we are aware of the challenges associated with inference of clonal architecture and clock-like signatures, and thus did not make explicit inference of subclonal age from genomic data.

Analyzing TCGA LUSC genomic and proteomic data, we note that tumors with higher proliferation rate (Cyclin B1 expression) tend to be of advanced stage and have a high number of driver mutations, and that high proliferation rate is associated with high somatic mutation burden as well (**Supplementary Figures 27 - 29**). But inferred phylogeny and clock-like signatures provide indirect indications of tumor evolutionary dynamics under certain assumptions, and single cell measurements can provide further insights into tumor evolution.

### Heterogeneity analysis at single cell resolution

We analyzed single cell RNAseq data from tumors of 5 non-small cell lung cancer patients (**Figure 5A**). For each tumor, samples from the core, middle, and edge regions were analyzed. 1,472-18,496 single cells were profiled per patient using 10X Genomics platform (**Figure 5B**), and based on single cell gene expression signatures, cells were annotated as tumor, immune, alveolar, fibroblasts cells etc. (Lambrechts et al., 2018). Based on single cell RNAseq data we inferred somatic copy number alteration status in tumor cells and clonal architecture using HoneyBadger (see Methods for details). The central idea behind the reconstruction is that copy number gain or loss of genomic segments are expected to affect expression of consecutive genes systematically on that segment, and subclonal cell populations sharing the same copy number alteration would show similar changes in expression of affected genes.

**Figure 5.**
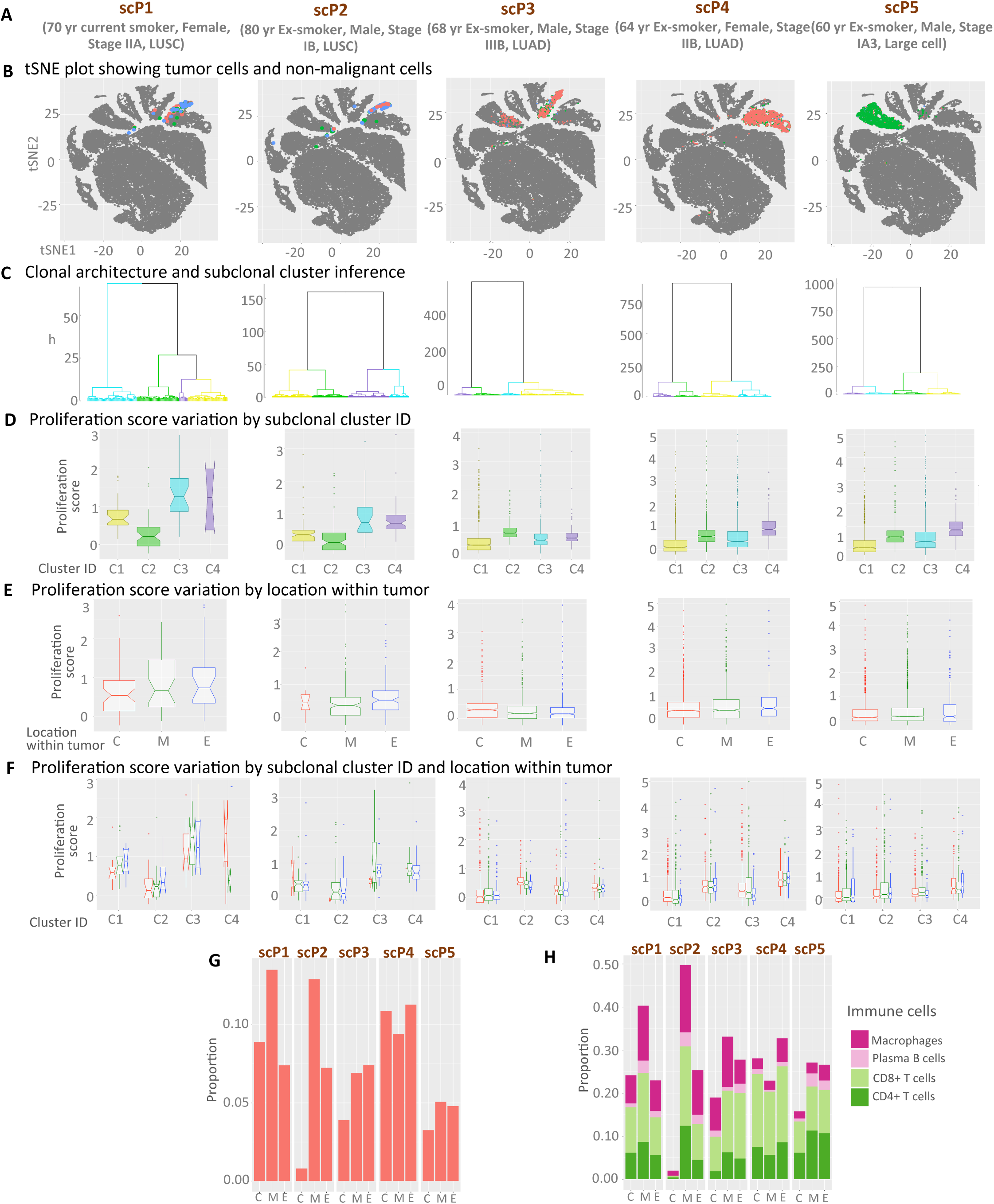
Intra-tumor heterogeneity at single cell resolution. **A)** Clinical information of 5 patients for which multi-region single cell sequencing data was performed by Lambrechts *et al.*. **B)** tSNE plots show the transcriptomic heterogeneity of the tumor cell populations, and also non-malignant cell populations for reference. Tumor cells from different regions - Core, Middle, and Edge of each tumor are colored with red, green, and blue, respectively, while all non-malignant cells are shown in grey. **C)** Tumor clonal architecture inferred from single cell RNAseq data-guided CNV calls. For each tumor 4 major subclonal clusters are numbered C1, C2, C3, and C4, marked with different colors. **D)** Boxplots showing distribution of proliferation scores of all tumor cells grouped by their subclonal cluster membership. **E)** Boxplots showing distribution of proliferation scores of all tumor cells grouped by different geographical regions - core(C), middle (M), and edge (E). **F)** Boxplots showing distribution of proliferation scores of all tumor cells grouped by different geographical regions and cluster ID. Color-codes are consistent between B, E, and F, and also between C and D. **G)** Proportion of immune cells out of total cells isolated for each patient. **H)** Proportion of different tumor relevant immune cell populations out of total immune cells for each tumor.

Clonal architecture inference indicated that the tumors in our cohort had a small number of distinct subclonal clusters, with minor differences among the cells within those subclones - as evident from the branch-lengths of the inferred tumor phylogenetic tree (**Figure 5C**). We compared top 4 major subclonal clusters down the phylogenetic hierarchy for systematic differences in scores for proliferation and other pathways discussed in Figure 2, but the results were similar for alternative cut-offs. Significant differences in proliferation rate among the sublones in a tumor would be suggestive of non-neutral clonal dynamics. At this end, for each tumor cell, we calculated pathway-level score for proliferation-related genes, as before, and then compared the distributions of the score among sets of tumor cells grouped by their subclonal membership and/or tumor location. Most subclones were ubiquitous i.e. had high number of representative cells in tumor core, middle, and margin regions. We found that in a majority of the tumors, there were significant differences between the subclones in their proliferation scores, and these differences were generally consistent across the tumor regions (**Figure 5D and 5E and Supplementary Figure 30**; nested ANOVA p-values are listed in **Supplementary Table 15**). Owing to challenges in calling mutations from single cell sequencing data, we did not attribute such growth differences to subclonal driver alterations. But there were similar differences between the subclones in other oncogenic pathways as well. For instance, replication stress is a major contributor to apoptosis (Hills and Diffley, 2014; Lukášová et al., 2019), and we found consistent differences in apoptotic pathway scores between the subclones in the tumors (**Supplementary Table 15**) - which is also concordant with observations made using multi-region bulk sequencing data (**Figure 2D and Supplementary Figure 31**) and TCGA data (**Supplementary Figure 15**). Nonetheless, some other cancer-related pathway activities also showed significant intra-tumor regional differences, even within the same subclones (**Figure 5F** and **Supplementary Figure 32**). For instance, hypoxia score was typically higher in the tumor core, while EMT scores were typically lower at the edge (**Supplementary Figure 30**). Similarly, proliferation score for the subclones were typically lower in the core relative to that in the tumor periphery - which is consistent with the observations that much of the tumor growth occurs near the tumor margin. This suggests that regional contexts can affect proliferative characteristics and growth dynamics within a tumor, beyond that attributed to subclonal differences.

Although the tumors were genetically rather homogeneous i.e. most major subclonal clusters had sufficient presence in all tumor regions (**Figure 5F**), we found large intra-tumor regional differences in the overall burden of infiltrating immune cells (**Figure 5G**), and the abundance of different immune cell types therein (**Figure 5H**). This corroborates the observation made using bulk sequencing data that genetic heterogeneity does not sufficiently captures the extent of intra-region non-genetic heterogeneity.

## DISCUSSION

In this study, we investigated intra-tumor heterogeneity (ITH) patterns in lung squamous cell carcinoma at levels of genomic, transcriptomic, tumor-immune cell interactions, and histological characteristics by multi-region profiling and single cell data analysis to compare ITH at different levels. We have three major observations.

First, our findings show that even though the extent of ITH differs between genetic and non-genetic levels, regional patterns of heterogeneity at these levels are biologically consistent. Despite high somatic mutation burden, data from others and us showed that spatial pattern of genetic ITH in lung squamous cell carcinoma is moderate, which might be due to spatial mixing of tumor subclones. In contrast, regional patterns of trancriptomic and immune heterogeneity were generally higher, indicating non-genetic sources of variation (Sharma et al., 2018), and impacted key cancer-related pathways, subtype characteristics, and proliferative potential. We acknowledge potential caveats. Limited sampling of tumor regions provides incomplete information about the tumor cell populations. Second, a tumor doesn’t grow as an expanding sphere; rather tissue structures, blood vessels, and other constraints influence tumor growth. Moreover, upon resection, tumor tissues deform - such that geographic distance has modest relevance and must be considered with care during computational modeling of tumor growth. Nonetheless, co-extraction of DNA and RNA from the same tumor cores minimized between-omics-level differences and allowed us to compare the topologies of the genomic, transcriptomic, and immune-level denograms. Anyhow similarity of dendrograms at genetic and transcriptomic level, and correspondence between purity-adjusted copy number and gene expression suggested that genetic heterogeneity is not buffered at expression level (**Figure 4B**). We observe coherent regional differences at other instances as well, including TERT expression v/s telomere length, neo-epitope burden v/s immune cell infiltration that indicate biologically consistent patterns of variations across multiple levels.

Second, phenotypic heterogeneity in terms of histopathological characteristics was not effectively captured at any one level; rather intra-tumor heterogeneity at different levels collectively impact tumor histological features. For instance, regional differences in gene expression, immune signatures, and subtype classification matched histological features (**Figure 4A** and **Supplementary Figure 1**) - indicating that intra-tumor functional heterogeneity is manifested in regional differences in histopathological characteristics. Presence of multiple different subtype characteristics within the same tumor have been reported elsewhere, but our study provides evidence linking regional differences in histological characteristics with transcriptomic and immune level heterogeneity (**Figure 4B**).

Third, integrative assessment of genetic and non-genetic heterogeneity patterns provide evidence for non-neutral evolutionary dynamics in these tumors (De and Ganesan, 2017). For instance, the burden of clock-like signatures suggested that detectable subclones likely arose at different times during tumor evolution. In a complementary analysis, we found that proliferation rate varied between subclones, indicating that those might have different growth dynamics. Interestingly, proliferation and other cancer-related pathways also showed significant intra-tumor regional differences, even within the same subclones. For instance, proliferation rate for most subclones was relatively lower in the tumor core. This has interesting implications for tumor evolution. For instance, if two subclones that have otherwise comparable growth rates in the same tumor context, exist in two geographic regions in a tumor, such regional differences in proliferation can influence their net population-level growth rates, allowing one subclone to be eventually more dominant over the other in the tumor with time. Alongside, intra-tumor immune heterogeneity patterns suggested that tumor-immune cell interactions impose negative selection by pruning the tumor cells carrying neo-epitopes that elicit strong immune response. These patterns are consistent with predator-prey dynamics between immune and tumor cells, and suggest that immunological pruning of tumor cell populations by neo-epitope depletion imposes negative selection on genetic variations during tumor evolution (Jia et al., 2018; Zhang et al., 2018).

Overall, our study shows that despite having coherent patterns of ITH at different genetic and non-genetic levels, multi-level assessment of heterogeneity is necessary to identify the determinants of phenotypic heterogeneity in clinically relevant characteristics, and to appreciate the roles microenvironment plays in influencing the mode of tumor evolution.

## METHODS

### Biospecimen collection and multi-region genomic profiling

We obtained surgically resected, fresh frozen tumor and matched normal tissue specimens from 9 de-identified, stage IX-IIIB lung squamous cell carcinoma patients under an IRB approved protocol. Details of the samples are provided as **Supplementary Table 1A**. For each tumor 3-6 different regions that were geographically distant were identified, and biopsied to obtain tissue sections. When possible, H&E staining of tumors was done to identify tumor rich regions in order to take biopsies. In total, 42 biopsies (10 normal and 32 tumor) were processed in this study. Pathological tumor purity was 40-80%. DNA and RNA were co-extracted from 29 biopsies (6 normal and 23 tumors) using TRizol method and were further processed for exome and RNA sequencing. For 8 other biopsies (2 normal and 6 tumors – from 2 patients) only DNA was isolated using Qiagen DNeasy kit and exome sequencing was performed. For 8 other samples (1 normal and 3 tumor regions each from 2 patients) only DNA was isolated and targeted panel sequencing (Agilent SureSelect run on a HiSeq) was performed for 257 genes. Details of all samples are provided in **Supplementary Table 1A**. For Exome and RNA sequencing (rRNA depletion) were performed on Illumina HiSeq 2000 using 150bp paired-end protocol.

### Genomic data analysis

FastQC (v0.11.7) was used for initial quality checks, and low-quality reads and PCR duplicates were removed. Next, we used BWA-mem (Li and Durbin, 2009) (v0.7.17-r1188) to map the reads onto human genome (GRCh38), and call variants using varScan2 (Koboldt et al., 2012) (mapping quality > 40, base quality > 20). The average sequencing depth ranged from ∼50-191x for exome-seq (**Supplementary Table 16**). Variants present in dbSNP or those with strand bias were excluded, and only ‘high confidence’ somatic variants with tumor allele frequency >5% at least in one tumor region and normal allele frequency <1% were selected. A vast majority of the reported somatic variants had no read support in matched normal tissues. For each somatic variant deemed as high confidence variant in at least one tumor region, we queried the corresponding base-position in other tumor regions in that tumor specimen, and if the variant allele was supported by reads with mapping quality >20 and base quality >25 and variant allele frequency > 2%, it was included (**Supplementary Table 2**). All identified somatic mutations were annotated with SnpEff (Cingolani et al., 2012) (v4.3t). Missense, nonsense, frameshift, or splicing mutations in known COSMIC cancer genes with high-predicted impact were marked.

We used desonstructSigs (Rosenthal et al., 2016) to identify patterns of mutational signatures on somatic variants. Mutational signatures were inferred for ubiquitous and non-ubiquitous (shared + unique) variations separately, also all variations together (ubiquitous + shared + unique). Also, analysis was done both using all 30 signatures as well as selected signatures relevant to lung cancer biology (Alexandrov et al., 2016).

Copy number variation analysis was done using FACETS (Shen and Seshan, 2016) based on exome-sequencing data. Reads with mapping quality >15 and base quality >20 were considered for CNV analysis, and genomic regions with inferred duplication, deletion, or LOH were identified. Whole genome duplication and ploidy-level changes were inferred based on chromosome-wide, haplotype-aware copy number inferences.

To infer telomere length in the tumor regions, we used TelSeq (Ding et al., 2014), which considers the reads with >= 7 TTAGGG/CCCTAA repeats to provide an estimate of telomere length. Telomere length was estimated for both tumor regions (TTL: Tumor Telomere Length) as well as normal (NTL: Normal Telomere Length). We reported telomere length for different tumor regions relative to their matched normal region from the same donor as log_2_ of fold change between tumor and normal [log_2_ (TTL/NTL)].

Targeted sequencing was done for two samples (P8 and P9) for 257 genes. Somatic mutations were called in different regions as described above. Heatmaps and dendrograms were made as described below. **Supplementary Tables 17, 3, 5** show targeted sequencing gene list, somatic mutation calls and distance matrices respectively.

### Heatmaps, genetic dendrograms, and heterogeneity estimates

Heatmaps, depicting regional abundance of somatic SNVs, were drawn using variant allele frequency (vaf) data for somatic SNVs across all regions of a tumor. Pairwise distance between the regions *d_ij_* in a patient was computed as distance in terms of difference in tumor-purity-adjusted variant allele frequency (vaf) for all detected SNVs, as d_ij_ =Σ_k_**_∈_**_V_|s_ik_-s_jk_ |/V where *s_ik_* is the vaf of *k*^th^ variant in *i*-th region in a tumor. Then multiregional tumor trees (unrooted dendrograms) were drawn using this distance metric using neighbor joining. The dendrograms were bootstrapped using *boot.phylo* function in the *ape* R package. Distances between pairs of regions in the trees represent the extent similarity between different regions of the same tumor at the genomic SNV level. We computed = Σi≠B l_i_/l_i=B_ where *l_i=B_* is the branch length of the phylogram from the healthy tissue to its nearest node, providing an estimate of the ratio of intra-patient regional diversity to tumor-benign tissue divergence.

For each tumor specimen, based on the catalog of somatic mutations and copy number data across multiple regions variant clusters were identified using PyClone (v0.13.1) (Roth et al., 2014). Default parameter setting was used to identify somatic mutation clusters. For downstream analyses we only analyzed mutation clusters of size ≥3 variants. Key conclusions were unaffected by the choice of threshold. Intra-region diversity and richness at genetic levels were computed using Shannon’s and Gini-Simpson’s indices on the relative abundances of the mutation clusters in each tumor region.

### Identification of subclones and evolutionary dynamics parameters utilizing Bayesian framework

We also used a Bayesian approach developed by Williams et al (Williams et al., 2018), as implemented in their SubClonalSelection.jl to draw clonal inferences from variant allele frequencies. Subclonal architecture of each sample was detected and the selective advantage (s) and time of appearance of each subclone (t) were measured simultaneously, as recommended by the authors. Input parameters for SubClonalSelection.jl were set as default and fitted with specific sample (read depth, minimum/Maximum range of VAF, minimum cellularity), specifically, mutation rate as the input parameter was the effective mutation rate per tumour doubling. All simulations were run for a 10^6 iterations.

### Transcriptomic data analysis

After initial quality-checks of the raw RNA sequencing reads using FastQC (v0.11.7), and removal of any low quality reads, STAR aligner (v 2.6.0c) (Dobin et al., 2013)was used to map the remaining reads onto human genome (GRCh38). RSEM (v1.3.1) (Li and Dewey, 2011) was used for transcript quantification, and log_2_ (TPM+1) (Transcripts per million) values were reported for different tumor regions and also matched benign regions. Principal Component Analysis (PCA) and hierarchical clustering was done on log_2_ (TPM+1) using FactoMineR package. ESTIMATE (v1.0.13)(Yoshihara et al., 2013) was used for predicting tumor purity, and the presence of stromal/immune cells in tumor tissues.

We measured the extent of perturbation in cellular pathways using a metric called tITH calculated using network based Jensen-Shannon Divergence (Park et al., 2016). nJSD was applied as a distance measure between two network states. For each pathway, tITH was defined based on two distance values, distance from tumor to benign tissue (*NT*) and distance from tumor to maximally ambiguous network (*TA*) into a single entropy-based metric of pathway-level dysregulation as follows:

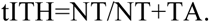

This metric was calculated for relevant pathways in each tumor region across all patient samples. Same score was used in assessing multi-drug resistance pathway activity.

We computed epithelial vs mesenchymal (EM) characteristics scores based on a published approach (Ramaker et al., 2017) using expression signature of 76 genes relevant for epithelial and mesenchymal characteristics with minor modification. Of the gene signatures, we could estimate gene expression [log_2_(TPM+1)] for 49 epithelial related genes (e-genes) and 11 mesenchymal related genes (m-genes) in our samples (**Supplementary Table 10 and 18**). For each gene, we estimate its mean (µ) and standard deviation (σ) in the 6 benign tissue samples, and then in *i*-th tumor region, we computed the Z-score (*Z_i_*) corresponding to its expression, *e_i_*as *Z_i_*=(*e_i_* - µ)/σ. Then, for each tumor region, EM score was calculated as:

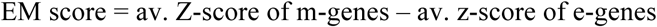

EM score was compared between regions from the same tumor. Proliferation (PI) and apoptotic (AI) indices were calculated using a similar approach using 124 proliferation-associated genes (Wilkerson et al., 2012)and 6 apoptosis-related genes. For each gene, Z-scores corresponding to its expression in tumor regions were calculated based on the mean and standard deviation of its expression in the benign samples, and then proliferation index (PI) and apoptosis index (AI) were defined as mean z-scores of all proliferation and apoptosis genes, respectively. Similar strategy was followed for calculating scores for hypoxia signature (Buffa et al., 2010) and Pemetrexed resistance signature.

mRNA expression based subgroups were also inferred for different regions of the same tumor across all samples. Predictor centroid data for different subgroups (primitive, classical, secretory and basal) was downloaded from Wilkerson *et.al.* (Wilkerson et al., 2012). For the same set of genes (as centroids, **Supplementary Table 18**), Spearman correlation coefficient between expression data of samples and centroids was calculated. Subgroup showing the highest correlation coefficient (with pval<0.01) was assigned to each sample.

Multiregional tumor trees at transcriptomic levels were constructed for every patient using RNA expression data for all genes across different regions, using an approach similar to that used at genomic level. Manhattan distance was computed between all regions of a patient sample using log_2_ (TPM+1) and then unrooted dendrograms were drawn using this distance metric.

### Immune signature analysis

Immune cell infiltrations were inferred from molecular signatures of immune cell types. ESTIMATE (v1.0.13) (Yoshihara et al., 2013) was used for predicting the level of immune infiltration in tumor tissues. CIBERSORT (Gentles et al., 2015) was used to estimate abundance of different immune cell populations from expression data. Standard LM22 signature gene file, and 1000 permutations were used to calculate deconvolution p values.

Class I and class II HLA types were predicted from both DNA and RNA data using HLAminer(Warren et al., 2012). Only most likely HLA class I alleles (Confidence (−10 * log10(Eval)) >= 30 and Score >= 500) were considered. HLA identified at both DNA and RNA levels were selected for further analysis.

MuPeXI (Bjerregaard et al., 2017) was used for predicting potential neo-epitopes (neo-epitopes) using exome and RNA sequencing data. It uses somatic mutation calls (SNVs and InDels), a list of HLA types, and a gene expression profile to predict tumor specific peptides. For any neoantigen to be classified as potential neo-epitope, it should be expressed, and have a high affinity to its respective expressed HLA alleles. All predicted neo-epitopes were classified into two categories based on their binding affinity to predicted HLA types; weak binders and strong binders. Strong binders were defined as: expression >1, mutRank < 0.5, Priority score > 0 and difference between normal and mutRank > 0.5 and weak binders as: mutRank < 2. mutRANK is %Rank of prediction score for mutant peptides as defined by netMHCpan 4.0 (Jurtz et al., 2017), which is used at backhand in MuPeXI. Based on this criteria weak binders and strong binders were predicted for all tumor regions across all samples. Further these neo-epitopes were classified into three categories based on their regional abundances; ubiquitous, shared and unique. This classification was done for both weak and strong binders separately.

TCR repertoire predictions were done on both DNA and RNA data using MiXCR (Bolotin et al., 2015) and VDJtools (Shugay et al., 2015). Paired-end RNA and exome fastq files for each sample were provided for MiXCR analysis. The exome data was analyzed using the default MiXCR pipeline parameters, and the RNA-seq files were analyzed using the pipeline and parameters recommended for RNA-seq data in the MiXCR documentation. For each sample, the clonotypes with the highest count (supporting reads) were selected from the exome data (> 3 supporting reads) and the RNA data (> 24 supporting reads). The count, fraction, and sequence of these clonotypes were then compared to other clonotypes from the same tumor to look for intratumor heterogeneity. In addition, the MiXCR output files were then converted to vdjtools format and analyzed using vdjtools to get basic statistics, spectratypes, segment usage graphs, and diversity statistics for each sample.

Anti-PD1 favor score was calculated for each tumor region using gene expression data [log_2_ (TPM+1)] for 28 genes that comprise of anti-PD1 favor signature (Gibney et al., 2016) (**Supplementary Table 18**). For each gene, Z-scores corresponding to its expression in tumor regions were calculated based on the mean and standard deviation of its expression in the benign samples, and then anti-PD1 favor score was defined as mean z-scores of the genes involved.

Similar to genomic and transcriptomic data, multiregional tumor trees were also made for every patient to infer immunogenic heterogeneity (iITH). In this case trees were made using two different datasets; i) immune cell proportions from CIBERSORT (Gentles et al., 2015) and ii) expression of neo-epitopes predicted from MuPeXI (Bjerregaard et al., 2017). Expression of predicted neo-epitopes was considered zero in a region if the neo-epitope is not detected in that region. Manhattan distance was computed between all regions of a patient sample using both the datasets independently and separate unrooted dendrograms were drawn using respective distance metrics.

### Single Cell RNA sequencing data analysis

single cell RNAseq data for 5 non-small cell lung tumors were obtained from the study published by Lambrechts et al. (Lambrechts et al., 2018) (E-MTAB-6149 and E-MTAB-6653). In brief, core, middle, and edge regions from each tumor were profiled, and single tumor, immune, and stromal cells from those regions were profiled on 10X platform. **Supplementary Table 1B** shows the details of samples used in this study. For individual tumor cells, we used the log2CPM values to calculated proliferation, apoptosis, hypoxia and EMT scores as described above, after taking Alveolar Type 2 (AT2) cells from the same patients as control.

We then used single cell gene expression data to infer copy number and clonal clusters using HoneyBadger (Fan et al., 2018). In brief, the inference is drawn by considering that copy number gain or loss would result in systematic increase or decrease in expression of genes that are co-localized in the genome, and cells that are within the same subclonal clusters would show more similarities than those that are distant in the clonal phylogeny. Under default setting HoneyBadger first explicitly infers per-cell copy number gain or loss from expression-based raw copy number log2 ratios, before making clonal architecture inference, which is unsuitable for large datasets. So we used raw copy number log2 ratios inferred from gene expression data by HoneyBadger directly to make clonal architecture inference, bypassing per-cell explicit copy number calling. Single cell profiling allows, in principle, complete phylogenetic reconstruction, but in practice, sparse single cell RNAseq data makes deep branching inferences unreliable. So, we compared top 4 major subclonal clusters down the phylogenetic hierarchy for systematic differences in proliferation and other cancer-related pathways.

The extent of immune infiltration was calculated based on the number of different types of immune cells in each region (core, middle, edge) as annotated by Lambrechts et al. Nested anova was used to test the statistical significance of the difference in growth rates and other tumor associated features in both clusters and regions. **Supplementary Table 19** gives details of proliferation, apoptosis, hypoxia, EMT scores in different regions and mutational clusters for all patients. Classification of different cells into different genomic clusters is provided in **Supplementary Table 20. Supplementary Table 21** shows the number and proportion of different immune cells in different regions for all patients.

### Histopathological screening of H&E stained slides

High-resolution digital images of Hematoxylin and Eosin (H&E) stained sections were evaluated by a board certified pathologist blinded to genomic data. Each image was scored for percent tumor nuclei and for the presence of lymphocytes on a scale of 1 (minimal) to 3 (robust) over the entire section.

### MetaITH pipeline

metaITH - a computational pipeline (https://github.com/sjdlabgroup/metaITH) provides an interface to perform meta-analysis of intra-tumor heterogeneity patterns across different levels, and for estimating transcriptomic signatures of disease-relevant biological processes. Heterogeneity-related utilities include heatmaps, dendrograms, and measures of intra-tumor divergence and diversity at different levels. Signature-related utilities include geneset signatures for proliferation, apoptosis, hypoxia, multi-drug resistance, anti-PD1 favor, but those could be also used to estimate other user-defined signatures as well.

## AUTHOR CONTRIBUTIONS

SD conceived the project. AS, SD designed the experiments. AS, AC performed the experiments. AS, EM, XH analyzed the data. AS, JM, GR, SD interpreted the data. AS, SD wrote the manuscript with input from other authors.

## ACKNOWLEDGMENTS

The authors acknowledge financial support from R01GM129066, P30CA072720, Robert Wood Johnson Foundation (S.D.). The authors thank the other members of the laboratories of S.D. and the Center for Systems and Computational Biology at Rutgers Cancer Institute for helpful discussions. The funders had no role in study design, data collection and interpretation, or the decision to submit the work for publication.

## A. Supplementary Tables (*included as separate excel worksheets)*

- **Supplementary Table 1:** Table listing clinical and pathological details of the samples involved in the study
- **Supplementary Table 2: A**) List of Varscan2 inferred exonic, somatic SNVs and InDels from exome sequencing data in all regions of all patient samples. Annotation column gives annotation details of each variation. Variants mapped to scaffolds are not reported. **B)** List of FACETS inferred Copy Number Alterations for Cancer Associated Genes (CAGs) from exome sequencing data in all patient samples. Chr: Chromosome, start: start position, end: end position, gene: gene name, tcn: total copy number state, lcn: minor allele copy number state, status: over all copy number status, Patient: patient sample ID, region: region (biopsy) id.
- **Supplementary Table 3:** Distance matrices for all samples using somatic SNVs. R1, R2, R3, R4, R5 represent tumor regions and N represent matched normal.
- **Supplementary Table 4:** Distance matrices for all samples using somatic SNVs identified from exome sequencing data. R1, R2, R3, R4, R5 represent tumor regions and N represent matched normal.
- **Supplementary Table 5:** Distance matrices for P8 and P9 using somatic SNVs inferred from targeted sequencing data. R1, R2, R3 represent tumor regions and N represent matched normal.
- **Supplementary Table 6:** Distance matrices for all samples using RNA expression data (log2(TPM+1)). R1, R2, R3, R4, R5 represent tumor regions and N represent matched normal.
- **Supplementary Table 7:** Matrix of log2(TPM+1) values from RNAseq data from all patient samples involved in the study. Lung cancer associated genes mentioned in the manuscript showing >4 or <0.25 fold change in tumors as compared to normal have been highlighted.
- **Supplementary Table 8:** Pathway-tITH entropy values estimated by nJSD for lung cancer associated pathways in all patient samples across all tumor regions compared to normal regions.
- **Supplementary Table 9:** Proliferation Score and Apoptosis Score for all samples across tumor regions calculated as described in Methods section.
- **Supplementary Table 10:** Epithelial v/s Mesenchymal characteristics score (calculated as described in Methods section). Higher score represents more epithelial like features.
- **Supplementary Table 11:** Table showing subtype classification score. It is spearman correlation coefficient between subtype centroids and our data (as described in Methods section).
- **Supplementary Table 12:** Table showing cell fractions of different immune cell types as inferred by CIBERSORT.
- **Supplementary Table 13:** HLA alleles identified by HLAminer tool from Exome (DNA) and RNAseq data (RNA). DNA-RNA overlap shows HLA alleles that were predicted from both datasets and are considered for downstream analysis of predicting neo-epitopes.
- **Supplementary Table 14: A)** Distance matrices for all samples using proportion of different immune cells as inferred by CIBERSORT. R1, R2, R3, R4, R5 represent tumor regions and N represent matched normal. **B)** Distance matrices for all samples using neo-epitopes (weak+strong). R1, R2, R3, R4, R5 represent tumor regions and N represent matched normal.
- **Supplementary Table 15:** Differences in different tumor associated features such as apoptosis, hypoxia, proliferation and EMT scores between different clusters and positions (Core, Middle and Edge) were tested using nested Anova statistics. Table below shows p-values for all samples.
- **Supplementary Table 16:** Table showing average coverage in exome sequencing data and approximate million reads in RNAseq data for all regions in all patient samples.
- **Supplementary Table 17:** List of genes used for targeted sequencing of P8 and P9.
- **Supplementary Table 18:** List of genes used in different tumor features analysis.
- **Supplementary Table 19:** Proliferation, Apoptosis, Hypoxia and EMT scores for all patients across all tumor cells from scRNAseq data.
- **Supplementary Table 20:** Cluster assignment of different cells. HoneyBadger with some in house modifications was used to infer clusters. Clusters have been assigned on different thresholds; 2, 3, 4, 5, and 6 clusters. For analysis we have used 4 clusters (highlighted in yellow) per sample.
- **Supplementary Table 21:** Sample wise details of number and proportion of immune cells inferred from publicly available scRNAseq data

## B. Supplementary Figures *(included in this document)*

**Supplementary Figure 1:**
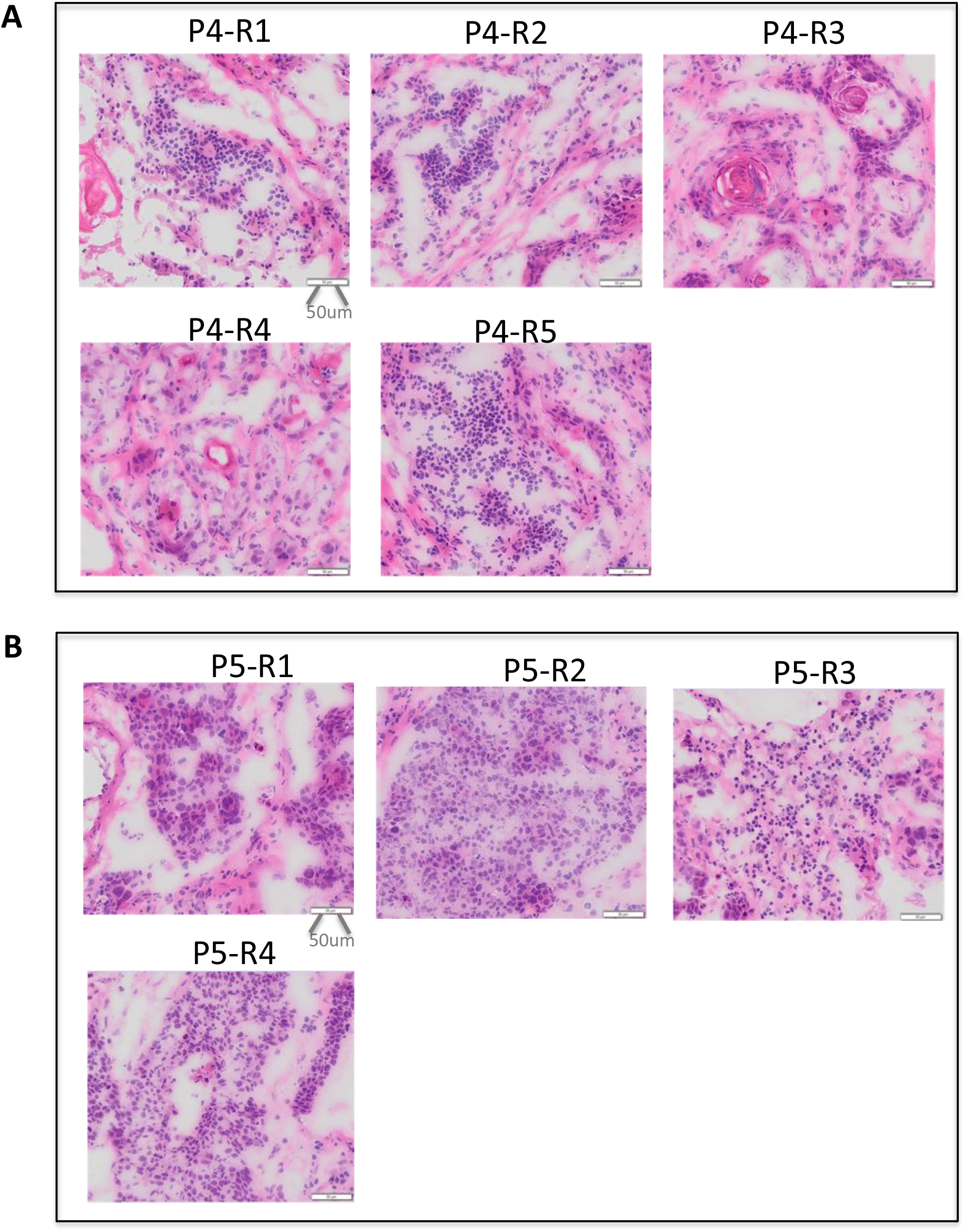
Representative images of different geographic regions from different regions are shown for tumor P4 and P5. A) P4-R1, P4-R2 and P4-R5 have higher immune infiltration as indicated by small dark stained cells, whereas P4-R3 and P4-R4 have less immune infiltration. B) Similarly, P5-R1, R2, R3 have more immune infiltration as compared to P5-R4. Scale bars of 50 um are at the bottom right of all the images.

**Supplementary Figure 2:**
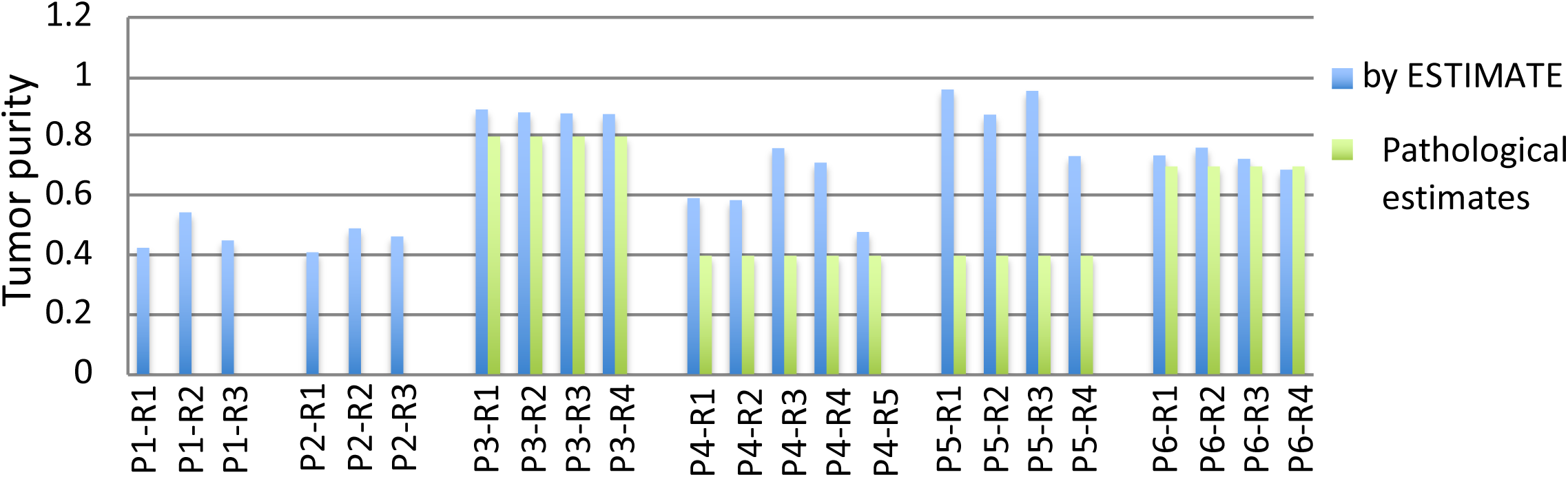
Tumor purity estimates as inferred computationally by ESTIMATE using RNAseq data and pathological estimates inferred by pathologist using H&E stained slides. Pathological tumor purity estimates were not available for all samples.

**Supplementary Figure 3:**
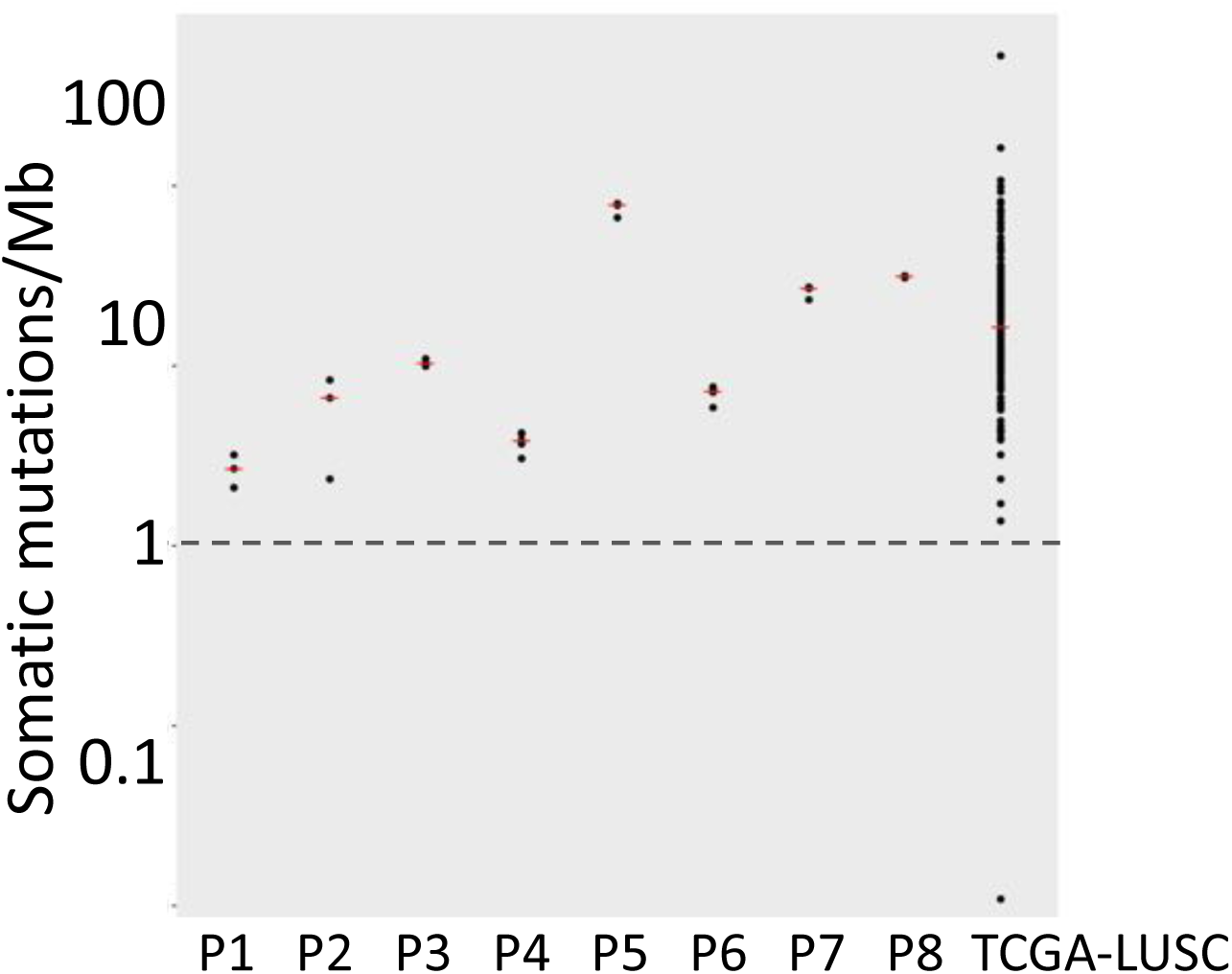
Estimated somatic mutation frequency in exonic regions for lung squamous cell carcinoma samples. Each dot represents estimated somatic mutation frequency in different regions in each tumor sample in our cohort, which is compared against the somatic mutation frequency for the Cancer Genome Atlas Lung squamous cell carcinoma (TCGA-LUSC) samples. Red bar depicts median values for each group.

**Supplementary Figure 4:**
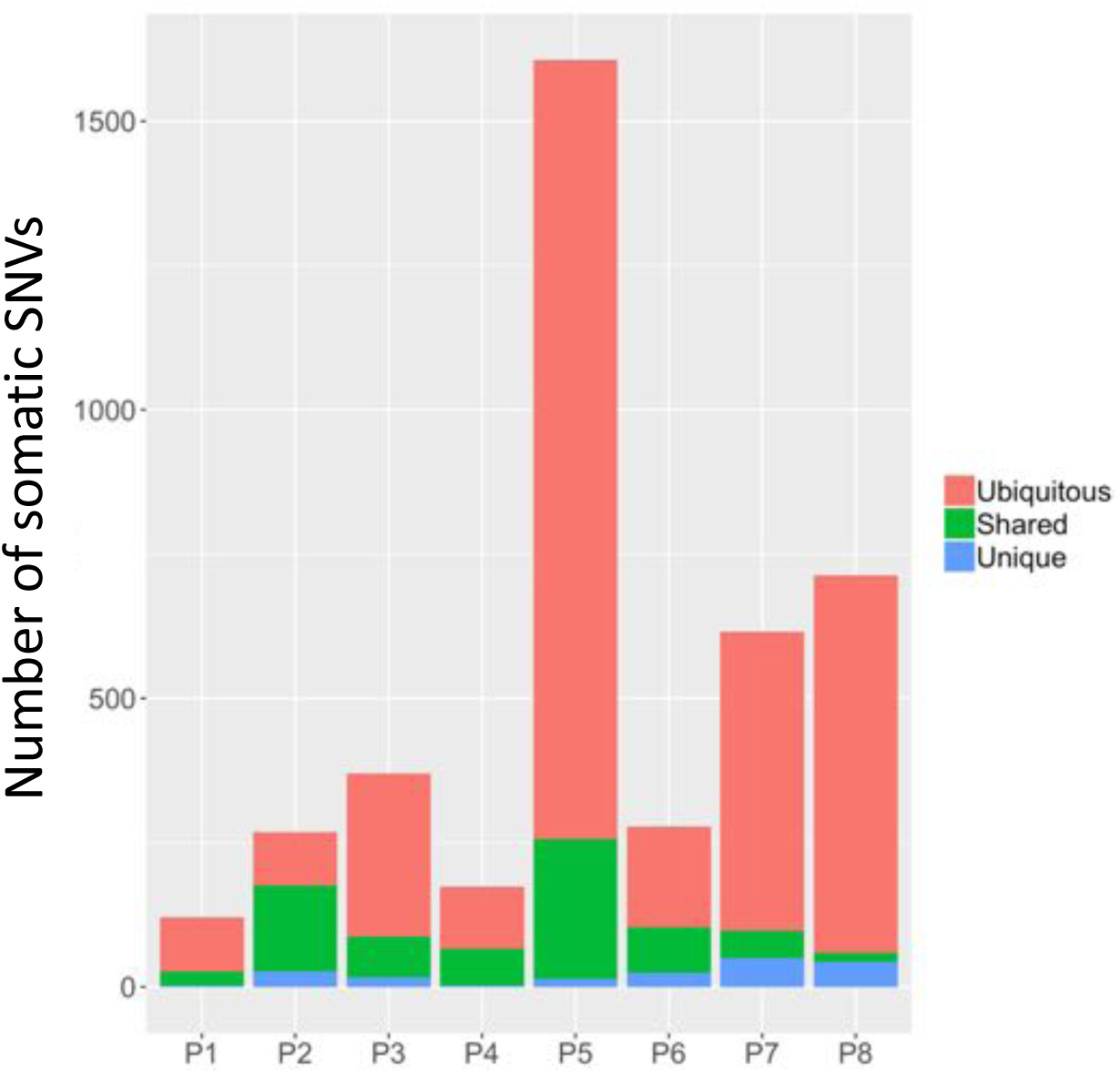
Stacked bar chart representing proportion of SNVs that are (i) ubiquitous i.e. present in all regions in a tumor sample (ii) shared i.e. present in multiple but not all regions, and (iii) unique i.e. present inly in one region profiled in a tumor. Approximately three-fourth of the somatic SNVs were ubiquitous, although the patterns varied between samples. For instance, in P2, ∼50% of the variants were shared and ∼30% were ubiquitous, while in P8, almost 90% of the variants were ubiquitous.

**Supplementary Figure 5:**
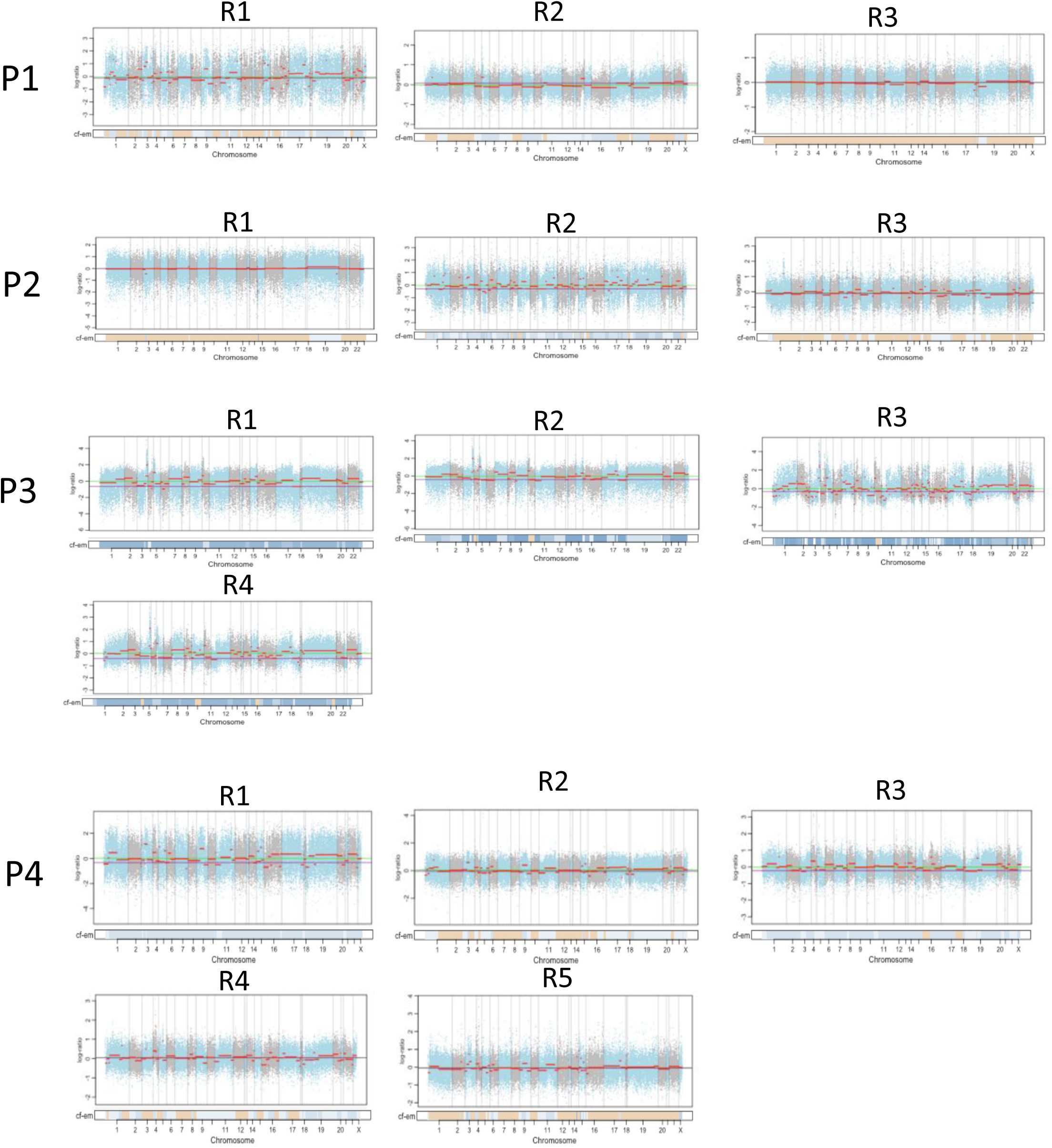
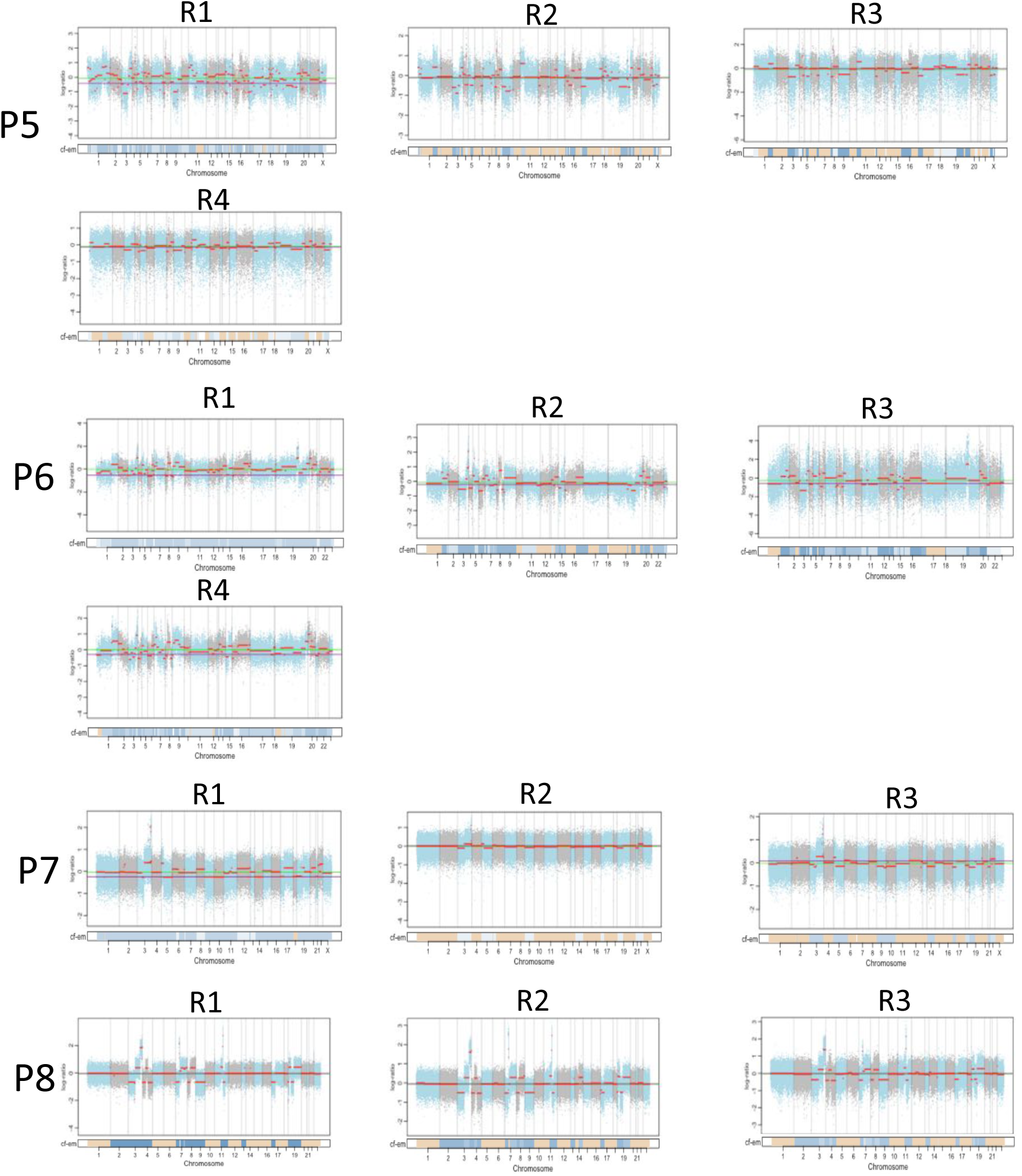
Copy number assessment of different tumor regions relative to matched benign samples based on exome sequencing data using FACETS. For each patient, copy number log ratio of total read count in a tumor region to that in the matched benign region was plotted along genomic bins in chromosome 1-22 and X, and copy number segmentation was performed. Alternating cyan and grey dots represent genomic bins from odd and even chromosomes respectively, and red horizontal bars indicate estimated copy number segments. Total lengths of chromosomal sections covered by exons differed between chromosomes, depending on gene density of respective chromosomes. Copy number plots are shown for sample P1-P4. Purple line: diploid state and green line: median. Copy number assessment of different tumor regions relative to matched benign samples based on exome sequencing data using FACETS. For each patient, copy number log ratio of total read count in a tumor region to that in the matched benign region was plotted along genomic bins in chromosome 1-22 and X, and copy number segmentation was performed. Alternating cyan and grey dots represent genomic bins from odd and even chromosomes respectively, and red horizontal bars indicate estimated copy number segments. Total lengths of chromosomal sections covered by exons differed between chromosomes, depending on gene density of respective chromosomes. Copy number plots are shown for sample P5-P8. Purple line: diploid state and green line: median.

**Supplementary Figure 6:**
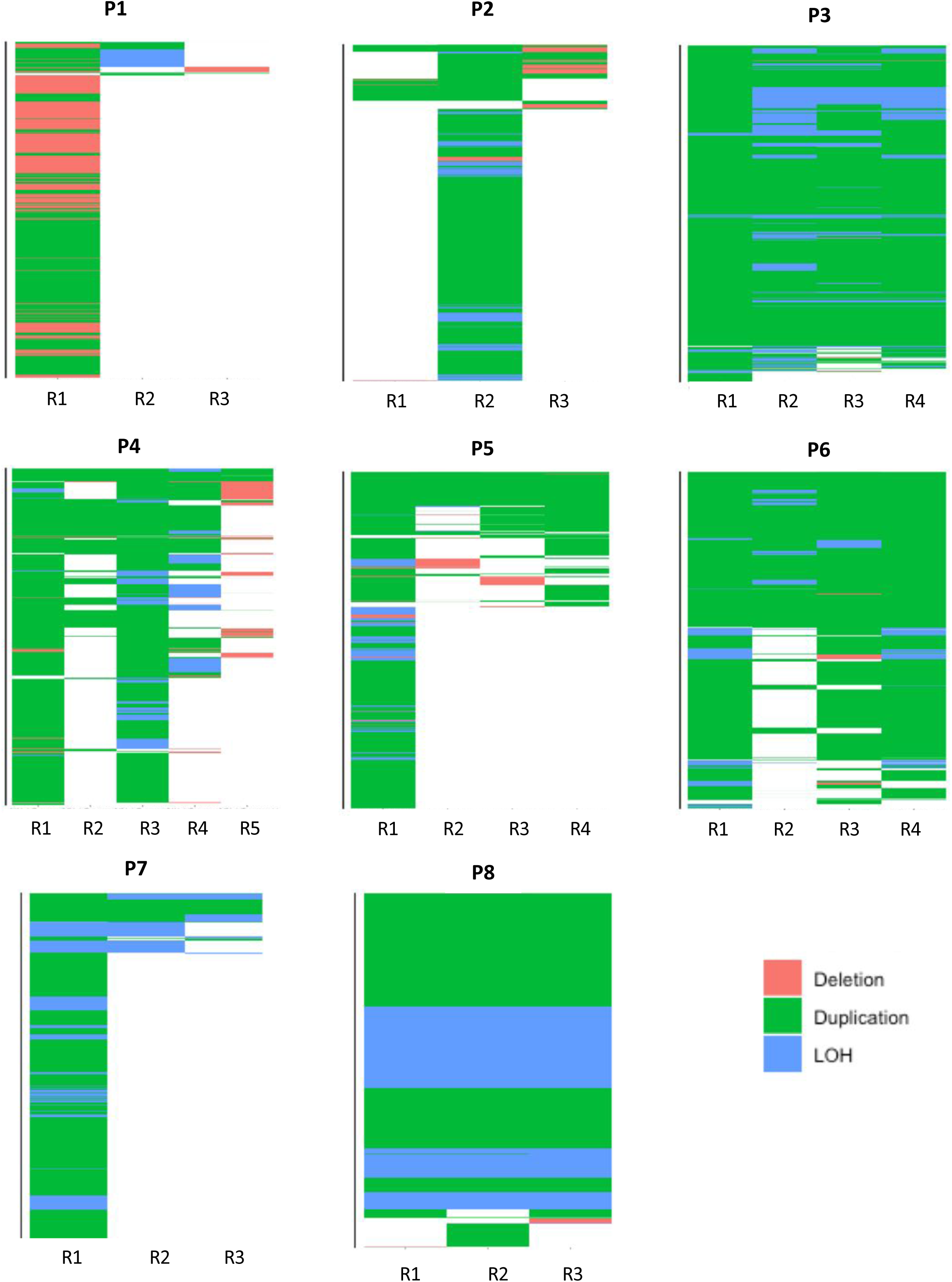
Heatmaps showing regional variations in deletion, duplications and LOH events in genes in different regions of all tumor samples.

**Supplementary Figure 7:**
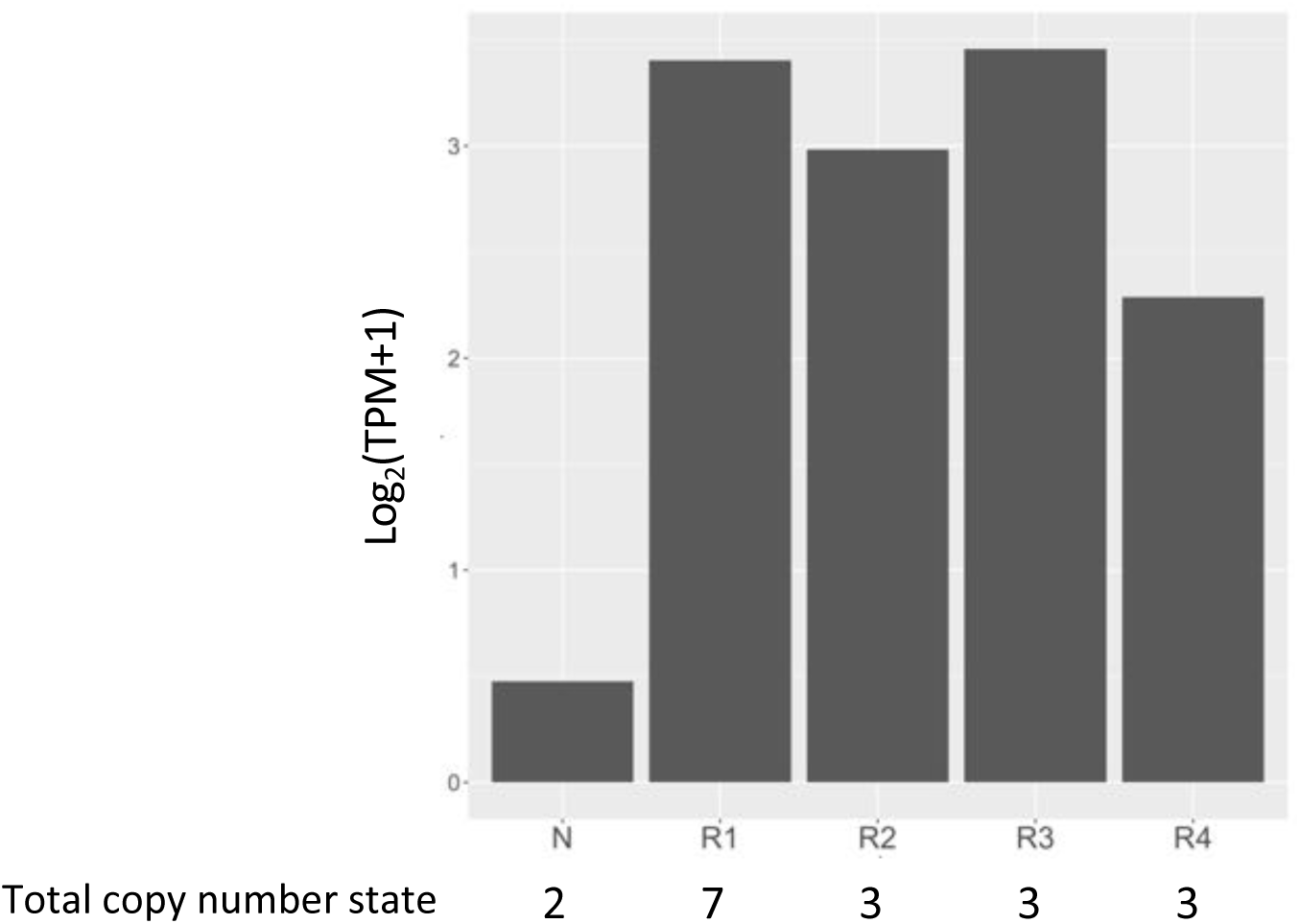
*SOX2* expression (log2(TPM+1)) in different tumor regions and benign tissue of sample P5. Bottom panel shows total copy number states inferred using FACETS. Benign tissue has copy number state = 2 (diploid). Tumor regions (R1, R2, R3, R4) show > 2 copy number depicting amplification in *SOX2* gene. Amplification is accompanied by increase in expression.

**Supplementary Figure 8:**
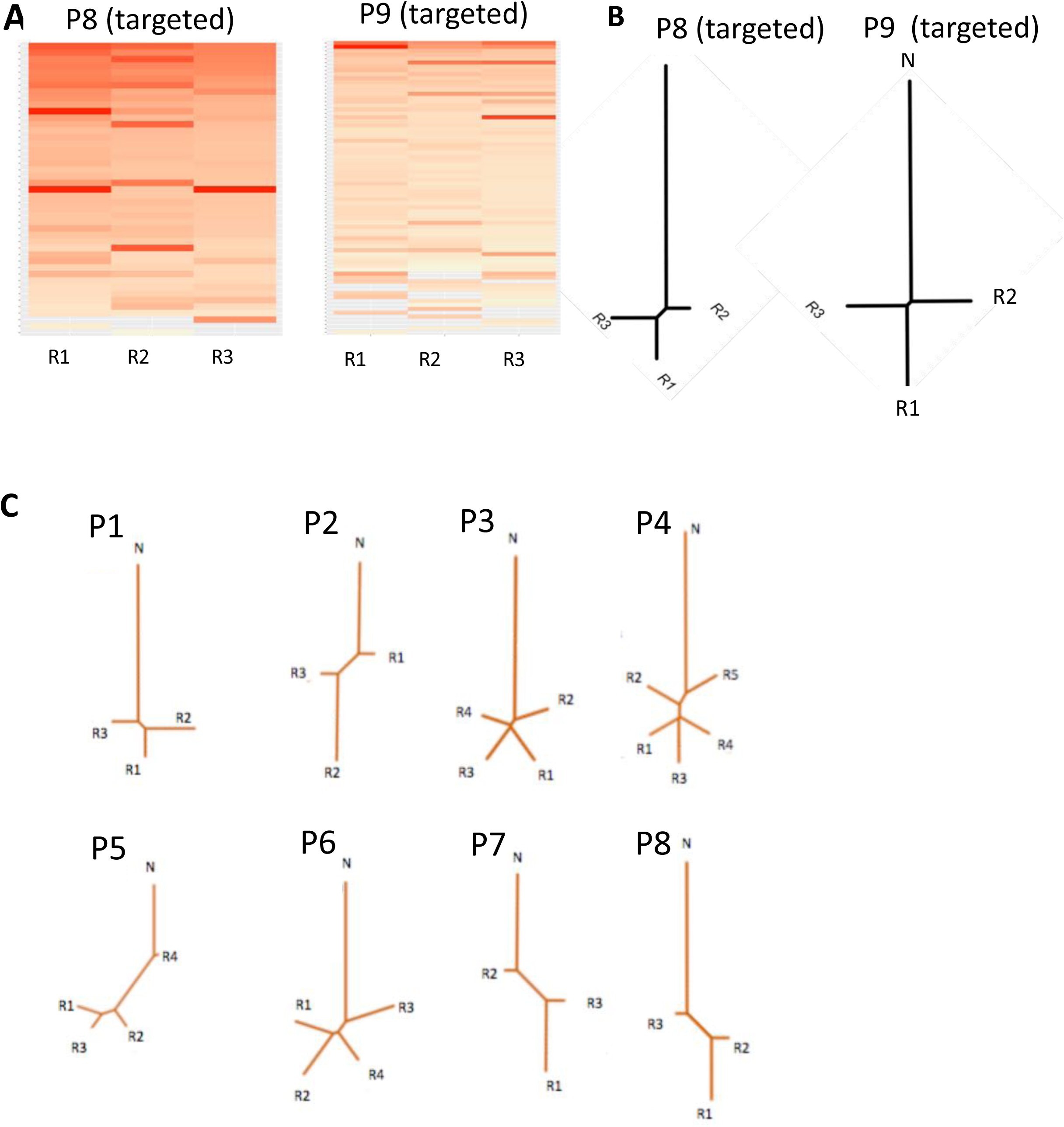
Genetic heterogeneity estimates from targeted sequencing data for two patients P8 and P9 of 257 genes. **A.** Heat maps showing regional variations in variant allele frequencies of single nucleotide somatic variations. Beige to red color gradient represents increasing variant allele frequency of somatic SNVs. Grey color represents absence of variation in a particular region. **B.** Dendrograms depicting distances between different tumor regions and normal inferred using somatic single nucleotide variations from targeted sequencing data. **C.** Purity-uncorrected dendrograms for samples P1-P8 based on exome sequencing data.

**Supplementary Figure 9:**
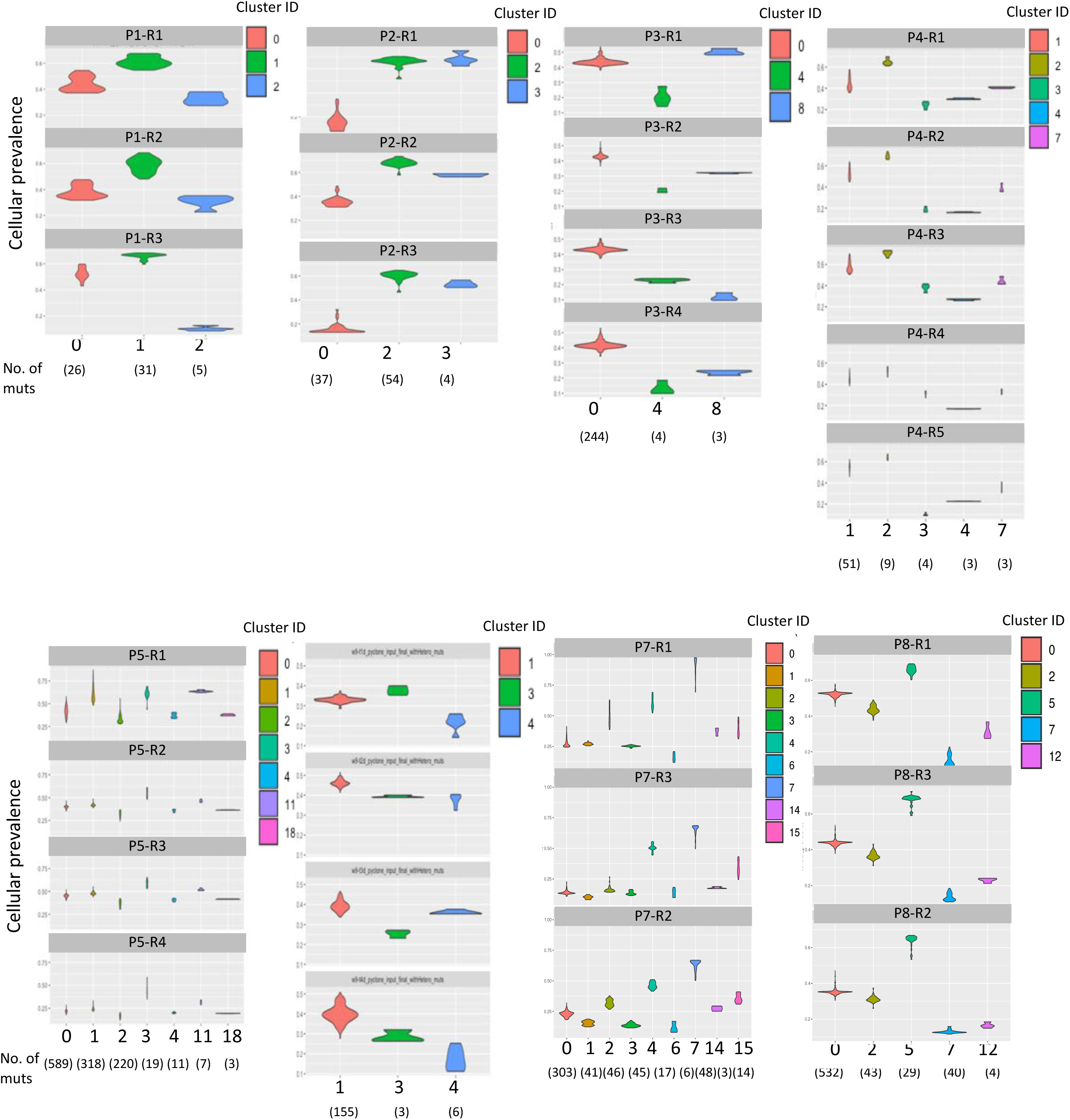
Inference about somatic mutation clusters and their proportional contributions in different tumor regions using PyClone. Somatic mutation clusters may serve as a proxy for subclonal entities, but it remains a computationally challenging problem and caution is warranted while inferring clonal architecture from mutation clusters. Different colors represent different mutational clusters and numbers in bracket below show number of mutations in each cluster. Any cluster with 3 or more mutations are plotted. Most clusters have very low number of mutations and hence cannot be reliably called as subclone. Overall there are 1-2 subclones across all samples with P5 as an exception with 3 clusters composed of considerable number of mutations. This observation is also supported by analysis done using orthogonal approach in Supplementary Figure 10.

**Supplementary Figure 10:**
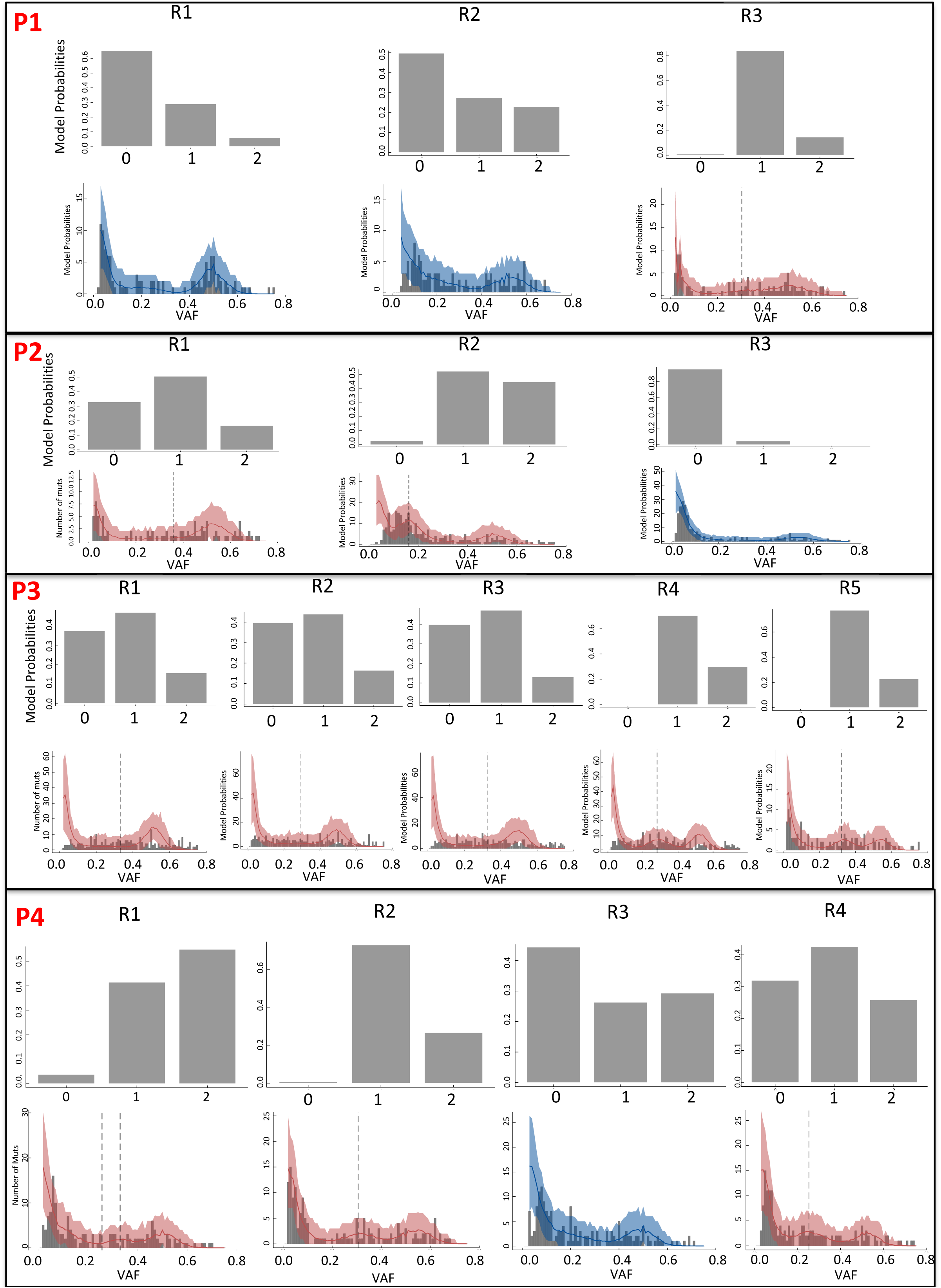
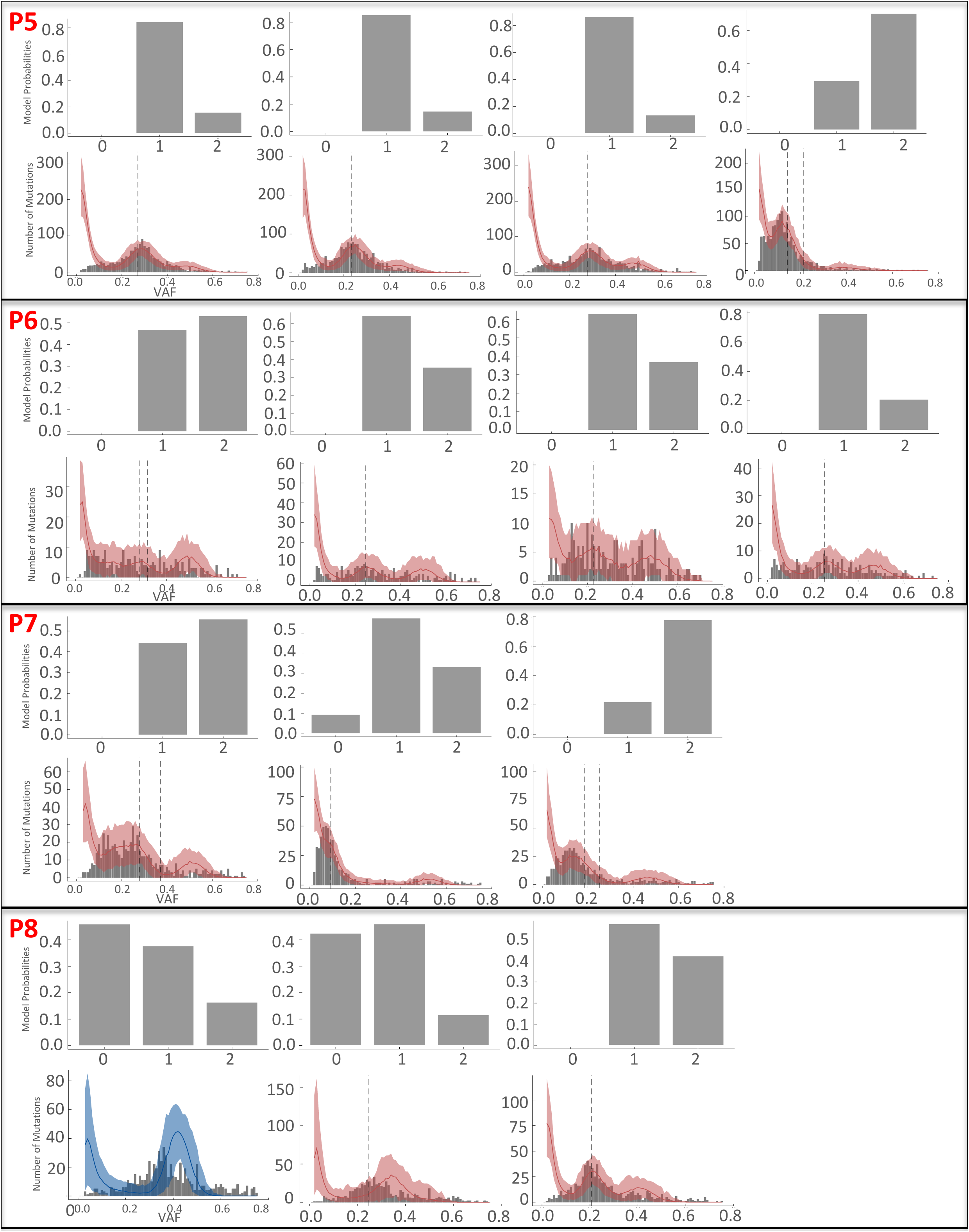
Clonal architecture inferred using Williams et al. approach. Upper panel of bar plots for all samples represent model probabilities of having different number of clones. X-axis is number of clones and Y-axis represents model probabilities. Lower panel of all samples show variant allele frequency distribution of all somatic SNVs. Dotted lines represent cut off for clusters. Two lines are for two subclones (model probability of having 2 subcloes is highest) and one line for one subclone (model probability of having one subclone is highest). For most of the samples number of subclones was 1-2, as shown by PyClone analysis also (Supplementary Figure 9). Overall, there is less heterogeneity in LUSC

**Supplementary Figure 11:**
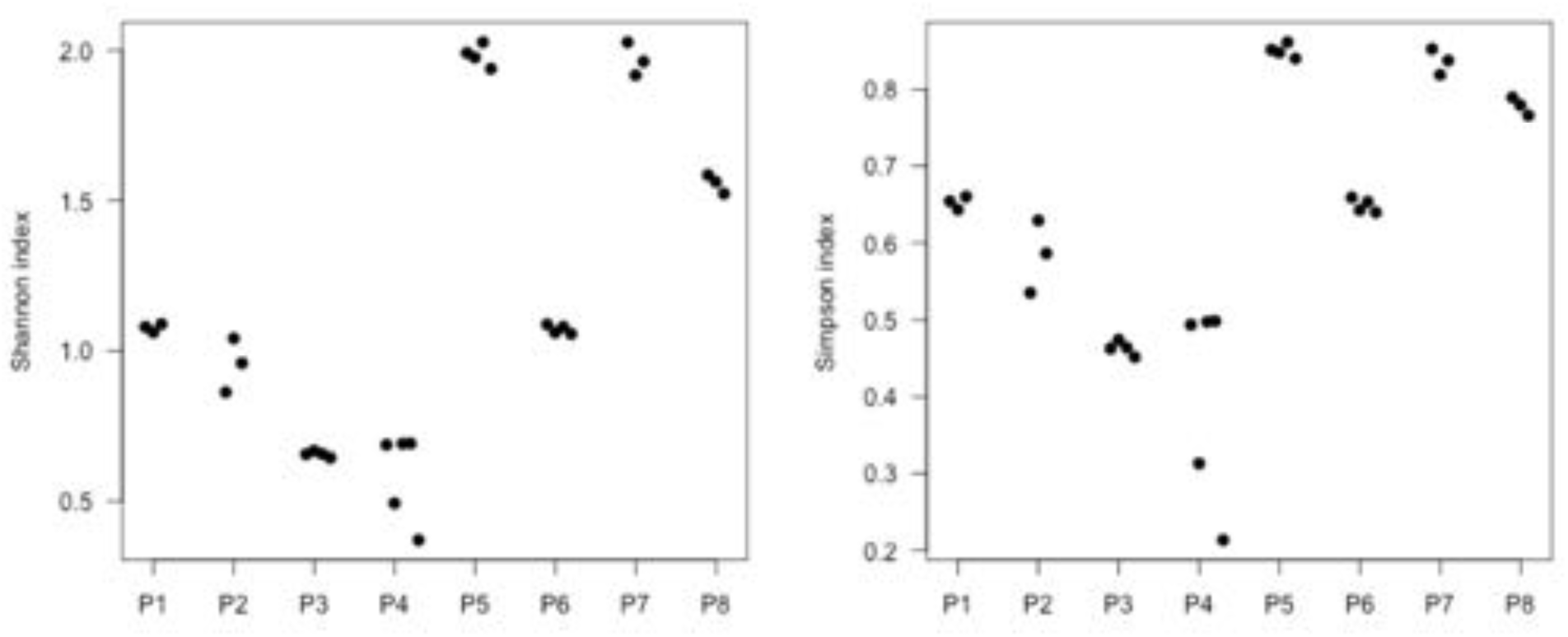
Shannon’s diversity index 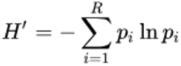 and Gini-Simpson’s index 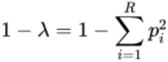 for each tumor region based on the estimated abundance of the PyClone mutation clusters. The values of these indices were very similar across the regions within a tumor. This under-appreciated the extent of between-region genetic differences, as reflected in the dendrogram analysis.

**Supplementary Figure 12:**
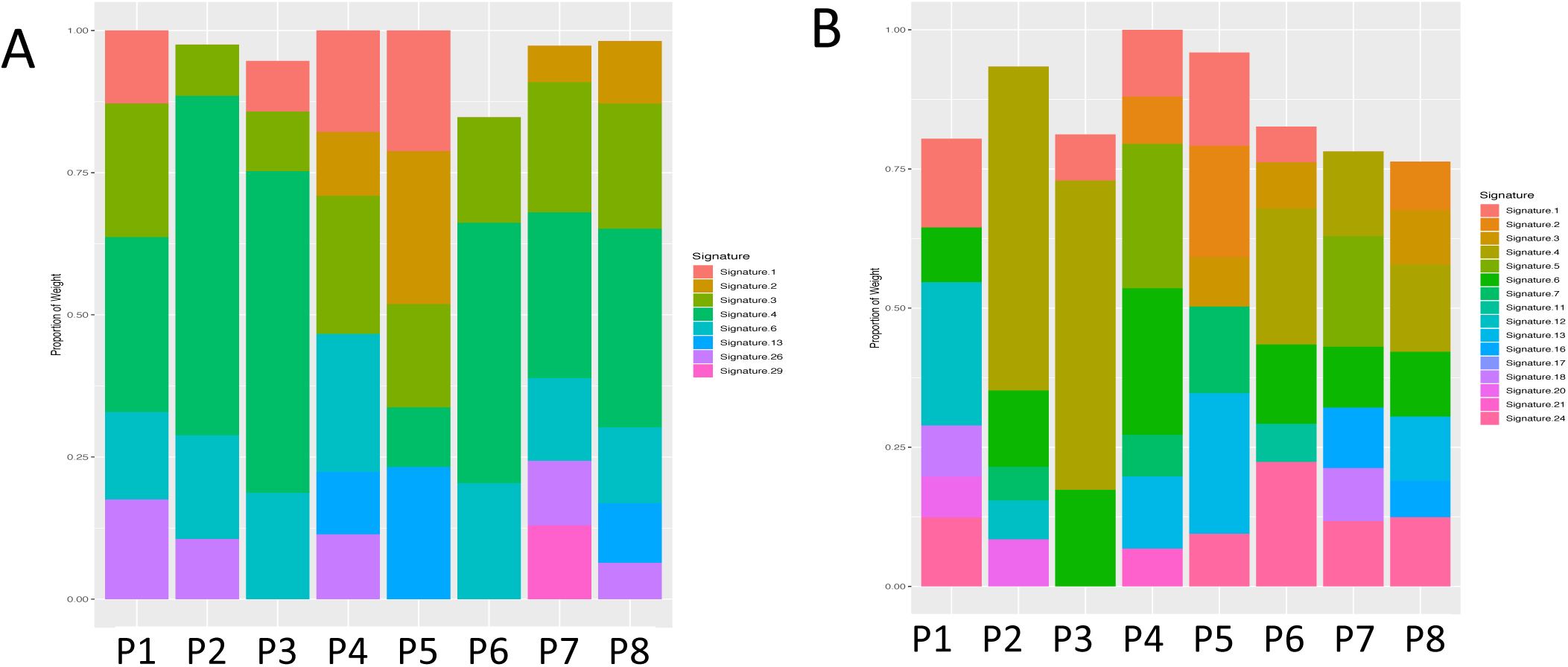
Stacked barplots showing contribution of **A)** selected mutational signatures relevant to lung cancer biology, and **B)** all mutational signatures from the COSMIC database - estimated from all somatic SNVs including ubiquitous and non-ubiquitous variants. Signature.1: Mutational process initiated by spontaneous deamination of 5-methyl cytosine, Signature.2: Mutational process due to APOBEC activity, Signature.3: Mutational process due to HR defect, Signature.4: Mutational process due to smoking, Signature.6: Mutational process due to defective DNA mismatch repair, Signature.13: Mutational process due to APOBEC activity, Signature.15: Mutational process due to defective DNA mismatch repair, Signature.26: Mutational process due to defective DNA mismatch repair, Signature.29: Mutational process due to tobacco chewing habit.

**Supplementary Figure 13:**
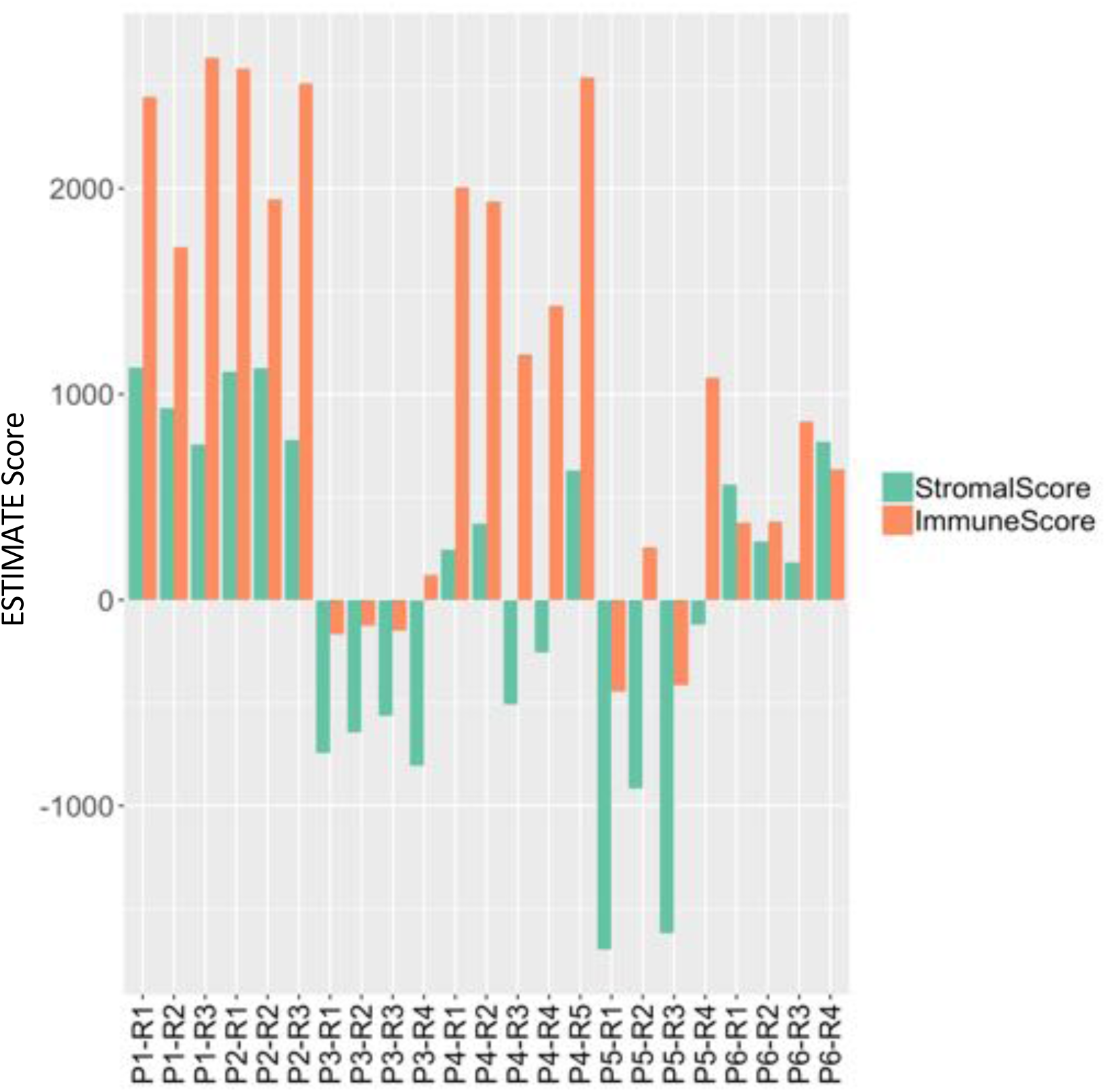
Patterns of immune score and stromal score estimated from RNAseq-based expression data using ESTIMATE algorithm indicates the extent of immune and stromal cell infiltration in different regions in the tumors for P1-P6 patients.

**Supplementary Figure 14:**
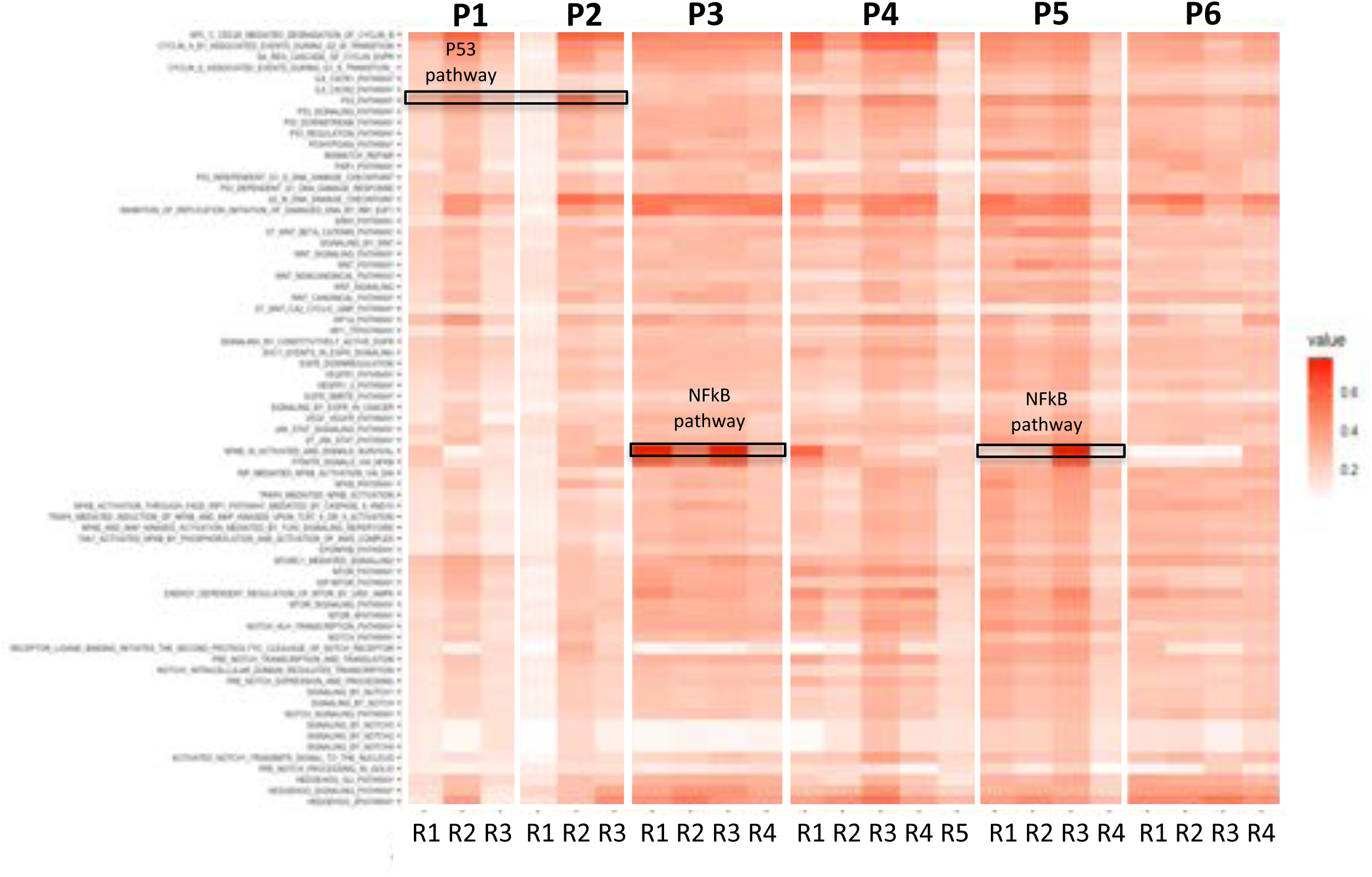
Extent of pathway deregulation calculated using nJSD, an entropy based distance metric between lung squamous carcinoma associated networks in tumor regions compared to matched benign tissues (P1-P6). *TP53* and *NFKB* pathways that show high between-region differences within a tumor are highlighted as examples.

**Supplementary Figure 15:**
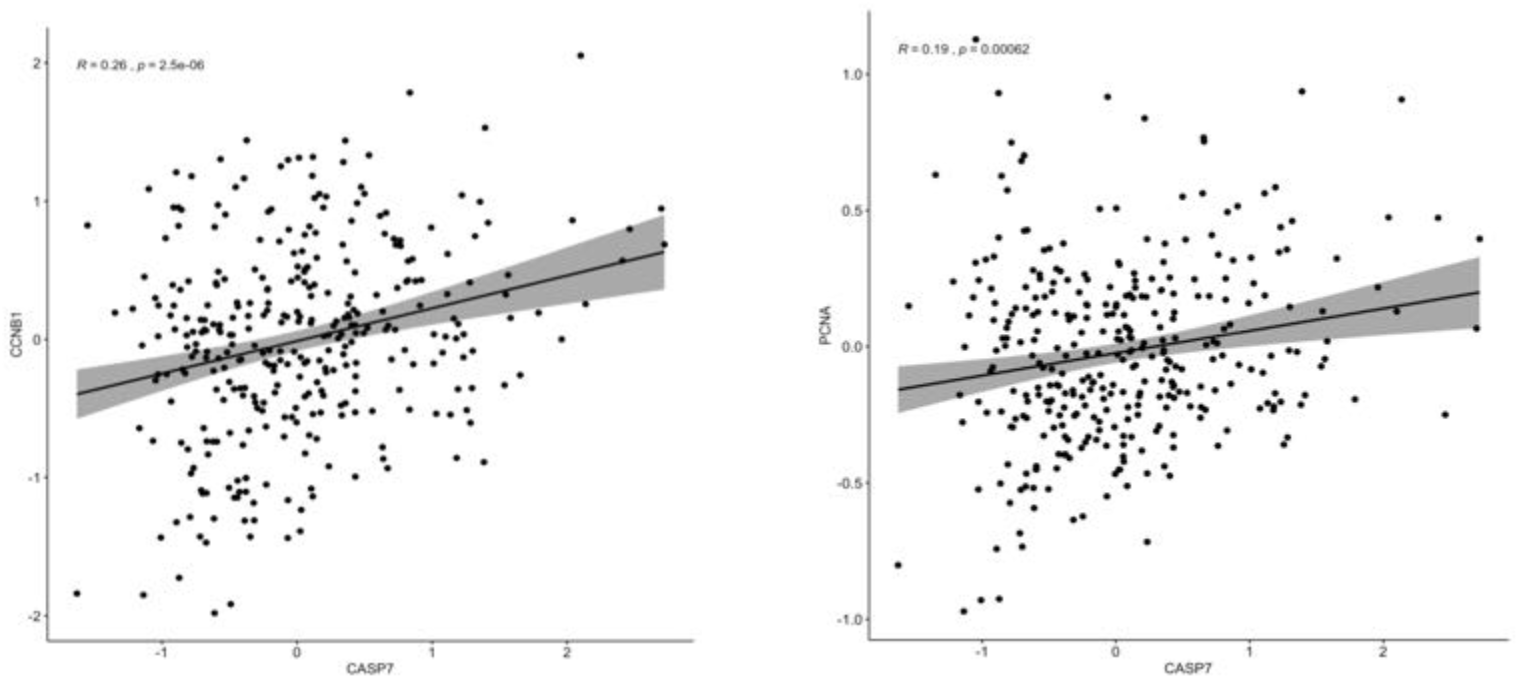
Correlation between **A)** proliferation associated gene *CCNB1* and apoptosis associated gene *CASP7* protein expression (RPPA data), and **B)** proliferation associated gene *PCNA* and apoptosis associated gene *CASP7* protein expression (RPPA data) the TCGA Lung squamous cell carcinoma cohort. Spearman Correlation coefficient and associated p-values are shown in top left corner.

**Supplementary Figure 16:**
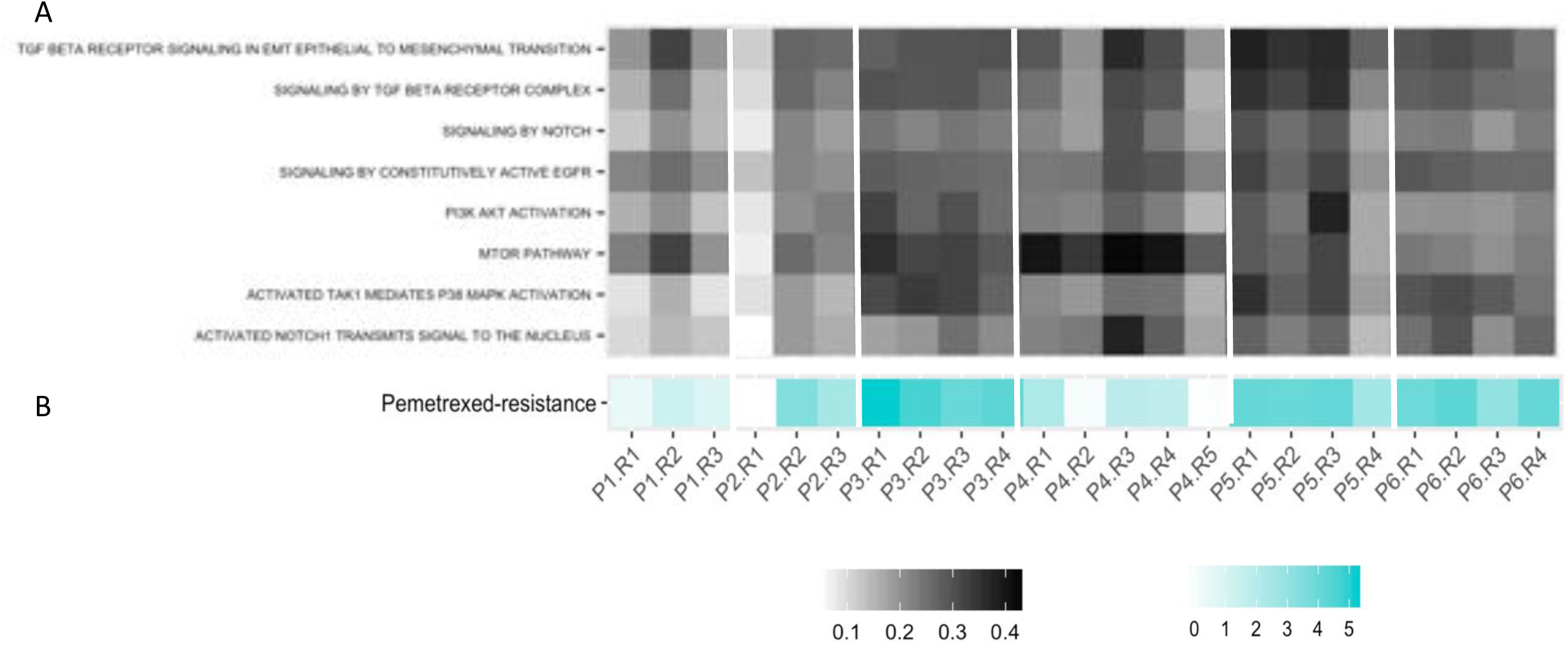
Drug-resistance scores **A)** Pathway deregulation scores (calculated using nJSD) for lung cancer associated multi-drug resistance associated pathways. Darker black color represents higher deregulation. **B)** Pemetrexed resistance score calculated using 25 gene expression signature. Higher expression indicates higher resistance.

**Supplementary Figure 17:**
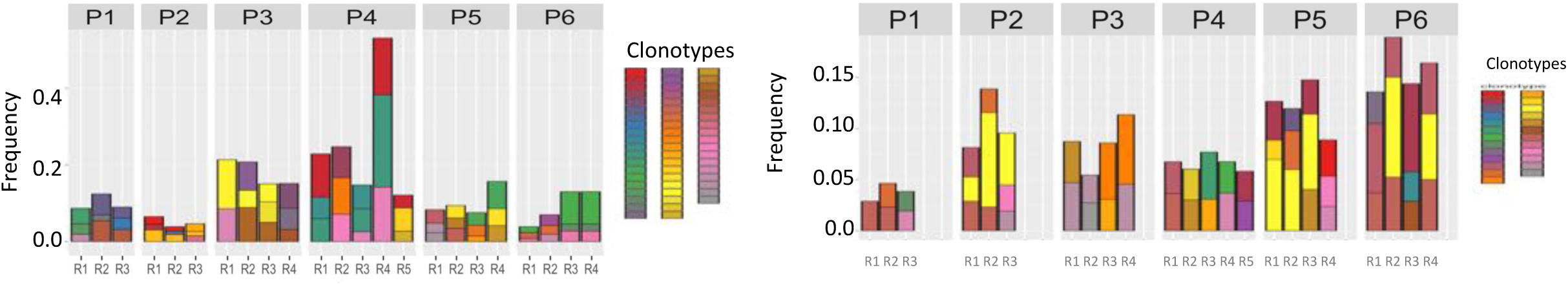
TCR repertoire heterogeneity inferred using Mixcr tool. A) TCR clonotypes based on TCR status inferred from RNAseq data in different regions of all samples. Each color represent one clonotypes in one sample. Some clones are shared in different regions B) TCR clonotypes inferred from exome sequencing data in different tumor regions. Each color represent one clone in one sample. Some clonotypes are shared in different regions. In both data, clonotypes between different tumor specimens are not comparable. No overlap was observed between clonotypes inferred from DNA and RNA-level data.

**Supplementary Figure 18:**
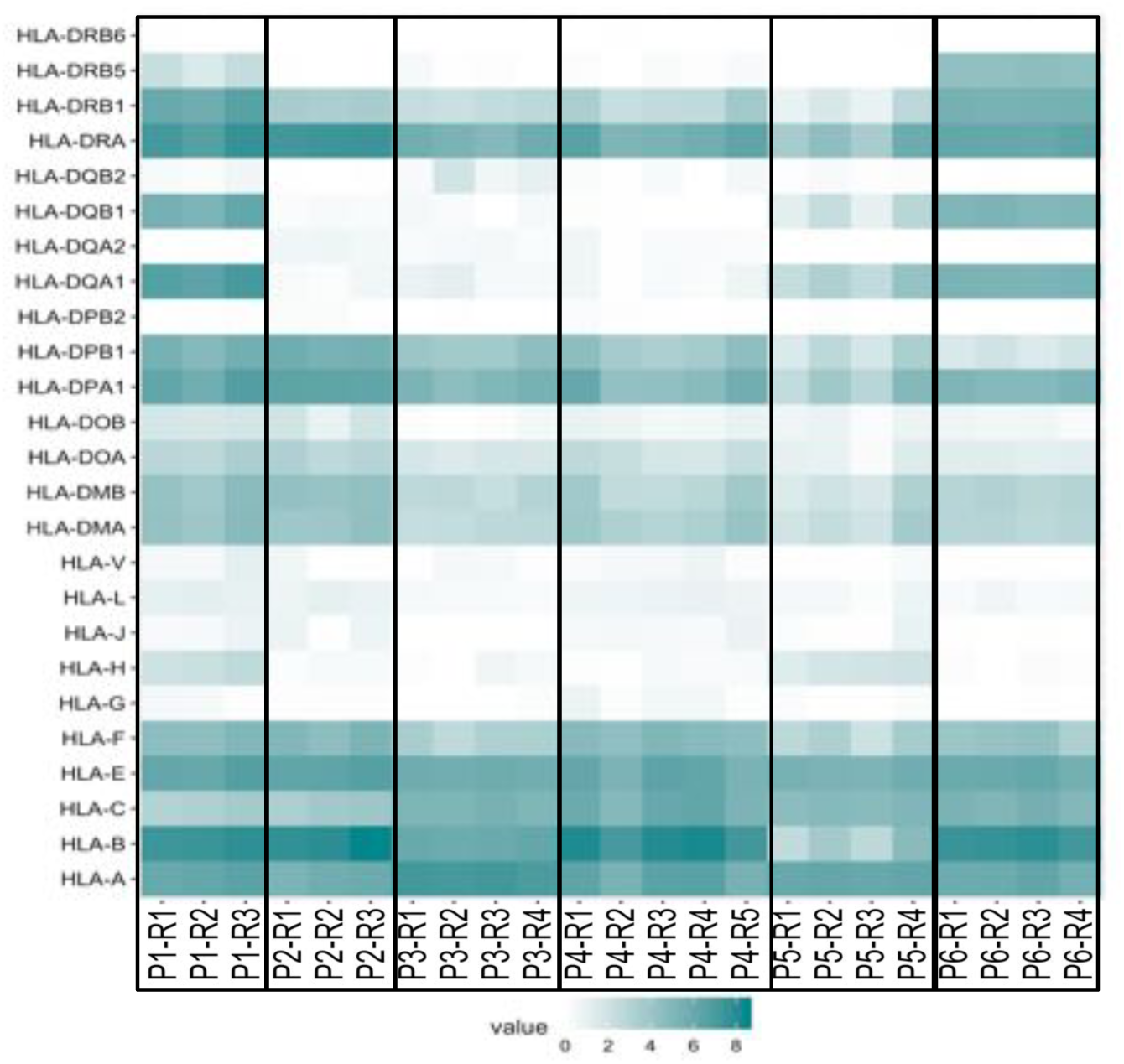
Patterns log2(TPM+1) expression values for HLA genes in different tumor regions for P1-P6 samples.

**Supplementary Figure 19:**
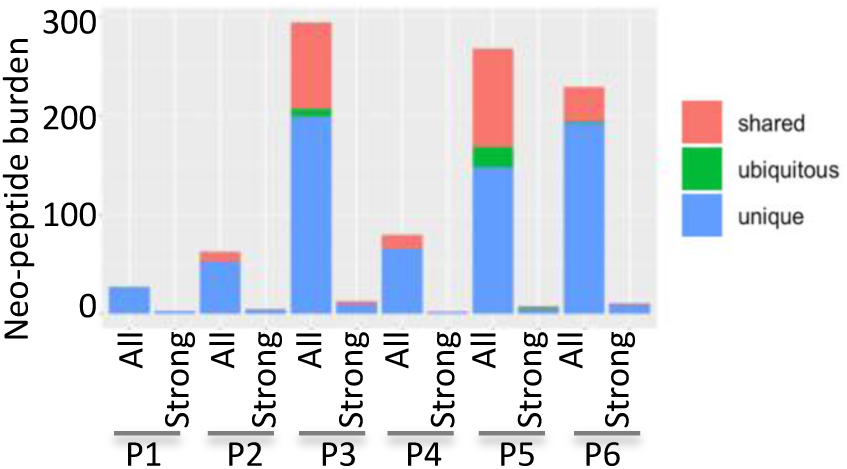
Stacked bar chart representing estimated neo-epitope burden that are (i) ubiquitous i.e. present in all regions in a tumor sample (ii) shared i.e. present in multiple but not all regions, and (iii) unique i.e. present only in one region profiled in a tumor. For each patient the first stacked bar shows all neo-epitopes, and the second one shows the pattern for neo-epitopes that are suspected to have strong immunogenic effects.

**Supplementary Figure 20:**
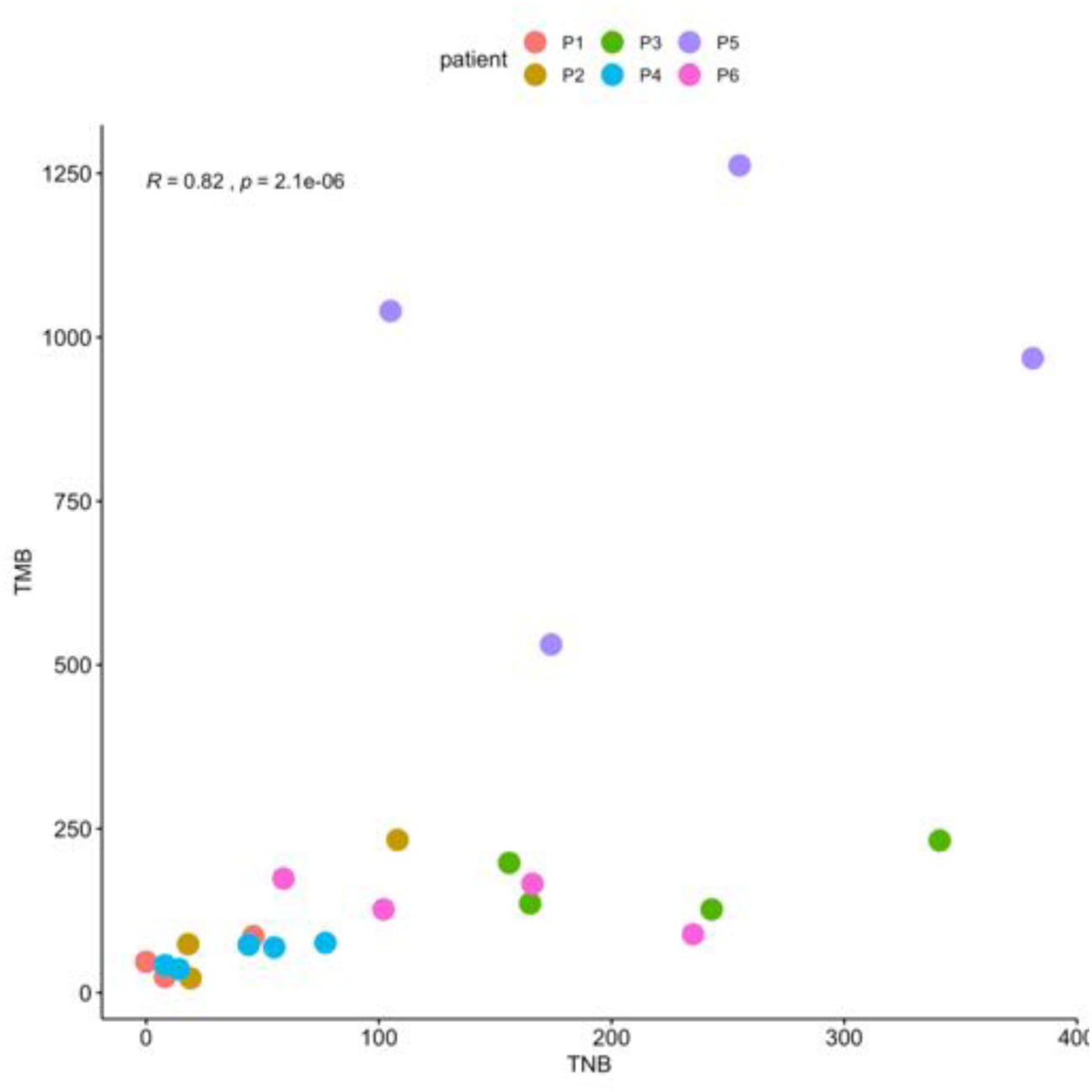
Scatter plot showing correlation (spearman correlation) between Tumor Mutation Burden (TMB) and Tumor Neo-epitope Burden (TNB) across all regions of all tumor samples (P1-P6). Different colors represent different samples. Correlation coefficient and p value are depicted on top left of the plot.

**Supplementary Figure 21:**
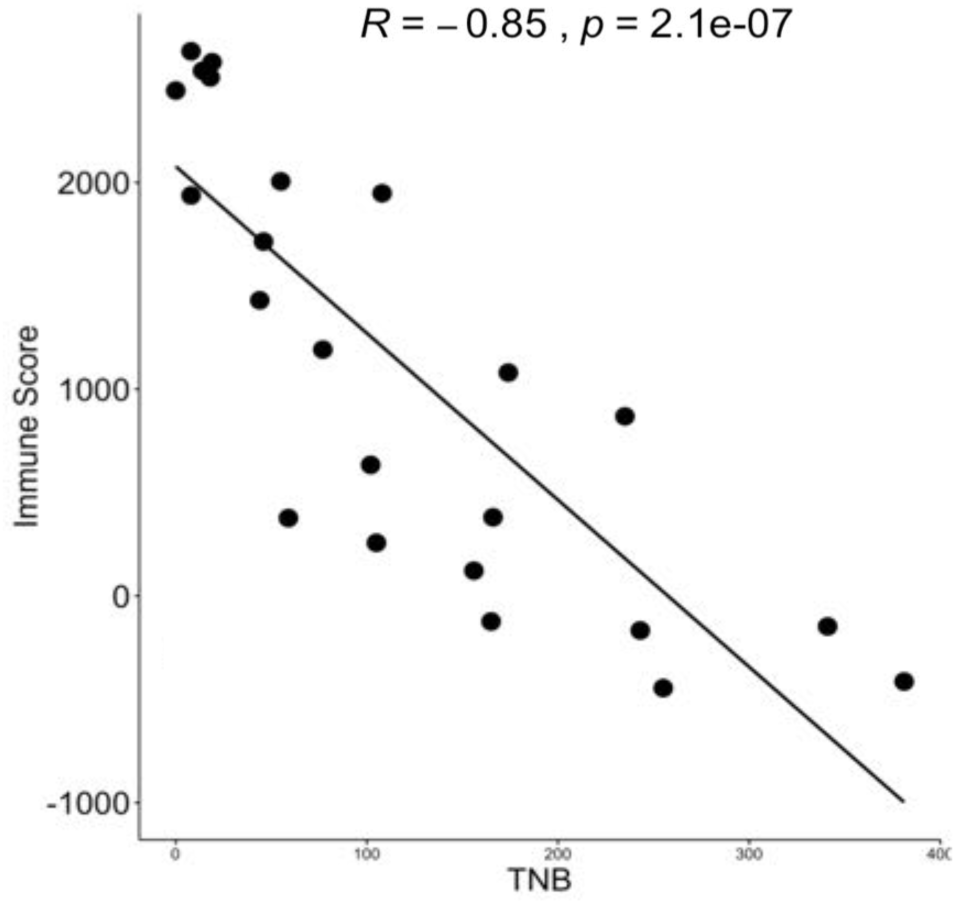
Scatter plot showing correlation (spearman correlation) between Tumor Neo-epitope Burden (TNB) and immune score (representing extent of immune infiltration calculated using ESTIMATE) across all regions of all tumor samples (P1-P6). Correlation coefficient and p value are depicted on top of the plot.

**Supplementary Figure 22:**
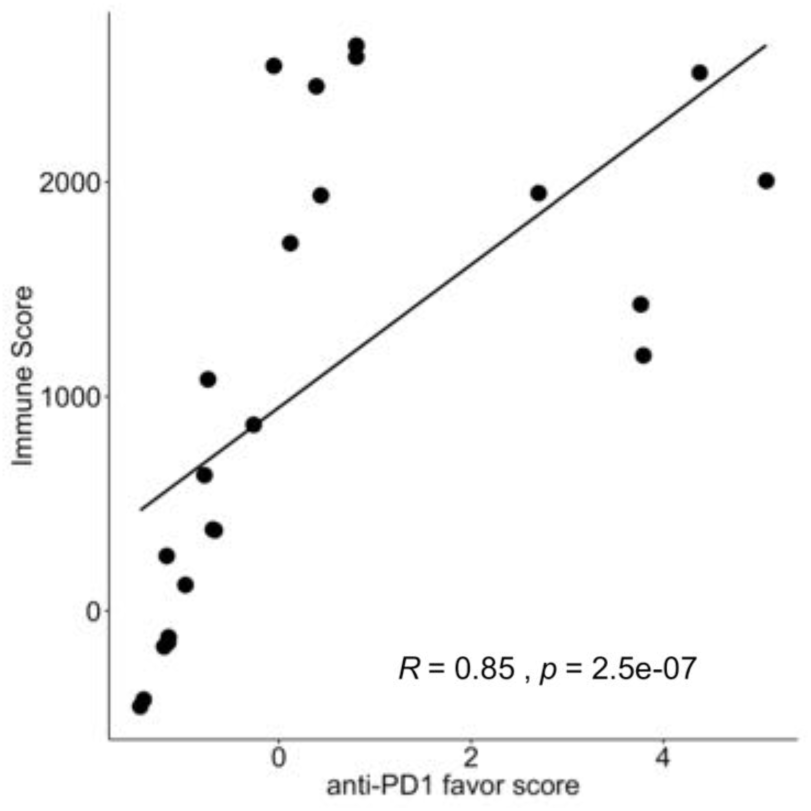
Scatter plot showing correlation (spearman correlation) between anti-PD1 favor score and Immune score (representing extent of immune infiltration calculated using ESTIMATE) across all regions of all tumor samples (P1-P6). Correlation coefficient and p value are depicted on the bottom of the plot.

**Supplementary Figure 23:**
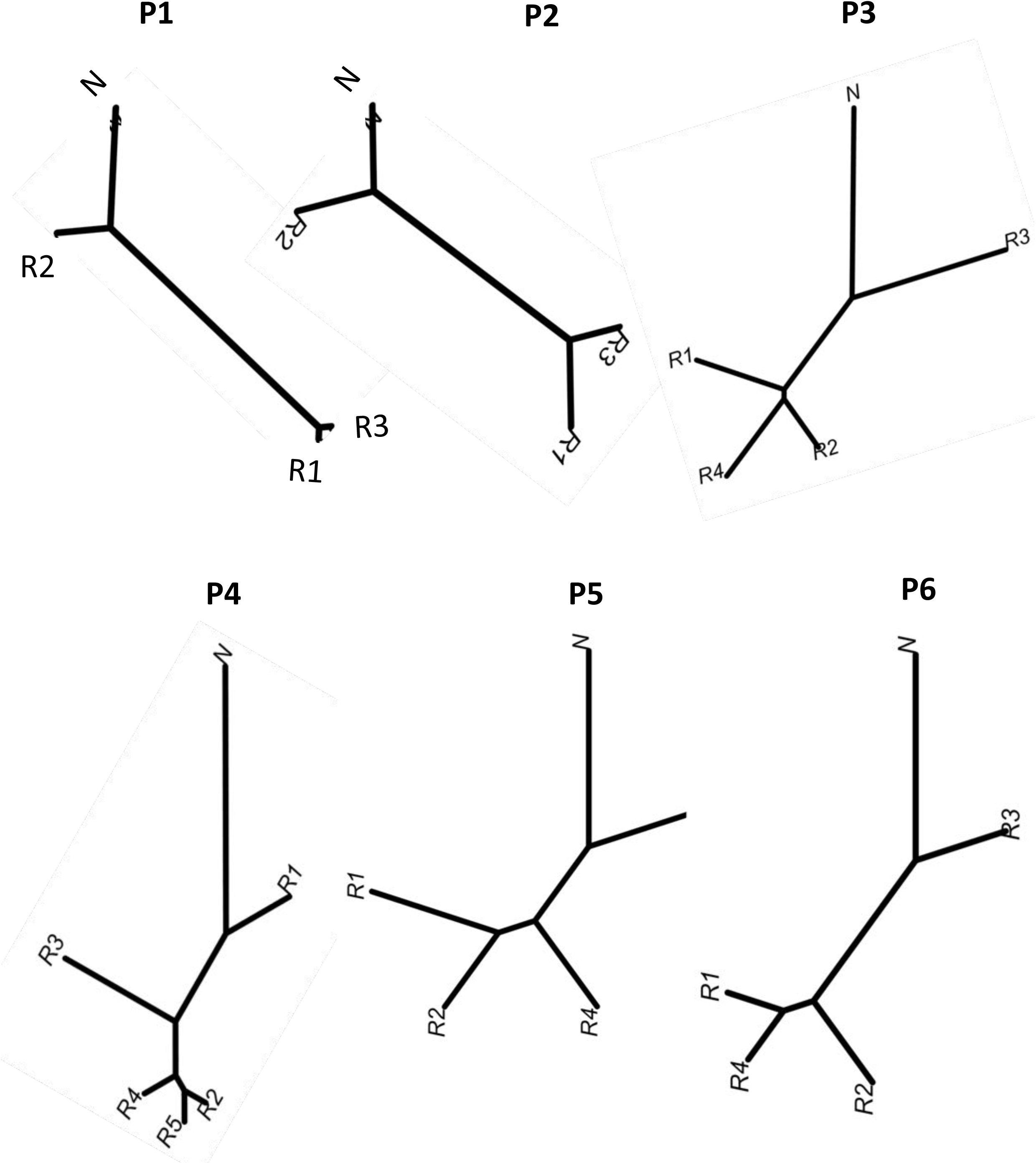
Dendrograms represent immunogenic similarity between different regions within a tumor for each patient sample based on neo-epitopes. The distances used to make dendrograms are derived from neo-epitope abundance and expression in different regions of a tumor.

**Supplementary Figure 24:**
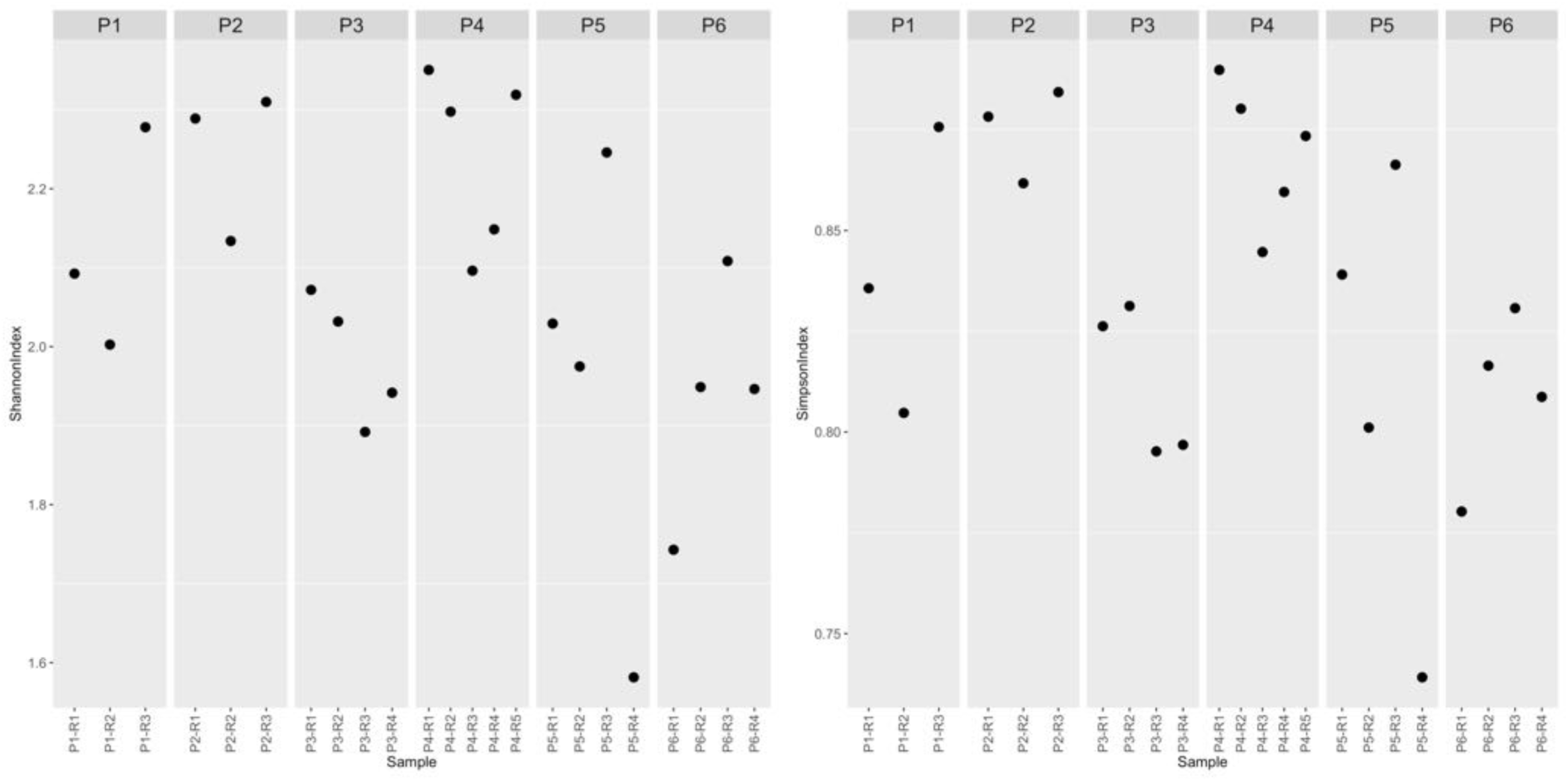
Shannon’s diversity index 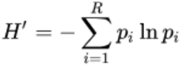 and Gini-Simpson’s index 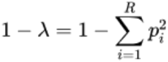 for each tumor region based on the estimated abundance of the Immune cell populations (CIBERSORT). Unlike genomic data the values of these indices were different across the regions within a tumor indicating more immunogenic heterogeneity as compared to genetic.

**Supplementary Figure 25:**
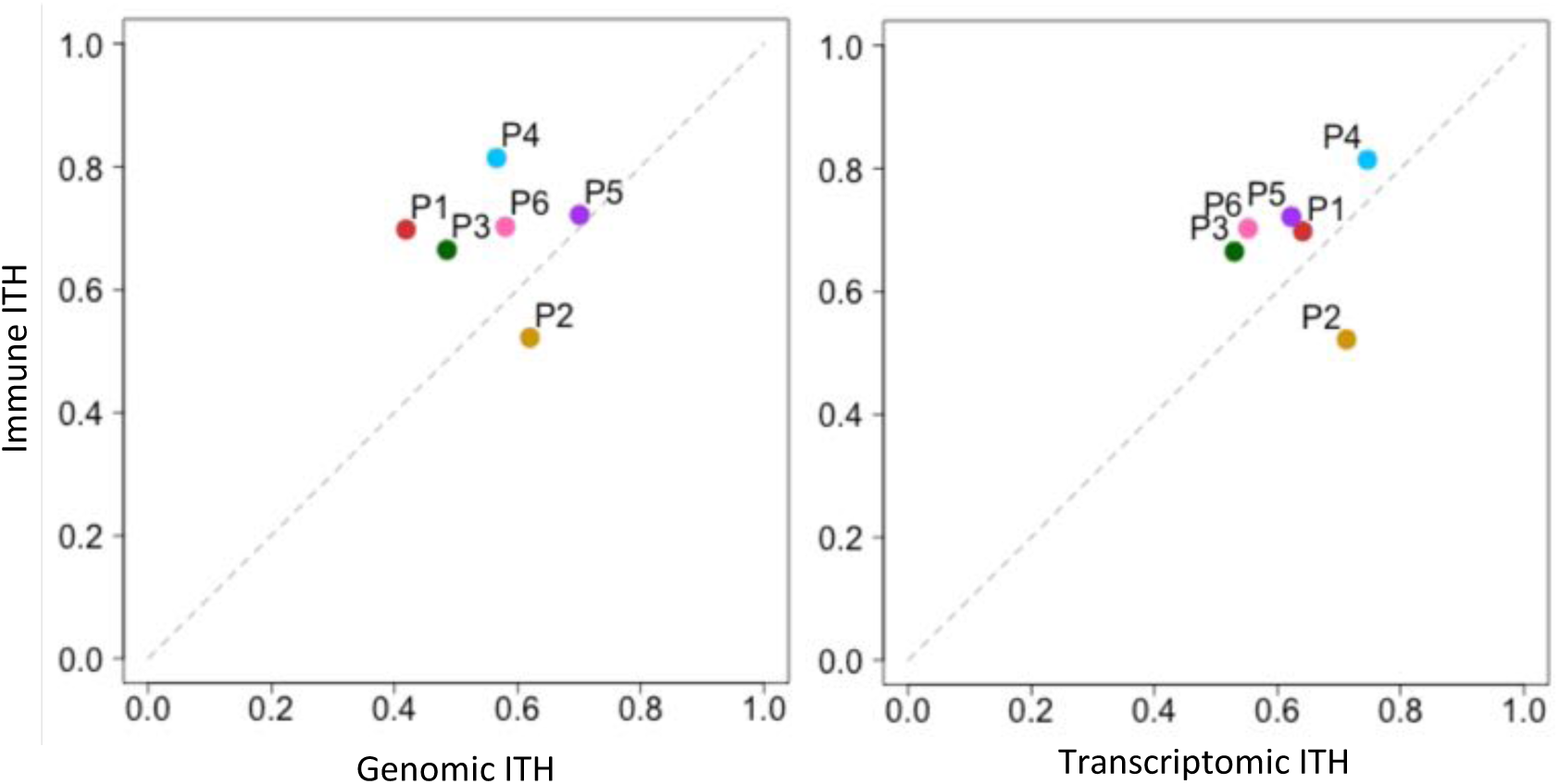
Scatterplot comparing the extent of immune and genetic divergence (left panel), and the extent of immune and transcriptomic divergence (right panel) for different tumor regions from their respective matched benign tissues. Color codes are same as Figure 2A.

**Supplementary Figure 26:**
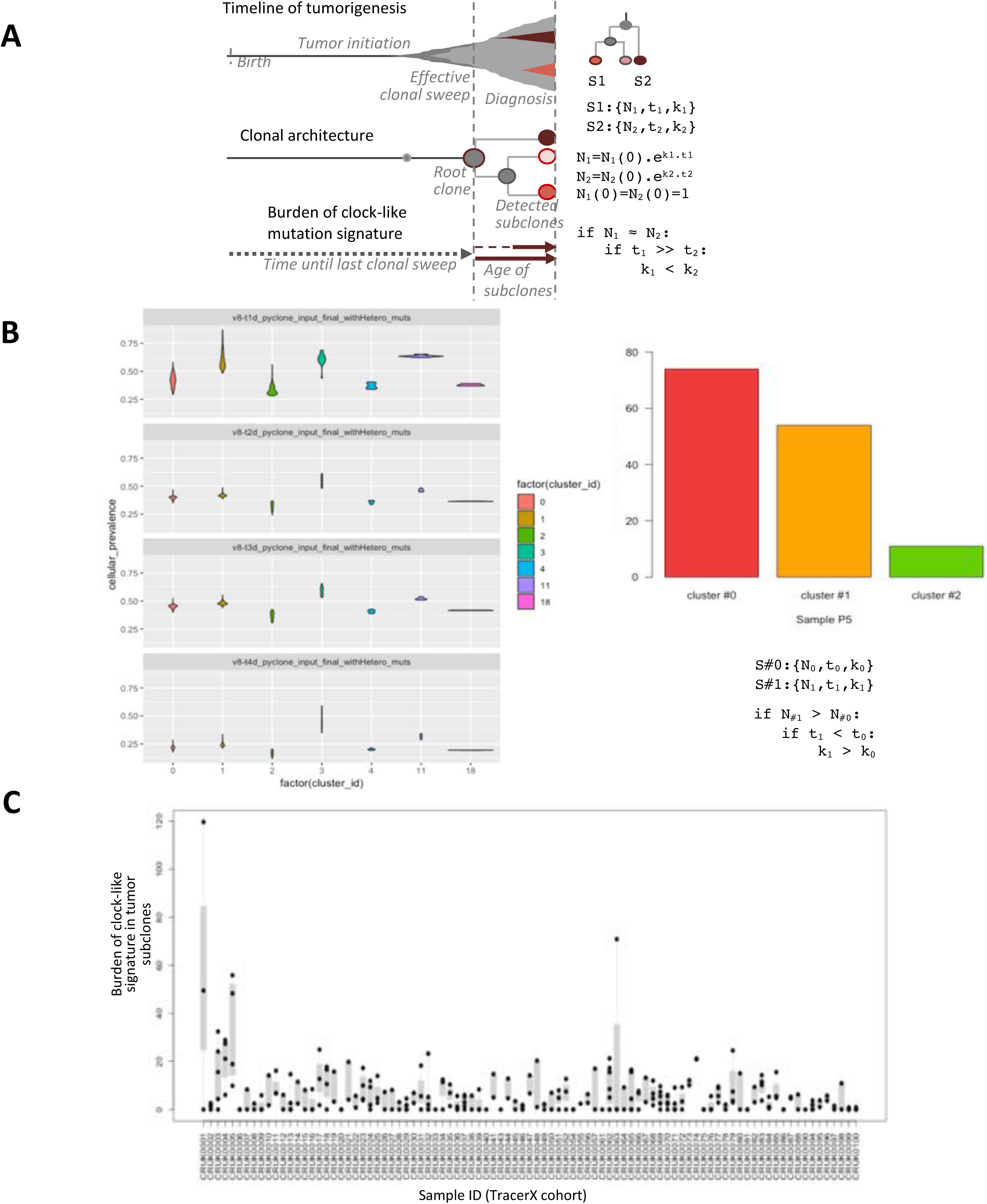
**A)** Inference about subclonal age and growth rates based on the burden of clock-like signature in somatic mutations private to subclones detected in a tumor. Effective clonal sweep indicates the point where the subclone becomes highly dominant within the tumor, within the resolution of genomic detection, such that it appears to be present at almost 100% cell fraction. **B)** In Sample P5, mutation clusters were identified using PyClone as shown in Supplementary Figure 9. The burden of clock-like signature was estimated for each mutation cluster, and plotted for cluster #0 – 2, which had sufficient mutations. Not all clusters in all samples had sufficient somatic mutations for signature analysis. **C**) Scatterplot showing the estimated burden of clock-like signatures in tumor subclones for the patients in the TracerX cohort, plotted for different clones of all TracerX samples. Each dot represents different subclones of a patient. Some patients have substantial variations in the burden of clock-like signatures among tumor subclones.

**Supplementary Figure 27:**
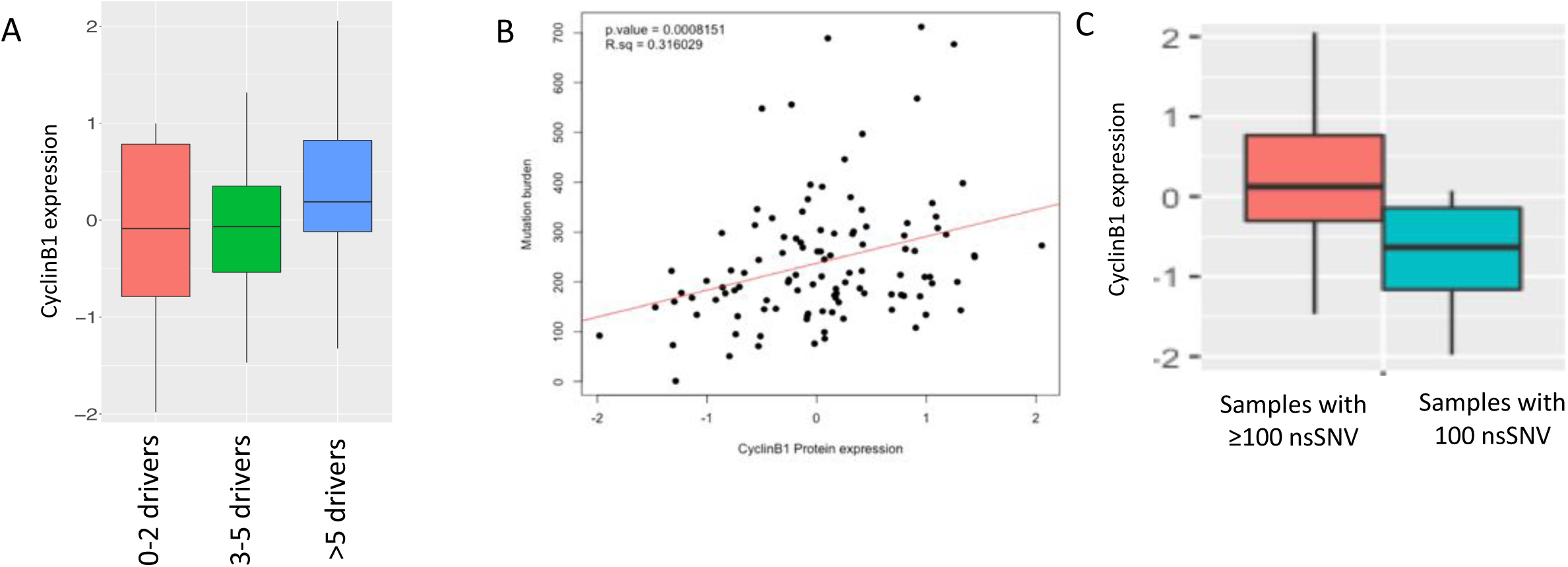
**A)** Cyclin B expression in TCGA lung squamous cell carcinoma (LUSC) tumors grouped by the number of known oncogenic driver mutations. **B)** Correlation between somatic non-silent single nucleotide variant (SNV) burden and Cyclin B1 protein expression (RPPA data) in the TCGA Lung squamous cell carcinoma cohort. The regression line is marked in red, and Spearman correlation coefficient and p-value are mentioned in top left. **C)** Boxplot showing Cyclin B1 expression for the TCGA lung squamous cell carcinoma samples with at least 100 somatic SNVs, and those with less than 100 somatic SNVs. The difference is statistically significant (Mann Whitney U test; p-value < 0.01). Somatic mutation burden appear to correlate with proliferative potential of the tumors, because a vast majority of the somatic mutations arise during incorrect repair of genomic lesions during replication.

**Supplementary Figure 28:**
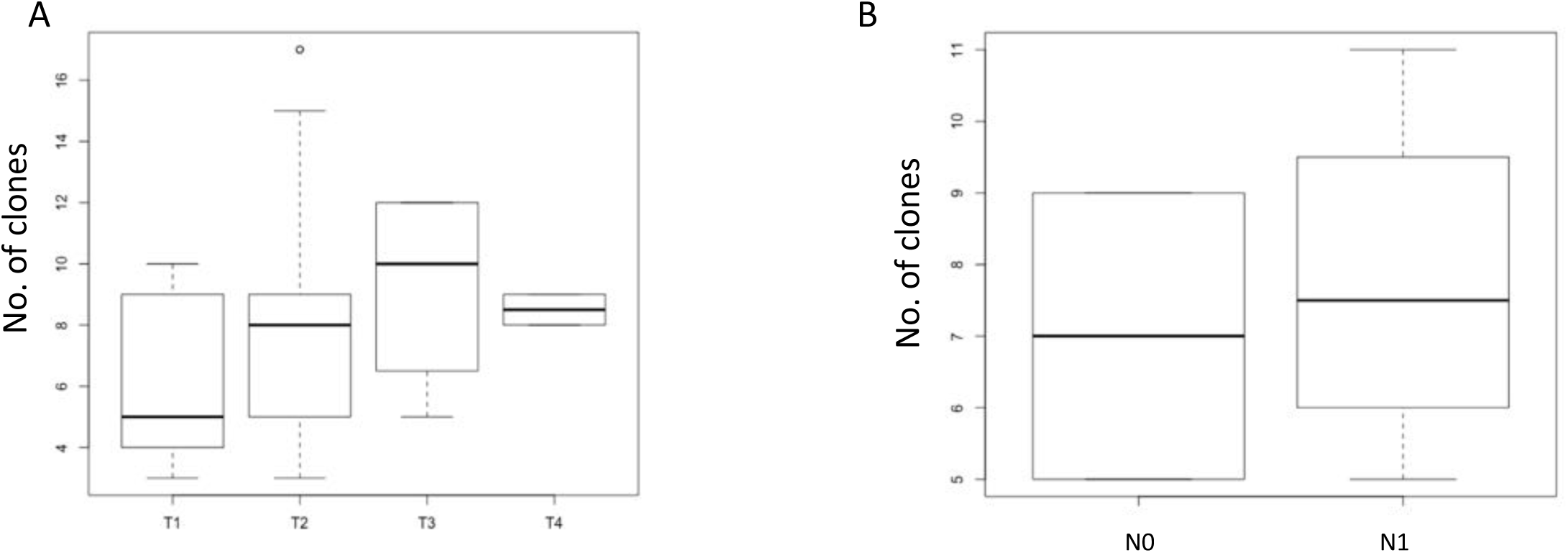
**A)** Boxplot showing number of clones in different T stages of LUSC tumor samples from TCGA data. There is a trend of increasing number of clones with higher stages, although the difference is not significance because of less number of samples in each category. **B)** Boxplot showing more number of clones in different N stages of LUSC samples from TCGA data. Like T stages, samples with lymph node invasion also show trend of more number of clones. This difference is also not significant because of less number of samples.

**Supplementary Figure 29:**
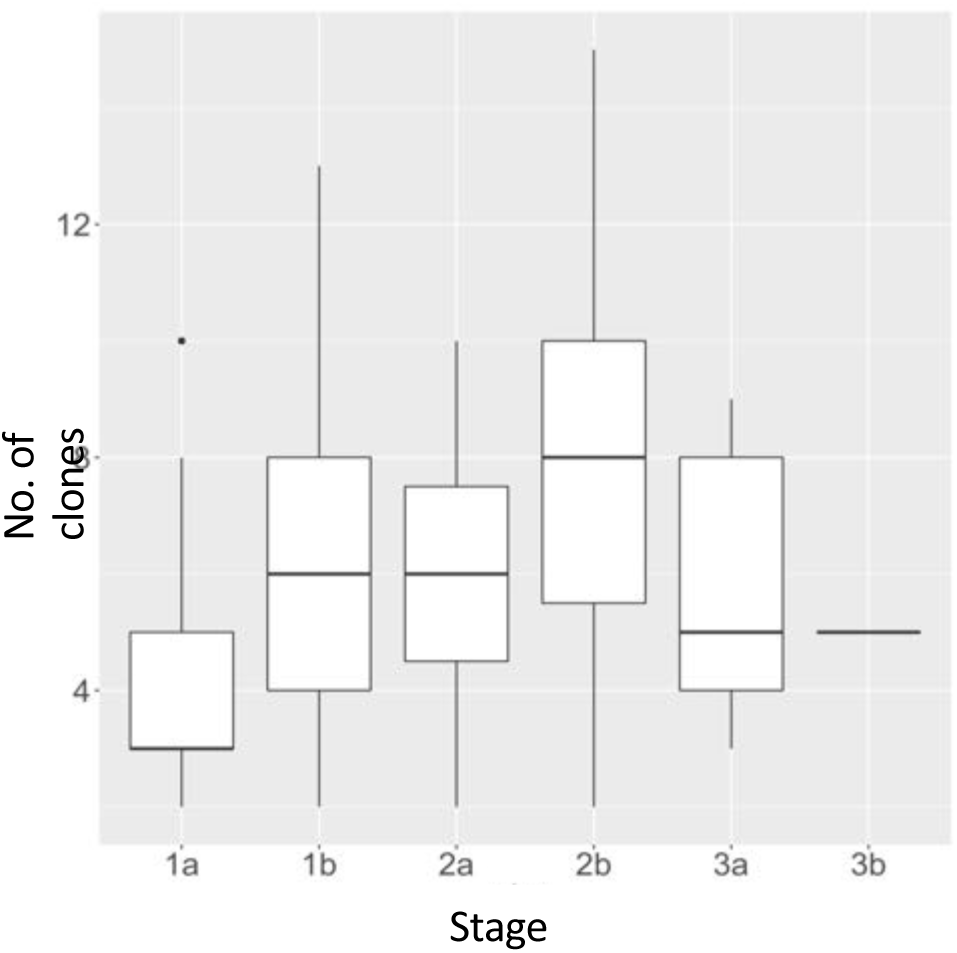
A) Boxplot showing number of clones in different stages of 91 lung tumor samples from TracerX data. There is a trend of increasing number of clones with higher stages, which reversed in Stage III. Stage 3b has only 1 sample.

**Supplementary Figure 30:**
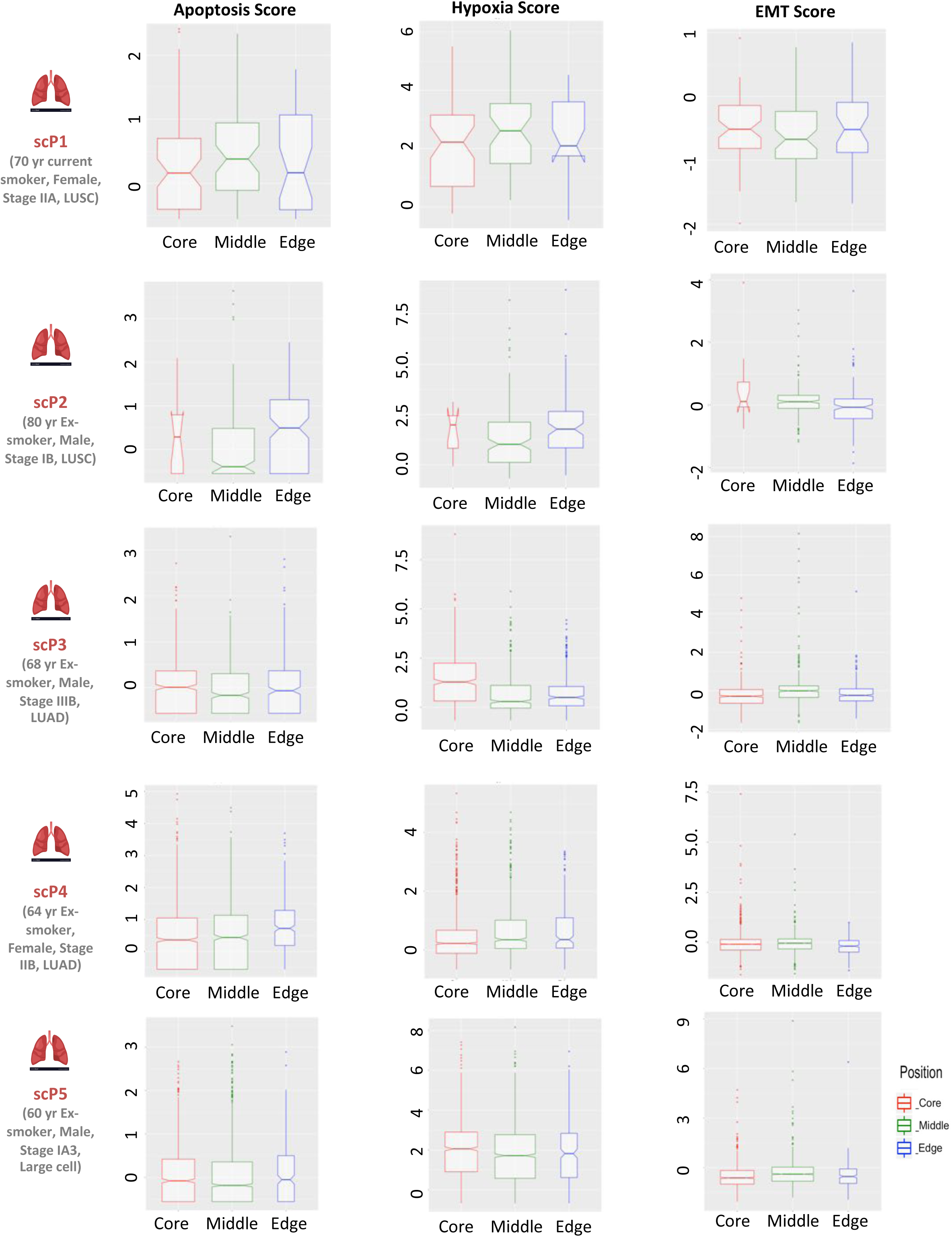
Boxplots showing distribution of apoptosis, hypoxia and EMT scores in different regions of tumors (Core, Middle and Edge). Scores were calculated from log2CPM data from scRNAseq using same approach as used for bulk RNAseq data (See Methods). Each row represents one patient. X-axis represents different clusters and y-axis represents apoptosis, hypoxia and EMT scores. Each point corresponds to single cell.

**Supplementary Figure 31:**
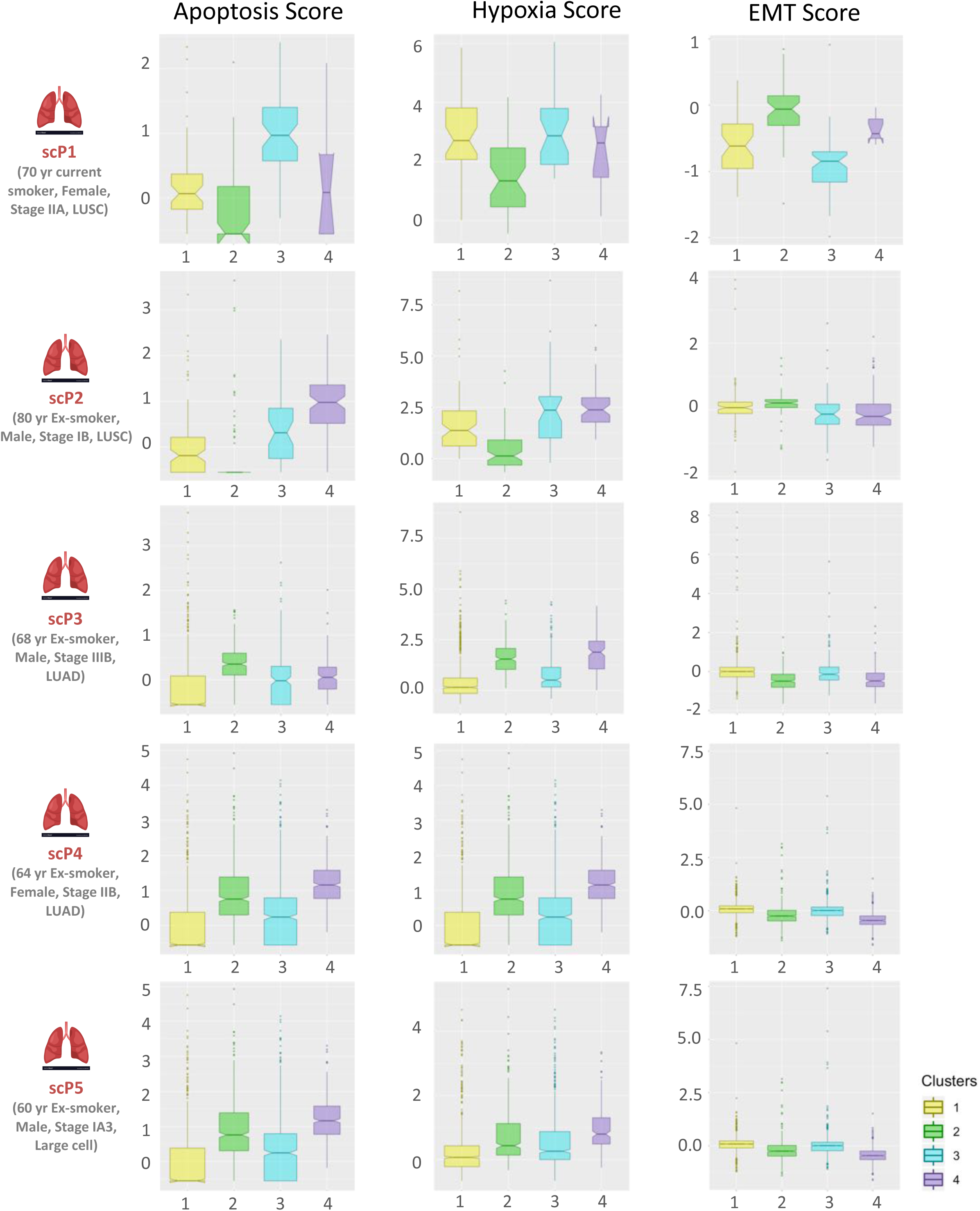
Boxplots showing distribution of apoptosis, hypoxia and EMT scores in different clusters of tumors. Each dot represents single cell from respective cluster. Scores were calculated from log2CPM data from scRNAseq using same approach as used for bulk RNAseq data (See Methods). Each row represents one patient. X-axis represents different clusters and y-axis represents apoptosis, hypoxia and EMT scores.

**Supplementary Figure 32:**
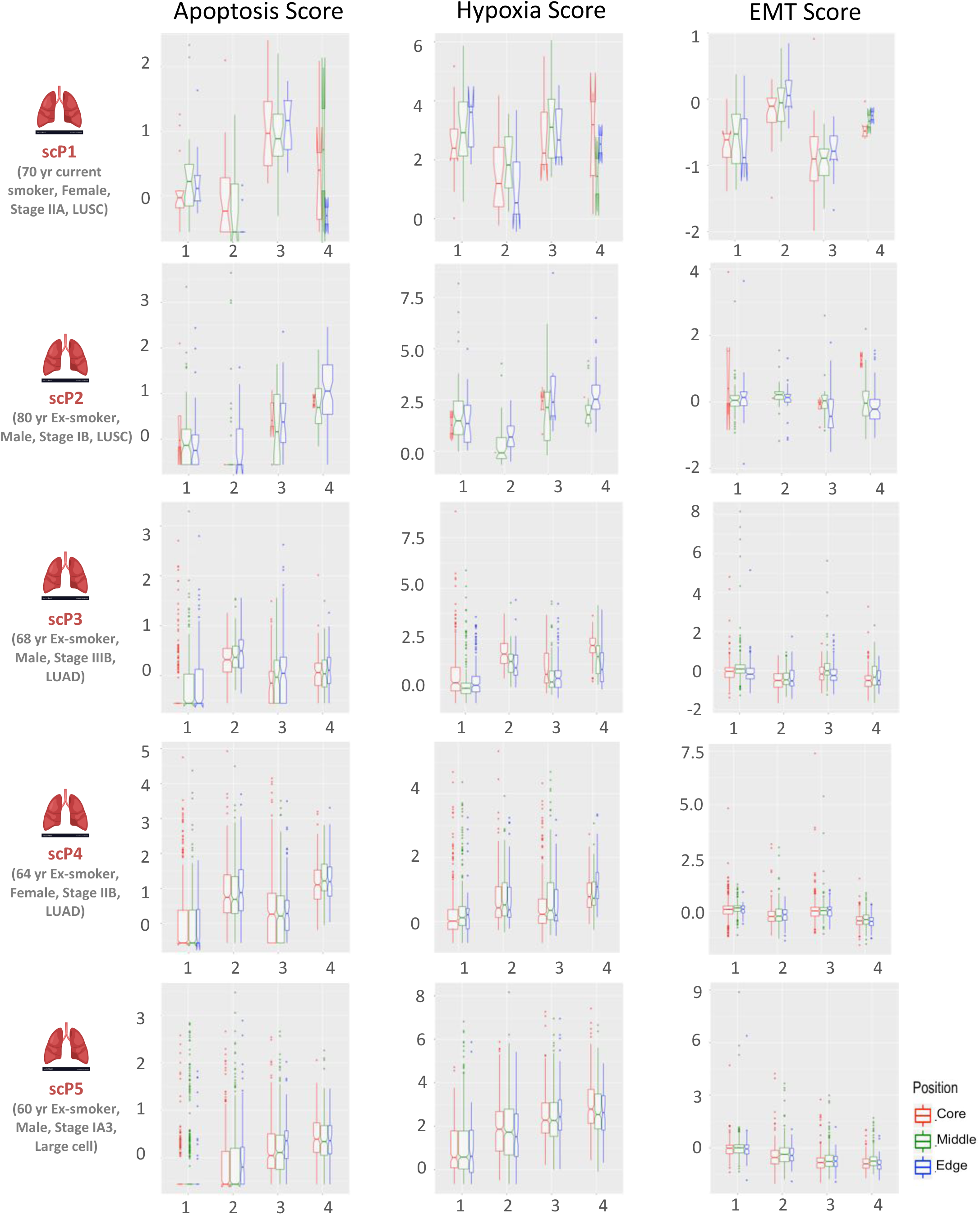
Boxplots showing distribution of apoptosis, hypoxia and EMT scores for each cluster in different regions of tumors. Each dot represents single cell from respective cluster. Scores were calculated from log2CPM data from scRNAseq using same approach as used for bulk RNAseq data (See Methods). Each row represents one patient. X-axis represents different clusters and y-axis represents apoptosis, hypoxia and EMT scores.

## REFERENCES

Aghili, L., Foo, J., DeGregori, J., and De, S. (2014). Patterns of Somatically Acquired Amplifications and Deletions in Apparently Normal Tissues of Ovarian Cancer Patients. Cell Rep. 7, 1310–1319.

Alexandrov, L.B., Nik-Zainal, S., Wedge, D.C., Aparicio, S.A.J.R., Behjati, S., Biankin, A. V, Bignell, G.R., Bolli, N., Borg, A., Børresen-Dale, A.-L., et al. (2013). Signatures of mutational processes in human cancer. Nature 500, 415.

Alexandrov, L.B., Ju, Y.S., Haase, K., Van Loo, P., Martincorena, I., Nik-Zainal, S., Totoki, Y., Fujimoto, A., Nakagawa, H., Shibata, T., et al. (2016). Mutational signatures associated with tobacco smoking in human cancer. Science (80-.). 354, 618–622.

Andor, N., Graham, T.A., Jansen, M., Xia, L.C., Aktipis, C.A., Petritsch, C., Ji, H.P., and Maley, C.C. (2016). Pan-cancer analysis of the extent and consequences of intratumor heterogeneity. Nat Med 22, 105–113.

Ardeshir-Larijani, F., Bhateja, P., Lipka, M.B., Sharma, N., Fu, P., and Dowlati, A. (2018). KMT2D Mutation Is Associated With Poor Prognosis in Non-Small-Cell Lung Cancer. Clin. Lung Cancer 19, e489–e501.

Bjerregaard, A.-M., Nielsen, M., Hadrup, S.R., Szallasi, Z., and Eklund, A.C. (2017). MuPeXI: prediction of neo-epitopes from tumor sequencing data. Cancer Immunol. Immunother. 66, 1123–1130.

Bolotin, D.A., Poslavsky, S., Mitrophanov, I., Shugay, M., Mamedov, I.Z., Putintseva, E. V, and Chudakov, D.M. (2015). MiXCR: software for comprehensive adaptive immunity profiling. Nat. Methods 12, 380–381.

Brock, A., Chang, H., and Huang, S. (2009). Non-genetic heterogeneity — a mutation-independent driving force for the somatic evolution of tumours. Nat. Rev. Genet. 10, 336–342.

Buffa, F.M., Harris, A.L., West, C.M., and Miller, C.J. (2010). Large meta-analysis of multiple cancers reveals a common, compact and highly prognostic hypoxia metagene. Br. J. Cancer 102, 428–435.

Fan, J., Lee, H.-O., Lee, S., Ryu, D.-E., Lee, S., Xue, C., Kim, S.J., Kim, K., Barkas, N., Park, P.J., et al. (2018). Linking transcriptional and genetic tumor heterogeneity through allele analysis of single-cell RNA-seq data. Genome Res. 28, 1217–1227.

Cingolani, P., Platts, A., Wang, L.L., Coon, M., Nguyen, T., Wang, L., Land, S.J., Lu, X., and Ruden, D.M. (2012). A program for annotating and predicting the effects of single nucleotide polymorphisms, SnpEff: SNPs in the genome of Drosophila melanogaster strain w1118; iso-2; iso-3. Fly (Austin). 6, 80–92.

Cristescu, R., Mogg, R., Ayers, M., Albright, A., Murphy, E., Yearley, J., Sher, X., Liu, X.Q., Lu, H., Nebozhyn, M., et al. (2018). Pan-tumor genomic biomarkers for PD-1 checkpoint blockade-based immunotherapy. Science 362, eaar3593.

Dalerba, P., Kalisky, T., Sahoo, D., Rajendran, P.S., Rothenberg, M.E., Leyrat, A.A., Sim, S., Okamoto, J., Johnston, D.M., Qian, D., et al. (2011). Single-cell dissection of transcriptional heterogeneity in human colon tumors. Nat. Biotechnol. 29, 1120–1127.

De, S. (2011). Somatic mosaicism in healthy human tissues. Trends Genet. 27, 217–223.

De, S., and Ganesan, S. (2017). Looking beyond drivers and passengers in cancer genome sequencing data. Ann. Oncol. Off. J. Eur. Soc. Med. Oncol. 28, 938–945.

Ding, Z., Mangino, M., Aviv, A., Spector, T., Durbin, R., and UK10K Consortium, R. (2014). Estimating telomere length from whole genome sequence data. Nucleic Acids Res. 42, e75.

Dobin, A., Davis, C.A., Schlesinger, F., Drenkow, J., Zaleski, C., Jha, S., Batut, P., Chaisson, M., and Gingeras, T.R. (2013). STAR: ultrafast universal RNA-seq aligner. Bioinformatics 29, 15–21.

Gentles, A.J., Newman, A.M., Liu, C.L., Bratman, S. V, Feng, W., Kim, D., Nair, V.S., Xu, Y., Khuong, A., Hoang, C.D., et al. (2015). The prognostic landscape of genes and infiltrating immune cells across human cancers. Nat Med 21, 938–945.

Gibney, G.T., Weiner, L.M., and Atkins, M.B. (2016). Predictive biomarkers for checkpoint inhibitor-based immunotherapy. Lancet. Oncol. 17, e542–e551.

Hills, S.A., and Diffley, J.F.X. (2014). DNA Replication and Oncogene-Induced Replicative Stress. CURBIO 24, R435–R444.

Hou, J., Lambers, M., den Hamer, B., den Bakker, M.A., Hoogsteden, H.C., Grosveld, F., Hegmans, J., Aerts, J., and Philipsen, S. (2012). Expression Profiling-Based Subtyping Identifies Novel Non-small Cell Lung Cancer Subgroups and Implicates Putative Resistance to Pemetrexed Therapy. J. Thorac. Oncol. 7, 105–114.

Jamal-Hanjani, M., Wilson, G.A., McGranahan, N., Birkbak, N.J., Watkins, T.B.K., Veeriah, S., Shafi, S., Johnson, D.H., Mitter, R., Rosenthal, R., et al. (2017). Tracking the Evolution of Non–Small-Cell Lung Cancer. N. Engl. J. Med. 376, 2109–2121.

Jia, Q., Wu, W., Wang, Y., Alexander, P.B., Sun, C., Gong, Z., Cheng, J.-N., Sun, H., Guan, Y., Xia, X., et al. (2018). Local mutational diversity drives intratumoral immune heterogeneity in non-small cell lung cancer. Nat. Commun. 9, 5361.

Jurtz, V., Paul, S., Andreatta, M., Marcatili, P., Peters, B., and Nielsen, M. (2017). NetMHCpan-4.0: Improved Peptide-MHC Class I Interaction Predictions Integrating Eluted Ligand and Peptide Binding Affinity Data. J. Immunol. 199, 3360–3368.

Karaayvaz, M., Cristea, S., Gillespie, S.M., Patel, A.P., Mylvaganam, R., Luo, C.C., Specht, M.C., Bernstein, B.E., Michor, F., and Ellisen, L.W. (2018). Unravelling subclonal heterogeneity and aggressive disease states in TNBC through single-cell RNA-seq. Nat. Commun. 9, 3588.

Kim, W., Ludlow, A.T., Min, J., Robin, J.D., Stadler, G., Mender, I., Lai, T.-P., Zhang, N., Wright, W.E., and Shay, (2016). Regulation of the Human Telomerase Gene TERT by Telomere Position Effect—Over Long Distances (TPE-OLD): Implications for Aging and Cancer. PLOS Biol. 14, e2000016.

Koboldt, D.C., Zhang, Q., Larson, D.E., Shen, D., McLellan, M.D., Lin, L., Miller, C.A., Mardis, E.R., Ding, L., and Wilson, R.K. (2012). VarScan 2: Somatic mutation and copy number alteration discovery in cancer by exome sequencing. Genome Res. 22, 568–576.

Lambrechts, D., Wauters, E., Boeckx, B., Aibar, S., Nittner, D., Burton, O., Bassez, A., Decaluwé, H., Pircher, A., Van den Eynde, K., et al. (2018). Phenotype molding of stromal cells in the lung tumor microenvironment. Nat. Med. 24, 1277–1289.

Li, B., and Dewey, C.N. (2011). RSEM: accurate transcript quantification from RNA-Seq data with or without a reference genome. BMC Bioinformatics 12, 323.

Li, H., and Durbin, R. (2009). Fast and accurate short read alignment with Burrows-Wheeler transform. Bioinformatics 25, 1754–1760.

Li, S., Garrett-Bakelman, F.E., Chung, S.S., Sanders, M.A., Hricik, T., Rapaport, F., Patel, J., Dillon, R., Vijay, P., Brown, A.L., et al. (2016). Distinct evolution and dynamics of epigenetic and genetic heterogeneity in acute myeloid leukemia. Nat. Med. 22, 792–799.

Lukášová, E., Řezáčová, M., Bačíková, A., Šebejová, L., Vávrová, J., and Kozubek, S. (2019). Distinct cellular responses to replication stress leading to apoptosis or senescence. FEBS Open Bio 9, 870–890.

Williams, M.J., Werner, B., Heide, T., Curtis, C., Barnes, C.P., Sottoriva, A., and Graham, T.A. (2018). from bulk sequencing data. 50.

Macheret, M., and Halazonetis, T.D. (2015). DNA Replication Stress as a Hallmark of Cancer. Annu. Rev. Pathol. Mech. Dis. 10, 425–448.

Marusyk, A., Almendro, V., and Polyak, K. (2012). Intra-tumour heterogeneity: a looking glass for cancer? Nat. Rev. Cancer 12, 323–334.

Mazor, T., Pankov, A., Johnson, B.E., Hong, C., Hamilton, E.G., Bell, R.J.A., Smirnov, I.V., Reis, G.F., Phillips, J.J., Barnes, M.J., et al. (2015). DNA Methylation and Somatic Mutations Converge on the Cell Cycle and Define Similar Evolutionary Histories in Brain Tumors. Cancer Cell 28, 307–317.

Mcgranahan, N., and Swanton, C. (2017). Review Clonal Heterogeneity and Tumor Evolution : Past, Present, and the Future. Cell 168, 613–628.

McGranahan, N., and Swanton, C. (2017). Clonal Heterogeneity and Tumor Evolution: Past, Present, and the Future. Cell 168, 613–628.

Park, Y., Lim, S., Nam, J.-W., and Kim, S. (2016). Measuring intratumor heterogeneity by network entropy using RNA-seq data. Sci. Rep. 6, 37767.

Patel, A.P., Tirosh, I., Trombetta, J.J., Shalek, A.K., Gillespie, S.M., Wakimoto, H., Cahill, D.P., Nahed, B. V, Curry, W.T., Martuza, R.L., et al. (2014). Single-cell RNA-seq highlights intratumoral heterogeneity in primary glioblastoma. Science (80-.). 344, 1396–1401.

Ramaker, R.C., Lasseigne, B.N., Hardigan, A.A., Palacio, L., Gunther, D.S., Myers, R.M., and Cooper, S.J. (2017). RNA sequencing-based cell proliferation analysis across 19 cancers identifies a subset of proliferation-informative cancers with a common survival signature. Oncotarget 8, 38668–38681.

Rosenthal, R., McGranahan, N., Herrero, J., Taylor, B.S., and Swanton, C. (2016). DeconstructSigs: delineating mutational processes in single tumors distinguishes DNA repair deficiencies and patterns of carcinoma evolution. Genome Biol. 17, 31.

Rosenthal, R., Cadieux, E.L., Salgado, R., Bakir, M. Al, Moore, D.A., Hiley, C.T., Lund, T., Tanić, M., Reading, J.L., Joshi, K., et al. (2019). Neoantigen-directed immune escape in lung cancer evolution. Nature.

Roth, A., Khattra, J., Yap, D., Wan, A., Laks, E., Biele, J., Ha, G., Aparicio, S., Bouchard-Côté, A., and Shah, S.P. (2014). PyClone: statistical inference of clonal population structure in cancer. Nat. Methods 11, 396–398.

Sharma, A., Jiang, C., and De, S. (2018). Dissecting the sources of gene expression variation in a pan-cancer analysis identifies novel regulatory mutations. Nucleic Acids Res. 46, 4370–4381.

Shen, R., and Seshan, V.E. (2016). FACETS: allele-specific copy number and clonal heterogeneity analysis tool for high-throughput DNA sequencing. Nucleic Acids Res. 44, e131–e131.

Shugay, M., Bagaev, D. V, Turchaninova, M.A., Bolotin, D.A., Britanova, O. V, Putintseva, E. V, Pogorelyy, M. V, Nazarov, V.I., Zvyagin, I. V, Kirgizova, V.I., et al. (2015). VDJtools: Unifying Post-analysis of T Cell Receptor Repertoires. PLoS Comput. Biol. 11, e1004503.

Suda, K., Kim, J., Murakami, I., Rozeboom, L., Shimoji, M., Shimizu, S., Rivard, C.J., Mitsudomi, T., Tan, A.-C., and Hirsch, F.R. (2018). Innate Genetic Evolution of Lung Cancers and Spatial Heterogeneity: Analysis of Treatment-Naïve Lesions. J. Thorac. Oncol. 13, 1496–1507.

Thorsson, V., Gibbs, D.L., Brown, S.D., Wolf, D., Bortone, D.S., Ou Yang, T.-H., Porta-Pardo, E., Gao, G.F., Plaisier, C.L., Eddy, J.A., et al. (2018). The Immune Landscape of Cancer. Immunity 48, 812–830.e14.

Tomasetti, C., Vogelstein, B., and Parmigiani, G. (2013). Half or more of the somatic mutations in cancers of self-renewing tissues originate prior to tumor initiation. Proc Natl Acad Sci U S A 110, 1999–2004.

Wangari-Talbot, J., and Hopper-Borge, E. (2013). Drug Resistance Mechanisms in Non-Small Cell Lung Carcinoma. J. Can. Res. Updates 2, 265–282.

Warren, R.L., Choe, G., Freeman, D.J., Castellarin, M., Munro, S., Moore, R., and Holt, R.A. (2012). Derivation of HLA types from shotgun sequence datasets. Genome Med. 4, 95.

Wilkerson, M.D., Yin, X., Hoadley, K.A., Liu, Y., Hayward, M.C., Cabanski, C.R., Muldrew, K., Miller, C.R., Randell, S.H., Socinski, M.A., et al. (2010). Lung squamous cell carcinoma mRNA expression subtypes are reproducible, clinically important, and correspond to normal cell types. Clin. Cancer Res. 16, 4864–4875.

Wilkerson, M.D., Yin, X., Walter, V., Zhao, N., Cabanski, C.R., Hayward, M.C., Miller, C.R., Socinski, M.A., Parsons, A.M., Thorne, L.B., et al. (2012). Differential Pathogenesis of Lung Adenocarcinoma Subtypes Involving Sequence Mutations, Copy Number, Chromosomal Instability, and Methylation. PLoS One 7, e36530.

Yoshihara, K., Shahmoradgoli, M., Martínez, E., Vegesna, R., Kim, H., Torres-Garcia, W., Treviño, V., Shen, H., Laird, P.W., Levine, D.A., et al. (2013). Inferring tumour purity and stromal and immune cell admixture from expression data. Nat. Commun. 4, 2612.

Zhang, A.W., McPherson, A., Milne, K., Kroeger, D.R., Hamilton, P.T., Miranda, A., Funnell, T., Little, N., de Souza, C.P.E., Laan, S., et al. (2018). Interfaces of Malignant and Immunologic Clonal Dynamics in Ovarian Cancer. Cell 173, 1755–1769.e22.

